# Lanthanide Cathodophores for Multicolor Electron Microscopy

**DOI:** 10.1101/2023.12.11.570835

**Authors:** Sohaib Abdul Rehman, Jeremy B. Conway, Amy Nichols, Edward R. Soucy, Amanda Dee, Kristal Stevens, Simon Merminod, Isabella MacNaughton, Abigail Curtis, Maxim B. Prigozhin

## Abstract

Electron microscopy (EM) and fluorescence imaging are indispensable techniques that provide complementary information on cellular organization. Combining these two modalities is a long-standing challenge in bioimaging. In principle, it should be possible to use the electron beam both for ultrastructural imaging and for molecular localization. The latter could be accomplished by directly exciting suitable biomolecular labels and detecting their luminescence – a process termed cathodoluminescence (CL). Here, we achieve multicolor, single-particle CL imaging of sub-20-nm lanthanide nanocrystals (cathodophores) in the same field of view on the surface of a mammalian cell while simultaneously imaging cellular ultrastructure. In pursuit of this goal, we have developed a comprehensive framework for single-particle CL imaging of lanthanide nanocrystals. By mitigating nonlocal excitation due to secondary electrons, we achieved single-particle detection of multiple spectrally distinct types of sub-20-nm cathodophores. The smallest detectable cathodophores were sub-12 nm in diameter. We found that the CL emission rate scaled linearly with nanocrystal diameter. Furthermore, even in the absence of inert shells, cathodophores were not quenched in the context of mammalian cells processed for EM imaging using heavy-metal staining and sputter-coating. These findings establish cathodophores as promising biomolecular tags for multicolor EM. Moreover, our results inform general design rules for precise control and rational engineering of future generations of single-particle cathodoluminescent nanoprobes.

## Introduction

Precise localization of proteins within the context of cellular ultrastructure, such as the plasma membrane, organelles, and chromatin, is integral to proper protein function and is fundamental to a myriad of biological phenomena. Unfortunately, no single microscopy technique is perfectly suited for imaging both cellular ultrastructure and specific proteins. EM enables ultrastructural imaging at nanoscale spatial resolution, but has relatively low molecular specificity compared to fluorescence imaging. The opposite is generally true of fluorescence microscopy. Thus, EM and fluorescence imaging are orthogonal and complementary in their ability to localize different cellular constituents using distinct contrast mechanisms. Correlative light and electron microscopy techniques combine the benefits of EM and fluorescence imaging^1^. However, such methods are complicated by incompatible sample preparation protocols in fluorescence microscopy and EM^2^. Furthermore, since the EM and fluorescence images are acquired sequentially, CLEM requires challenging nanoscale registration of the two types of images^3^.

Theoretically, it should be possible to use the same scanned electron beam to detect heavy-metal-stained cellular ultrastructure, and to directly excite luminescence in suitable nanoprobes that could serve as multicolor labels for multiplexed molecular detection. This strategy harnesses the light-electron-matter interaction termed cathodoluminescence (CL)^4^. A bioimaging approach based on CL as a contrast mechanism would have two key advantages. First, because free electrons have a much shorter wavelength than photons, both EM and CL imaging would have nanoscale resolution limited by the focused electron beam (∼5 nm), providing a perfect spatial match between the two contrast mechanisms at the molecular scale. Second, CL imaging would have the ability to acquire both ultrastructural (via secondary electron detection) and molecular (via CL) information from the same 2D scan that would not require any additional image correlation or registration (Fig. 1A&B).

**Figure 1:**
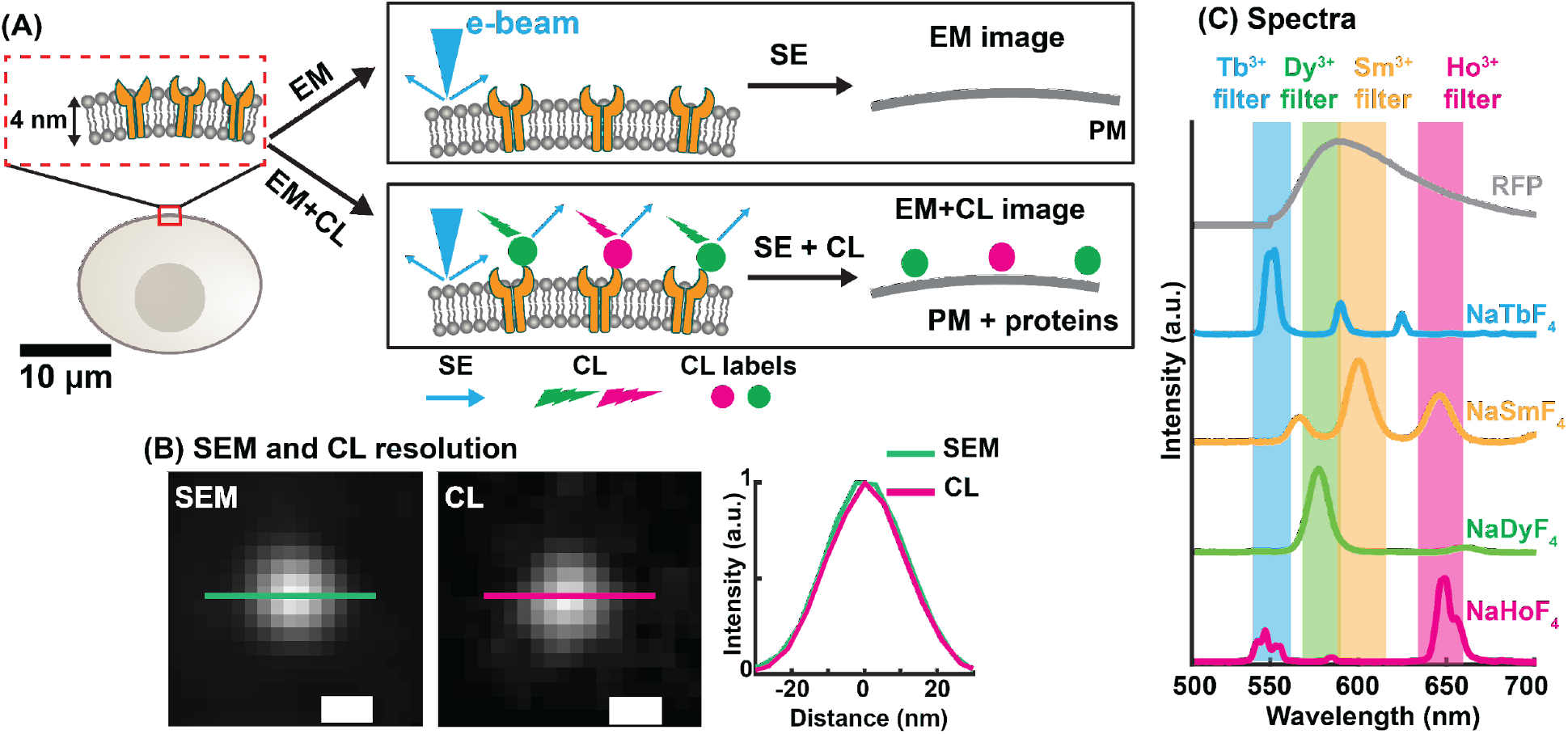
Concept and rationale for multicolor electron microscopy via cathodoluminescence imaging of single cathodophores. **(A)** Schematic showing the plasma membrane (PM) and three transmembrane receptor proteins. In EM, only the heavy-metal-stained membrane is observed. CL imaging of receptors labeled with cathodophores (shown as green and magenta circles) allows localization of cathodophores along with the plasma membrane. **(B)** Scanning electron microscopy (SEM) and CL images of the same cathodoluminescent nanocrystal. Cross-sectional profiles are overlapped on the right for comparison. Identical size of the nanocrystal in both SEM and CL images confirms a match in spatial resolution between the two detection methods. **(C)** Ensemble CL spectra of four distinct cathodophores. Shaded regions represent transmission bands of optical filters that were used in CL imaging. For comparison of spectral linewidths, the spectrum of red fluorescent protein (RFP) is included. **Scale bars:** (B) 20 nm.

Ideal nanoprobes for CL imaging would be brightly luminescent and resistant to electron-induced damage to enable fast and sensitive detection, small (less than 20 nm in diameter, roughly matching the size of an immunoglobulin antibody) to efficiently label and precisely localize proteins, and available in a multicolor palette of spectrally distinct types to permit multiplexed detection of several protein targets in a single experiment. In light of these factors, sodium-fluoride-based lanthanide nanoparticles – inorganic nanocrystals that contain ions from the lanthanide series – are promising candidates given their stability under the electron beam^5^ and their narrow spectral features, which are suitable for multiplexing using different lanthanide dopants (e.g., Ho^3+^, Dy^3+^, Sm^3+^, Tb^3+^)^6^ (Fig. 1C). Unfortunately, CL from multiple, spectrally distinct, sub-20-nm nanocrystals within the same field of view has never been demonstrated. Earlier efforts were focused either on single-color imaging of nanocrystals of various sizes^5,7–10^, or multicolor imaging of large, >100 nm nanoparticles and nanoparticle aggregates^11,12^.

Here, we found that nonlocal excitation of nanocrystals by scattered secondary electrons was the primary limiting factor preventing multicolor CL imaging of molecular-scale nanocrystals. We mitigated nonlocal excitation to achieve multicolor CL imaging of nanocrystals in the same field of view with nanoscale resolution, which is an essential piece of evidence for single-particle CL detection. This advance allowed detection of single lanthanide nanocrystals down to sub-12 nm in diameter. We also synthesized dual-doped nanocrystals to generate a distinct spectral signature, thus increasing the multiplexing capability. In general, spectral profiles of single nanocrystals were consistent with ensemble measurements. We call the nanocrystals *cathodophores* (used interchangeably with *nanocrystals* and *nanoparticles*) by analogy with fluorophores in fluorescence imaging (see SI Figs. 1–4 for TEM images of cathodophores).

Achieving single-particle sensitivity allowed investigating the CL properties of single nanocrystals, thereby establishing several rational design rules for cathodophore engineering. First, cathodophores of the form Na*Ln*F_4_ (where *Ln*^*3+*^ is the lanthanide dopant) displayed bright CL emission and distinct spectra determined by the Ln^3+^ dopant. Second, the CL emission rate scaled linearly with nanocrystal diameter – a favorable scaling for detecting even smaller nanoprobes in the future. Finally, we demonstrated multicolor CL imaging of cathodophores on the surface of a mammalian cell processed for EM, establishing both their biocompatibility and resistance to quenching by heavy-metal staining and sputter-coating. In summary, we developed a comprehensive pipeline for nanoscale, single-particle CL imaging, and used this methodology to establish lanthanide nanocrystals as a promising candidate for multiplexed molecular detection in multicolor EM.

## Results and Discussion

### Nonlocal electron excitation can prohibit single-particle multicolor CL imaging

CL imaging was performed using a scanning electron microscope (SEM) with a custom parabolic mirror and an optical detection system to collect CL (SI Fig. 5). We observed that in samples containing two different types of nanocrystals, NaHoF_4_ and NaDyF_4_ (i.e., doped with Ho^3+^ or Dy^3+^ ions, respectively), excitation of a single nanocrystal (doped with either Ho^3+^ or Dy^3+^ ions, but not both) led to CL signal in both Ho^3+^ and Dy^3+^ color channels (Fig. 2A). In optical microscopy, this result would be highly anomalous. It would suggest that a single fluorophore emits over two distinct spectral ranges. Therefore, we suspected an EM-specific phenomenon: nonlocal excitation of CL in nanocrystals that were present in the field of view of the parabolic mirror but were not actively imaged by the primary electron beam.

**Figure 2:**
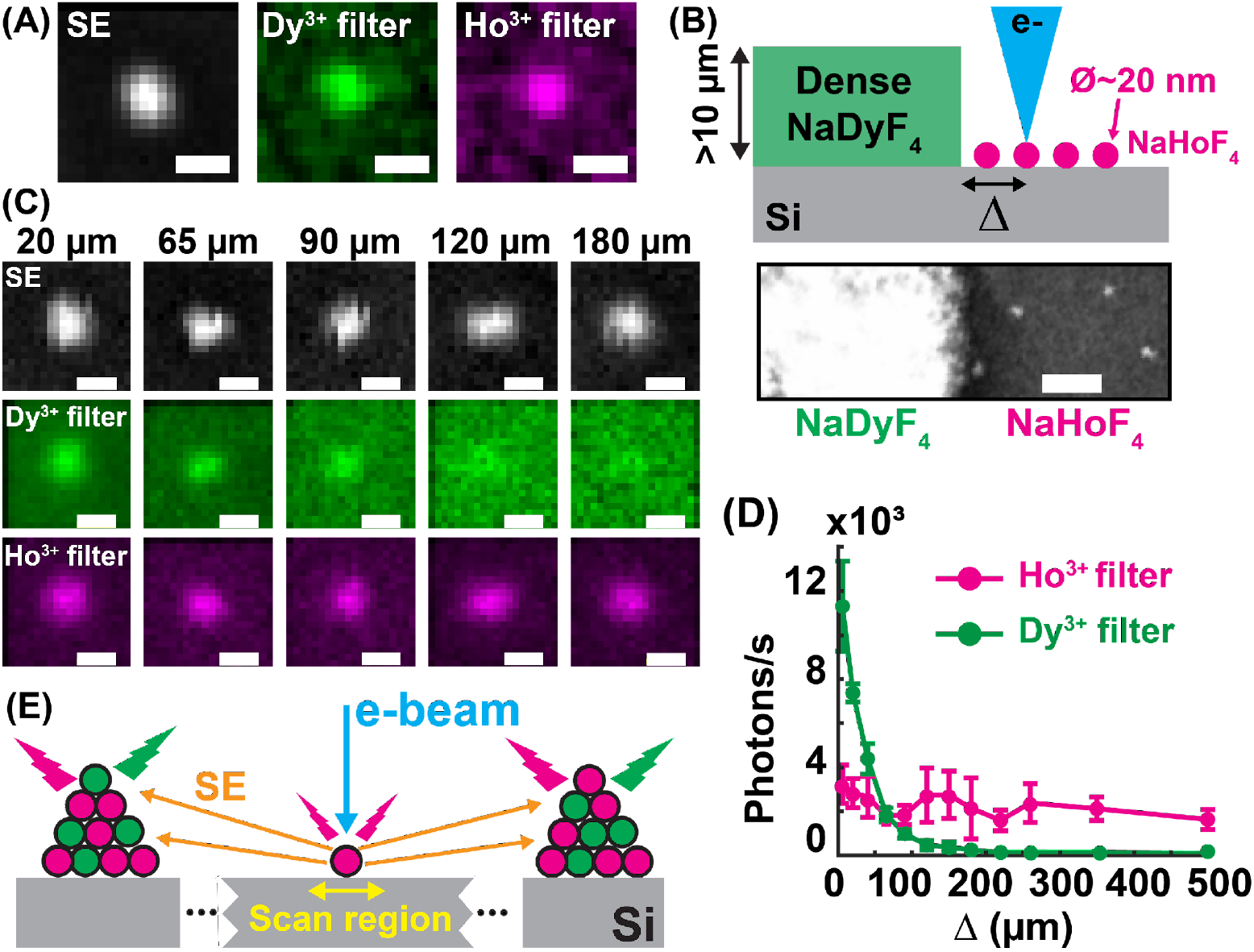
Nonlocal excitation of CL by stray secondary electrons limits multicolor single-particle imaging in the presence of nanoparticle aggregates. **(A)** A single lanthanide nanocrystal imaged within a dense sample containing both NaHoF_4_ and NaDyF_4_ nanocrystals. Anomalous luminescence was observed in both Ho^3+^ and Dy^3+^ color channels. **(B)** A schematic and an SEM image of a sample containing a concentrated edge of NaDyF_4_ nanocrystals on the left and sparsely distributed single NaHoF_4_ nanocrystals on the right. Single NaHoF_4_ nanocrystals were imaged at different distances from the thick NaDyF_4_ layer to characterize the effect of nonlocal excitation. (Top: schematic, not to scale; bottom: SEM image). **(C)** SEM and CL images of sparse NaHoF_4_ cathodophores at different distances from the NaDyF_4_ edge. CL was observed in both Dy^**3+**^ and Ho^**3+**^ color channels when NaHoF_4_ cathodophores were imaged close to the NaDyF_4_ edge due to the nonlocal excitation of NaDyF_4_ nanocrystals by stray secondary electrons. **(D)** Rate of CL detection from single NaHoF_4_ nanocrystals in Ho^3+^ and Dy^3+^ color channels at different distances from the NaDyF_4_ edge. **(E)** An illustration of nonlocal CL signal: The primary electron beam excites a single nanocrystal, which emits both CL and SE. Nearby nanoparticle aggregates are excited by SE, generating aberrant non-local CL signal. **Scale bars:** (A, C) 30 nm, (B) 200 nm.

To determine whether non-local excitation was the source of the crosstalk, we separated cathodophores of two different colors on a Si wafer (total area ∼1 cm^2^). Half of the wafer was covered with a thick layer (>1 μm) of luminescent NaDyF_4_ nanocrystals, while the other half of the wafer was coated with sparsely dispersed individual luminescent NaHoF_4_ nanocrystals (Fig. 2B). When we imaged the individual isolated NaHoF_4_ nanocrystals located >200 µm away from the edge of the dense NaDyF_4_ layer, no CL signal was observed in the Dy^3+^ channel. This result was expected because of the minimal crosstalk between NaHoF_4_ emission and the Dy^3+^ spectral channel (∼3%, Fig. 1C, SI Fig. 6). However, when NaHoF_4_ nanocrystals located closer than 200 µm to the NaDyF_4_ edge were imaged, luminescence was detected in the Dy^3+^ channel in addition to the expected signal in the Ho^3+^ channel (Fig. 2C&D). CL signal in the Dy^3+^ channel increased from <100 photons/s at >100 µm to ∼12,000 photons/s at 5 µm from the NaDyF_4_ edge. We attributed this anomaly to the nonlocal excitation of the dense NaDyF_4_ layer by stray secondary electrons originating from the single NaHoF_4_ nanocrystals actively imaged by the primary electron beam (Fig. 2E).

A typical EM sample of lanthanide nanocrystals deposited on a Si substrate commonly contained both single isolated nanoparticles and nanoparticle aggregates. When a single nanocrystal was imaged within such a sample, it served as a source of secondary electrons, which in turn excited nanoparticle aggregates in the field of view of the parabolic mirror (SI Fig. 7). Because of this nonlocal electron excitation, CL signal could contain a contribution from nanoparticle aggregates located tens of microns away from the nanocrystal that was actively imaged with the primary electron beam. These nonlocal excitations prohibited multicolor single-particle CL imaging. Fig. 2E illustrates how secondary electrons could cause nonlocal excitation of nanoparticle aggregates, leading to unexpected CL in multiple color channels. Interestingly, since the nanocrystals were composed of heavier elements than the Si substrate, they produced more secondary electrons. Consequently, a nanocrystal imaged by the electron beam appeared to emit in multiple color channels, while the background originating from the Si substrate did not.

### Mitigation of nonlocal electron excitation permits multicolor single-particle imaging of spectrally distinct lanthanide nanocrystals

We asked whether nonlocal excitation would prohibit multicolor imaging in samples containing only single nanocrystals, i.e., in the absence of nanoparticle aggregates. Such samples require careful preparation, but are more relevant for future biological applications where single cathodophores tag individual proteins. First, we used Monte Carlo simulations of electron trajectories to examine the excitation crosstalk between two adjacent nanocrystals of 20 nm diameter each (Fig. 3A). The simulations showed that for two neighboring nanocrystals, if one was excited with the electron beam, <2% of the energy was deposited into the neighboring nanocrystal, even if the two nanoparticles were in direct contact. Such minimal crosstalk indicated that multicolor CL imaging with cathodophores should be possible in samples containing individual non-aggregated nanocrystals.

**Figure 3:**
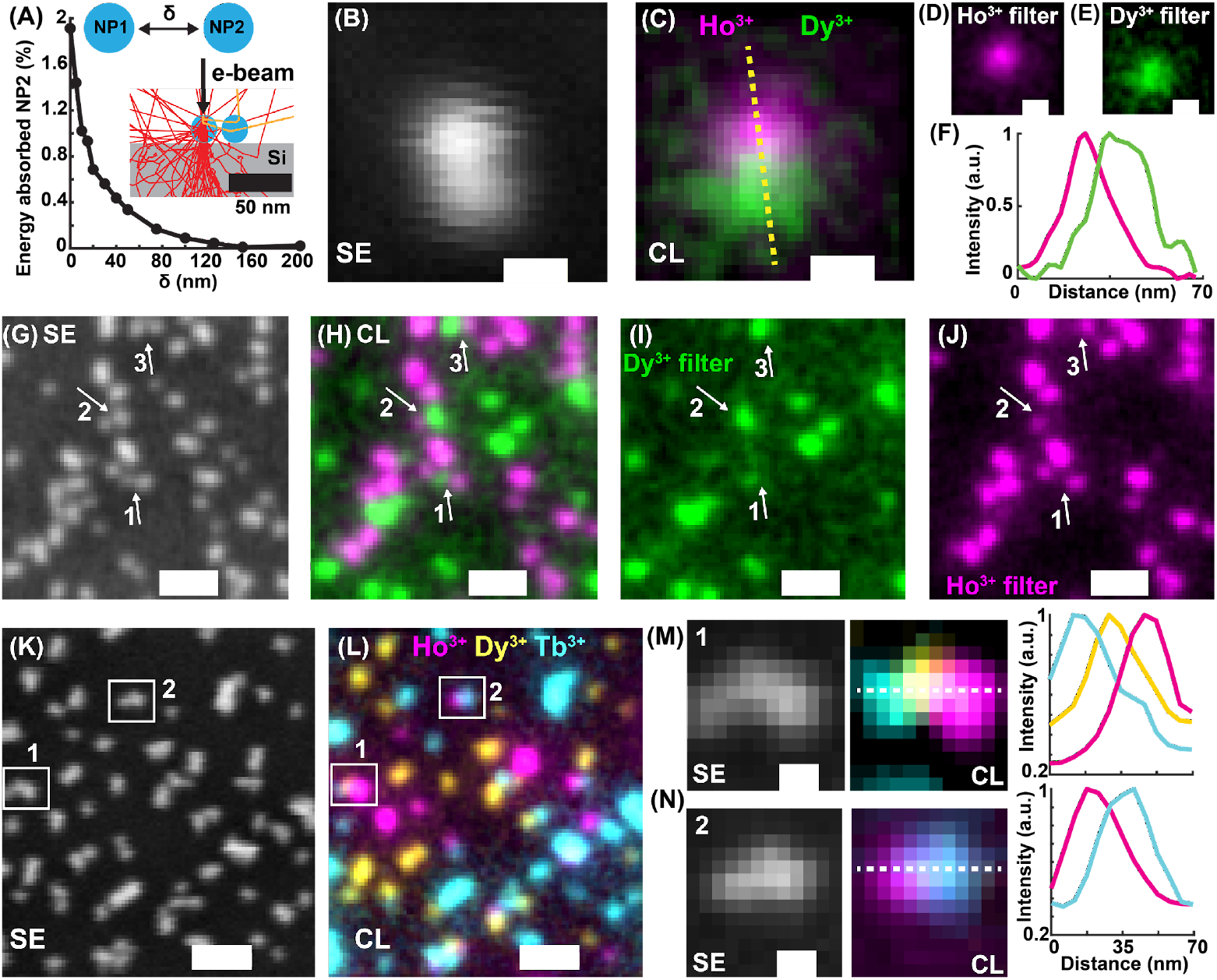
Multicolor single-particle CL imaging of spectrally distinct cathodophores at the nanoscale. **(A)** Excitation crosstalk between two adjacent nanocrystals as analyzed by Monte Carlo simulations. Nanoparticle 1 (NP1) was excited with a 3 keV electron beam and the percentage of energy absorbed by NP2 (normalized to that of NP1) was calculated at different distances, *δ*, from NP1. 5,000 trajectories were simulated at each distance *δ*. **(B)** SE image of two cathodophores located in direct proximity. **(C)** Composite CL image of the cathodophores in (B), obtained by merging signals in **(D)** Ho^3+^ color channel and **(E)** Dy^3+^ color channel. **(F)** Cross-sectional profiles of the CL image along the dotted line shown in (C). **(G)** SE image of a field of view containing NaHoF_4_ and NaDyF_4_ cathodophores. **(H)** Dual-color CL image of the field of view in (G), obtained by merging **(I)** Dy^3+^ color channel, and **(J)** Ho^3+^ color channel. Each nanocrystal was detected in one of the two color channels, even when located close to cathodophores of a different color; see arrows. **(K)** SE image of a field of view containing NaHoF_4_, NaDyF_4_, and NaTbF_4_ cathodophores. **(L)** Composite CL image of the field of view in (K) obtained by merging CL signals from Ho^3+^, Dy^3+^, and Tb^3+^ color channels. **(M, N)** Zoomed-in SE (left) and CL images (right) from regions indicated in (K, L), and CL cross-sectional profiles along the dotted lines in (M) and (N). **Scale bars:** (B–E) 20 nm, (G–L) 100 nm, (M, N) 20 nm.

To test whether this computational result could be confirmed experimentally, we optimized the sample preparation to minimize nanoparticle aggregates, obtaining a 2D distribution of single nanocrystals on the Si wafer (SI section 2). Indeed, using this protocol, we were able to perform multicolor imaging with NaHoF_4_ and NaDyF_4_ nanocrystals in direct proximity (Fig. 3B&C, SI Fig. 8). Images of the nanocrystals in Ho^3+^ and Dy^3+^ color channels (Fig. 3D&E) and their cross-sectional profiles (Fig. 3F) show that CL signal from the two nanocrystals was spatially separated, which allowed distinguishing and classifying them. We also imaged a dense sample of mixed NaHoF_4_ and NaDyF_4_ nanocrystals (Fig. 3G–J, SI Fig. 9) and again were able to observe spectrally distinct luminescence at the single-particle level in the same field of view. Importantly, each nanocrystal appeared in only one of the two CL spectral channels, even when the nanocrystal was located within a few nanometers from other nanocrystals of a different spectral identity (Fig. 3B–F and arrows in Fig. 3G–J). We used a Bayesian approach to assign a color to each nanocrystal. Out of 54 nanocrystals in the image, 23 were classified as NaDyF_4_ and the remaining 31 as NaHoF_4_ (SI Fig. 12, SI section 8). Furthermore, we extended CL imaging to three colors using NaHoF_4_, NaDyF_4_, and NaTbF_4_ nanocrystals (Fig. 3K–N, SI Figs. 10&11). These lanthanide dopants were chosen because of their minimal spectral overlap. Fig. 3K&L show secondary electron and CL images of a field of view containing the three types of cathodophores (see SI Fig. 10 for individual spectral channels). Again, the nanocrystals emitted CL in spectrally distinct channels even when adjacent to nanocrystals of a different color (Fig. 3M&N).

Overcoming nonlocal excitation due to stray secondary electrons was the key development in achieving nanoscale, multicolor CL imaging of single cathodophores in one field of view. Because nonlocal excitation could lead to anomalous apparent luminescence from nanocrystals, multicolor imaging of multiple types of spectrally distinct nanoparticles in the same field of view served as an essential piece of evidence for single-particle CL detection. On the contrary, single-color CL imaging did not by itself establish single-particle CL sensitivity, even when the CL signal appeared to coincide with the image of the nanocrystal in the secondary electron channel. As described in the next section, achieving single-particle CL sensitivity not only permitted multicolor CL imaging, but also enabled accurate single-particle characterization of CL emission from individual nanocrystals of a single spectral identity. In the presence of nonlocal CL signal, such measurements led to erroneous brightness values that had no correlation with nanocrystal size (SI Fig. 13). Importantly, achieving multicolor single-particle imaging by eliminating nanoparticle aggregates also allowed rejecting the hypothesis that anomalous CL signal described in Fig. 2A was caused by excitons diffusing in the Si substrate rather than secondary electrons scattered in the free space.

### Achieving precise control over CL properties by engineering cathodophore architecture

Achieving single-particle sensitivity in CL imaging allowed tuning the spectra and enhancing the brightness of cathodophores by controlling their architecture. The brightness of cathodophores affects not only the speed and sensitivity of CL detection but also the spatial resolution and labeling specificity, as it determines the size of the smallest detectable cathodophores. A key parameter affecting the brightness of cathodophores is the concentration of CL-conferring color centers, i.e., lanthanide ions. To evaluate how the concentration of lanthanide ions affected CL brightness, we co-doped cathodophores with a combination of luminescent lanthanide ions (Ho^3+^or Dy^3+^) and non-luminescent host ions (Gd^3+^) and assessed their brightness at the single particle level. Gd^3+^ions were chosen because they do not emit light in the visible spectrum^13^ but occupy the same crystal lattice positions as the emissive lanthanide ions, thus maintaining the stoichiometry of cathodophores. See Fig. 4A and SI section 10 for detection rate calculations. Fig. 4B&C show CL from single cathodophores as a function of the concentration of Ho^3+^ and Dy^3+^ ions, respectively. The CL increased with an increase in dopant concentration before reaching a plateau. The enhancement in brightness can be attributed to the increase in the number of cathodoluminescent ions. However, at high dopant concentrations, the increase in brightness may be counteracted by quenching due to energy migration and cross-relaxation between dopant ions, similar to upconverting NPs^14^.

**Figure 4:**
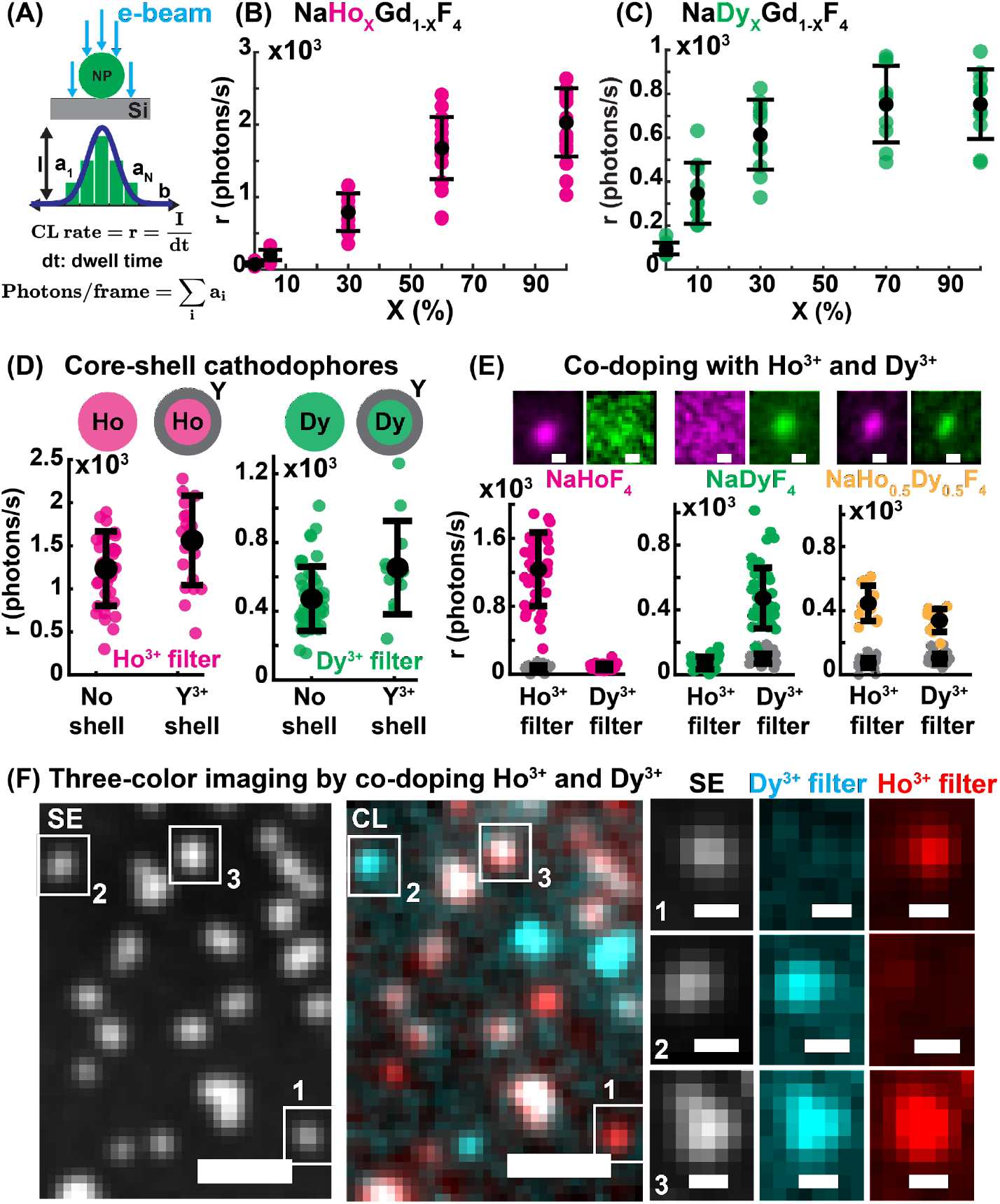
Controlling single-particle CL properties by engineering cathodophore architecture. **(A)** An illustration showing calculations to obtain the rate of CL detection, *r*, and the number of photons in an image frame from a single cathodophore. *a*_*N*_ is the intensity of the *N*^*th*^ pixel in the CL image, *I* is the amplitude of the 2D Gaussian fit of the CL image of a cathodophore, and *dt* is the pixel dwell time. **(B)** CL photon detection rate for different doping concentrations of Ho^3+^ in NaHo_x_Gd_1-x_F_4_ nanocrystals. CL intensity was measured in the Ho^3+^ color channel (cathodophore diameter: 20–25 nm). **(C)** CL photon detection rate for different concentrations of Dy^3+^ in NaDy_x_Gd_1-x_F_4_ nanocrystals. CL intensity was measured in the Dy^3+^ color channel (cathodophore diameter: 20–25 nm). **(D)** CL photon detection rate from cathodophores containing an inert NaYF_4_ shell, compared to the detection rate from cathodophores with no shell (cathodophore diameter: 18–23 nm in both cases). CL signal from cathodophores with Ho^3+^ and Dy^3+^ cores was measured in Ho^3+^ and Dy^3+^ color channels, respectively. **(E)** CL photon detection rate of cathodophores containing both Ho^3+^ and Dy^3+^ ions, i.e., NaHo_0.5_Dy_0.5_F_4_, in Ho^3+^ and Dy^3+^ color channels (cathodophore diameter: 18–23 nm). Photon detection rates for NaHoF_4_ and NaDyF_4_ nanocrystals of 18–23 nm diameter are shown for comparison. Gray dots show background CL from non-luminescent NaGdF_4_ nanocrystals in the respective color channels. **(F)** Three-color imaging with NaHoF_4_ (red), NaDyF_4_ (cyan), and NaHo_0.5_Dy_0.5_F_4_ (white) cathodophores. **Scale bars:** (E) 20 nm, (F: SE and CL images) 100 nm, (F: zoomed-in images) 20 nm. Error bars in (B–E) show mean and standard deviation of the data.

Additionally, surface quenching can significantly reduce the brightness of upconverting nanocrystals due to nonradiative energy transfer to surface defects and the solvent^15^. Indeed, epitaxial growth of optically inert shells on top of active luminescent cores can enhance the brightness of upconverting nanocrystals by up to two orders of magnitude^15^. To test whether cathodophores were getting quenched due to non-radiative losses at the nanoparticle surface, we epitaxially grew optically inert NaYF_4_ shells of ∼4 nm thickness on top of luminescent NaHoF_4_ and NaDyF_4_ cores and contrasted the CL response of these core-shell cathodophores with their core-only counterparts. We found that for core-only and core-shell cathodophores of 18–23 nm total diameter the detected CL rate was nearly identical (Fig. 4D). Although inert shells did not increase cathodophore brightness on the per-volume basis, accounting for the total number of emissive lanthanide ions in each type of nanocrystal led us to conclude that each emissive lanthanide ion in core-shell nanoparticles was ∼2–3 times brighter than in core-only nanocrystals. This increase in brightness could be due to either the passivating effect of the inert shell, or an increased overlap between the electron interaction volume and the optically active core in core-shell versus core-only nanocrystals. Regardless, because the larger diameters of core-shell nanocrystals offset their increased per-ion brightness, they did not provide a net gain in CL intensity per nanocrystal size. Hence, core-only nanocrystals were used for all subsequent measurements. The relative insensitivity of CL brightness to inert shells was likely at least in part due to the apparent saturation of cathodophore luminescence at the explored values of electron beam current (SI Fig. 17).

Next, we asked whether multiple lanthanide ions with non-overlapping emission spectra could be co-doped into a single nanocrystal to generate cathodophores with distinct spectral signatures. If available, such permutationally co-doped cathodophores could enable multiplexed protein detection using fewer spectral detection channels. Fig. 4E shows CL emission from cathodophores co-doped with Ho^3+^ and Dy^3+^ ions in equal proportions, resulting in the NaHo_0.5_Dy_0.5_F_4_ nominal chemical formula. The CL signal from these cathodophores was visible in both Ho^3+^ and Dy^3+^ color channels, while their singly doped counterparts exhibited luminescence in only one of the two channels. The CL signal from NaHo_0.5_Dy_0.5_F_4_ was dimmer than NaHoF_4_ and NaDyF_4_ in their respective emission channels. This reduced brightness could be because dual-doped nanocrystals contained fewer Ho^3+^ and Dy^3+^ ions than each of the singly doped cathodophores. Moreover, cross-relaxation pathways between the two types of ions, i.e., Ho^3+^ and Dy^3+^, could also result in a loss of CL signal^16^. Despite this reduced brightness, sub-20-nm NaHo_0.5_Dy_0.5_F_4_ cathodophores were detected in both Ho^3+^ and Dy^3+^ spectral channels (Fig. 4E). We also successfully performed three-color imaging using NaHo_0.5_Dy_0.5_F_4_ as an additional color along with NaHoF_4_ and NaDyF_4_ cathodophores (Fig. 4F). These results demonstrate the utility of co-doping multiple lanthanide ions into a single nanocrystal to obtain spectrally distinguishable cathodophores, thus increasing multiplexing capability.

### Characterization of single-particle CL emission in lanthanide cathodophores

Since single cathodophores are visible in the secondary electron channel, cathodophore localization can be done with secondary electrons, which have a much higher quantum yield than CL, thus allowing faster imaging. CL would then be used solely for spectral color assignment. This decoupling of cathodophore brightness from localization precision differentiates CL imaging from single-molecule localization microscopy and permits simultaneous nanoscale imaging and spectral identification with only a few tens of photons detected above the background for a single cathodophore. We, therefore, wished to determine the minimum cathodophore size that would still provide enough photons for color assignment. Small cathodophores are desirable because they would lead to better spatial resolution and, due to reduced steric effects, would permit efficient cell and tissue penetration and biomolecular targeting.

To understand how the CL emission rate scales with nanocrystal size and to determine the size limit of detection, we imaged single cathodophores of different sizes. For all dopants, cathodophore brightness scaled linearly with diameter (Fig. 5A). This linear relationship suggested that the brightness of a cathodophore was primarily determined by its axial dimension. This observation was consistent with Monte Carlo simulations of electron trajectories, which revealed that the total energy absorbed by a cathodophore excited with a 3 keV electron beam was proportional to the diameter of the cathodophore (SI Fig. 21). Specifically, this linear relationship arose due to the cylindrical shape of the interaction volume between the 3 keV electron beam and the nanocrystals in the 15–30 nm diameter range (SI Fig. 21). Ideally, the entire volume of a cathodophore would be excited by the electron beam, which would lead to a cubic relationship between cathodophore brightness and its diameter. However, matching the electron interaction volume to the volume of a 20-nm-diameter cathodophore would require using beam energies that were too low for our experimental setup (∼1 keV, see SI Fig. 19). Thus, we note that cathodophores were not excited optimally, suggesting opportunities for optimization.

**Figure 5:**
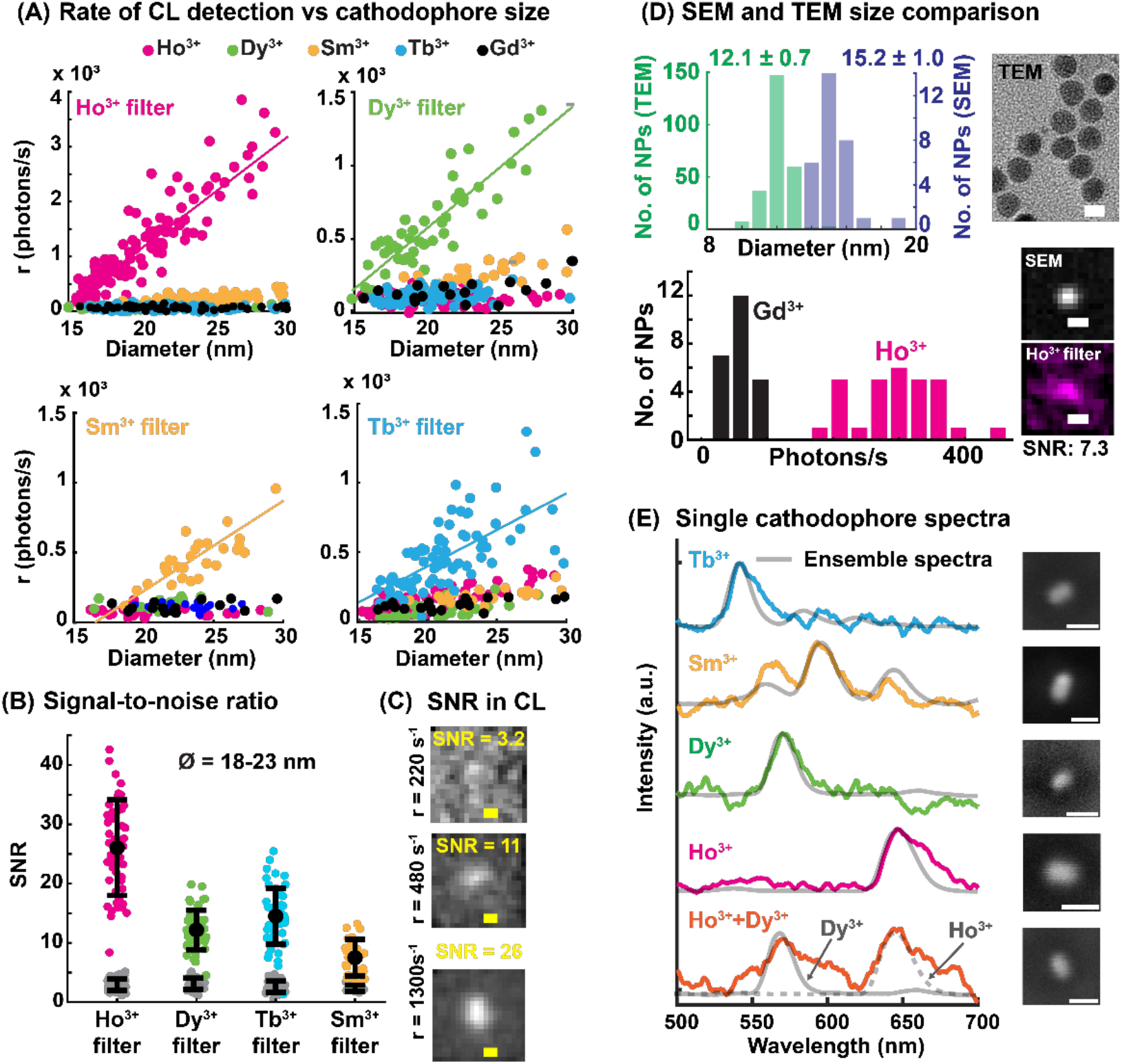
Characterization of single-cathodophore CL emission. **(A)** CL photon detection rate as a function of diameter (full width at half maximum of SE images) for NaHoF_4_, NaDyF_4_, NaSmF_4_, and NaTbF_4_ cathodophores across the spectral filters matched to their primary emission peaks. Linear fits to the CL signal as a function of diameter are also shown. **(B)** SNR of CL images for NaHoF_4_, NaDyF_4_, NaSmF_4_, and NaTbF_4_ cathodophores when imaged through their respective spectral filters. Data is for cathodophores of 18–23 nm diameter. SNR from non-emitting control NaGdF_4_ nanocrystals is shown as gray dots. **(C)** Representative CL images with different SNRs. **(D)** Top: Comparison of the cathodophore size calculated using TEM (diameter of the circle fitted to cathodophores) and SEM images (FWHM of the 2D Gaussian function fitted to the cathodophores). A TEM image of cathodophores is also shown. Bottom: Comparison of the CL detection rate from control (Gd^3+^) and Ho^3+^ cathodophores shown in the top row. Representative SEM and CL images of a cathodophore are also shown. **(E)** CL Spectra of single cathodophores (SEM images on the right) and the corresponding ensemble spectra (gray). **Scale bars:** (C) 20 nm, (D) TEM: 10 nm, SEM: 20 nm, (E) 50 nm. Error bars in (B) show the mean and standard deviation of the data.

Fig. 5A also shows that the brightness (detected photons/s) of cathodophores depended on the type of lanthanide ion, which was likely due to a combination of variations in the energy concentrated in the primary emission peak (e.g., 59% for Ho^3+^ and 40% for Sm^3+^, SI Fig. 6), differing levels of electrobleaching, different absorption cross-sections, and different quantum yields of lanthanide ions. The size of the smallest detectable cathodophore for a given dopant was limited by the background CL from the Si substrate, which followed Poisson statistics (SI Fig. 22–26). However, irrespective of the dopant, sub-20-nm cathodophores were detected and classified (Fig. 5B).

**Figure 6:**
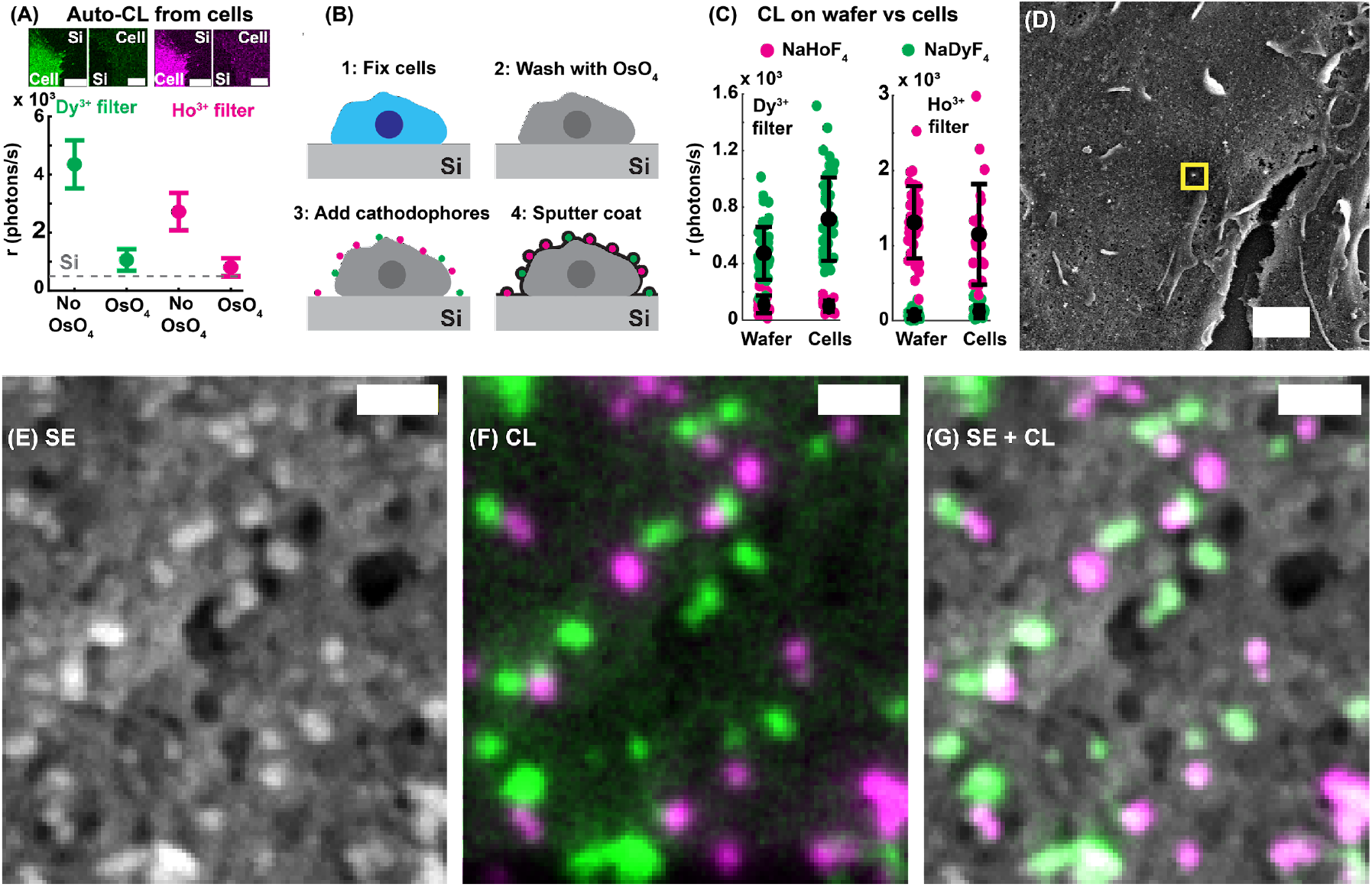
Biocompatible multicolor single-particle CL imaging using cathodophores. **(A)** Auto-CL photon detection rates from cells with and without OsO_4_ treatment in Ho^3+^ and Dy^3+^ color channels. CL background from the Si substrate is also shown for comparison (dashed line). **(B)** An illustration showing the sample preparation procedure for imaging cathodophores on cells. HEK293T cells were fixed, treated with OsO_4_, and dried with HMDS (HMDS drying is not shown in the illustration). Cathodophores were then drop-cast onto the cells. The biological sample with cathodophores was then sputter-coated with a Pt/Pd mixture for SEM imaging. **(C)** CL detection rate from single, 18– 23-nm-diameter NaHoF_4_ and NaDyF_4_ cathodophores on cells prepared for SEM imaging. Samples were prepared according to panel (A). **(D)** SE image of a cell prepared according to (A) with NaHoF_4_ and NaDyF_4_ cathodophores on the surface. **(E, F)** Zoomed-in SE (E) and CL (F) images of the region marked with a yellow square in (D). **(G)** Merged image of the SE and CL channels. **Scale bars:** (A) 10 µm, (D) 1 µm, (E–G) 100 nm. Error bars in (A, C) correspond to mean and standard deviation of the data.

Cathodophore diameters were calculated from secondary electron images, but due to their dependence on the electron beam size, these calculations could be overestimating the actual diameters. To exclude the effect of SEM resolution on the perceived size of the smallest detectable cathodophores, we synthesized uniformly shaped cathodophores of 12.1 ± 0.7 nm diameter (as determined by TEM), which were observed to be 15.2 ± 1.0 nm on our modified SEM (Fig. 5D). NaHoF_4_ cathodophores were co-doped with 20% Lu to achieve this uniform shape. Importantly, these cathodophores were consistently detected in CL images, suggesting that we could detect cathodophores down to at least 11 nm. Furthermore, the linear dependence of CL emission with nanocrystal size discussed above is favorable for further reducing the minimum size of detected nanocrystals. For example, under linear scaling, reducing the nanocrystal size by a factor of two would only reduce its signal proportionally, while under volumetric scaling, the signal would drop by a factor of eight.

To harness cathodophores as bioimaging probes in EM, cathodophores must demonstrate resistance to electron-beam-induced damage and single-particle spectral stability. Indeed, we observed spectral stability of single cathodophores, i.e., consistent spectral profiles between single-particle and ensemble measurements (Fig. 5E, SI Fig. 27–30), which was crucial to match the optical filters to the main emission peaks of cathodophores and thereby minimize the impact of broadband CL from the Si substrate.

### Biocompatible multicolor single-particle CL imaging of cathodophores

Having characterized the emission properties of optimally designed cathodophores on a Si substrate, we next tested their compatibility for bioimaging. Specifically, we sought to achieve multicolor detection of single cathodophores on the surface of a cultured mammalian cell. Here, a critical limitation was the auto-cathodoluminescence (auto-CL) from cells, which, by analogy with autofluorescence in fluorescence imaging, is the intrinsic luminescence of unlabeled cells excited with the electron beam^5,17^. Fixed HEK293T cells exhibited broadband auto-CL emission (SI Fig. 31) and resulted in an average of ∼4,300 and ∼2,700 photons/s in Dy^3+^ and Ho^3+^ color channels, respectively (Fig. 6A). This auto-CL not only reduced the SNR of cathodophores, but also resulted in unexpected CL signal in multiple color channels due to nonlocal excitation of auto-CL when imaging the cathodophores – a similar effect to the one presented in Fig. 2. We found that osmium tetroxide (OsO_4_), a standard fixative and staining agent in EM sample preparation, overcame the auto-CL of cells (Fig. 6A). The auto-CL from OsO_4_-treated cells was comparable to the background CL from the Si substrate, indicating that OsO_4_ treatment was an effective chemical method for background auto-CL suppression.

We then measured CL from cathodophores deposited on the surface of HEK293T cells prepared for standard SEM imaging (see Fig. 6B, SI section 18). Surprisingly, EM sample preparation, including OsO_4_ treatment that was used to mitigate auto-CL, and sputter-coating with a 2.5-nm-thick 80:20 Pt:Pd mixture to avoid sample charging, did not quench CL from NaHoF_4_ and NaDyF_4_ cathodophores (Fig. 6C). CL signal from the cathodophores on cells matched the CL signal of cathodophores on the Si substrate (Fig. 6C). Importantly, sharp spectral features of cathodophores, conducive to efficient spectral filtering and auto-CL background suppression, allowed performing high-SNR single-particle imaging even in a cellular setting. Similar to imaging with the Si substrate, we could detect and classify sub-20-nm cathodophores on the surface of mammalian cells. This result was critical in establishing the stability of cathodophore luminescence in a biological context.

Finally, we successfully performed multicolor imaging of NaHoF_4_ and NaDyF_4_ cathodophores on the surface of HEK293T cells (Fig. 6D–G). Each cathodophore emitted CL in only one spectral channel, making it readily distinguishable. Both the nanoscale location and the spectral identity of each cathodophore, as well as the cellular ultrastructure, were visualized simultaneously. These results demonstrate a proof of concept for multicolor, single-particle CL imaging on the surface of a mammalian cell, with simultaneous visualization of cellular topography. Given the susceptibility of fluorescent dyes to quenching by EM sample preparation protocols^2^, these findings highlight the remarkable photophysical stability of lanthanide nanocrystals. Collectively, these results establish cathodophores as promising candidates that are uniquely suitable for biomolecular labeling in multicolor EM.

## Conclusion and Future Directions

Cathodoluminescence is an attractive contrast mechanism for cellular imaging because it would permit simultaneous ultrastructural imaging and single-particle molecular localization at the nanoscale without the need for spatial registration of images from disparate microscopy techniques. Here, we demonstrate single-particle CL imaging using lanthanide nanocrystals (cathodophores). Cathodophores were well suited for multicolor imaging because of their narrow emission peaks (10–20 nm spectral width) characteristic of the 4f–4f transitions in lanthanide ions^6^. The ionic nature of lanthanide emission also made cathodophores relatively stable to electron beam irradiation. Another benefit of using cathodophores for CL imaging was their strong contrast in secondary electron images, which allowed precise localization without photon detection; CL color channels were used solely for spectral classification. For this reason, the required number of detected photons was much lower when compared to single-molecule localization microscopy, which relies on photon detection for localization^18^.

Overcoming nonlocal excitation by stray secondary electrons was the key advance enabling single-cathodophore measurements, allowing us to develop a comprehensive pipeline for single-particle CL imaging. We showed, for the first time to our knowledge, imaging of multiple cathodophores of different colors in the same field of view. The ability to image single cathodophores allowed studying CL properties of lanthanide nanocrystals, developing optimal imaging parameters, tuning the nanocrystal architecture to achieve enhanced brightness, and generating cathodophores with unique spectral signatures via co-doping. The CL emission rate of single cathodophores increased monotonically as a function of lanthanide ion concentration and was proportional to the nanocrystal diameter. These results suggest general design rules for the precise engineering of cathodophore luminescence. The smallest detectable cathodophores were sub-12 nm in diameter, making them comparable in size to immunoglobulin antibodies^19^. We also showed the biocompatibility of cathodophores by imaging them on the surface of heavy-metal-stained and sputter-coated mammalian cells. CL of cathodophores was spectrally stable and resistant to quenching in this biological context. These findings establish cathodophores as attractive protein labels for future applications in multicolor EM, which would enable visualization of multiple targets against the backdrop of cellular ultrastructure.

Many improvements to the state of the art in cathodophore imaging can be immediately envisioned. For example, to further enhance the multiplexing capability, cathodophores of additional colors can be obtained by exploring the remaining lanthanide ions, which would result in up to nine distinguishable colors^5^. This number could be significantly expanded by permutationally co-doping individual cathodophores with multiple lanthanide ions of different colors. Moreover, the SNR of cathodophores could be increased by better matching the electron excitation volume with the nanocrystal dimensions, designing new cathodophore architectures for increased brightness, and reducing CL background from the biological sample. Methodological developments that would enable these advances include exploring lifetime imaging, reducing the electron beam landing energy, and reducing the working distance in EM imaging. These strategies would further decrease the size of the smallest detectable cathodophores, which matches the size of immunoglobulin antibodies.

In the future, cathodophores can be used to understand the fascinating cell biology of membrane-associated structures that play a role in a vast number of cellular processes, including transmembrane signaling, endocytosis and exocytosis, cell-cell contact formation, autophagy, and cell division. The natural next step in the application of cathodophores to biological imaging would be their functionalization and targeting to label biomolecules of interest, as well as studies of their penetration into cells and tissues. To this end, existing approaches for functionalization and targeting of upconverting nanoparticles can be leveraged^20,21^. This, combined with 3D EM sample preparation techniques such as resin embedding and either ultrathin sectioning^22,23^ or focused ion beam milling^24^, would enable simultaneous visualization of cellular ultrastructure and proteins over entire cells and tissues. Integration of CL imaging with transmission EM, including cryo-EM and cryo-electron tomography, may also be achievable. Lastly, the findings of this work may encourage exploration and systematic characterization of alternative CL-active biolabels, including not only other types of nanoprobes, but also small-molecule cathodophores.

## Materials and Methods

### Nanocrystal synthesis

NaHoF_4_, NaDyF_4_, NaTbF_4_, NaSmF_4_, and NaGdF_4_ nanocrystals were synthesized by the coprecipitation method based on previously reported protocols^5,15,25,26^. Briefly, the appropriate lanthanide chloride hydrate was mixed with oleic acid and 1-octadecene. The reaction was placed under vacuum and the temperature was set to 160 °C for 30 min. Next, the solution was cooled to <30 °C. Sodium hydroxide and ammonium fluoride were combined in methanol, then added to the lanthanide oleate solution. The reaction was mixed for 60 min at room temperature. The temperature was increased to 70–80 °C and maintained for 30 min. Then, for the nanocrystal growth step, the reaction was placed under an argon atmosphere and heated rapidly to 320 °C. The reaction temperature was maintained for 60 min before cooling to <30 °C. For dual-doped nanocrystals (e.g., NaHo_0.5_Dy_0.5_F_4_) and partially doped nanocrystals (e.g., NaHo_0.3_Gd_0.7_F_4_), synthesis was performed as described above, except the total molar quantity of lanthanide chloride hydrates was divided into two quantities based on the desired molar ratio of the two lanthanides. For core-shell nanocrystals, cores were first synthesized as described above then washed with ethanol. Next, shells were synthesized via a second reaction as described above, except the washed core nanocrystals were added to the reaction during addition of ammonium fluoride and sodium hydroxide. Nanocrystals were stored as-synthesized in oleic acid and 1-octadecene at room temperature. Details provided in SI Section 1.

### Nanocrystal sample preparation

As-synthesized nanocrystals were washed by mixing them with ethanol and centrifuging. The resulting pellet was resuspended in n-hexane. This ethanol wash step was performed five times. Finally, the n-hexane-suspended nanocrystals were left undisturbed overnight. Nanocrystals were then pipetted from the top of the settled solution. The nanocrystal solution was drop-cast onto a TEM grid or a plasma-cleaned p-type Si wafer for characterization by TEM or SEM, respectively. Details provided in SI Section.

### CL imaging setup

Imaging was performed on a custom-modified ZEISS SUPRA 55VP FESEM, integrated with a CL imaging system. CL was collected by an off-axis, aluminum parabolic mirror with a collection solid angle of ∼1.34π steradian. The mirror had a focal length of 1 mm. The mirror directed CL out of the vacuum chamber of the SEM, where it was spectrally separated using dichroic mirrors, filtered by band-pass filters, and projected onto photomultiplier tubes (Hamamatsu, H7421-40) using 30-mm-focal-length lenses. CL was simultaneously collected over three color channels. The spectral filters were Chroma ET645/30X (Ho^3+^ color channel), Chroma FF03-575/25 (Dy^3+^ color channel), Semrock FF01-598/25 (Sm^3+^ color channel) and Chroma ET550/20X (Tb^3+^ color channel). Details provided in SI Section 3.

### Single-particle spectral measurements

For single-particle spectra, a cathodophore was repeatedly scanned with the electron beam. CL was collected by the parabolic mirror and focused on an sCMOS camera (Hamamatsu, Orca-Fusion BT) using a 100 mm focal length lens. A diffraction grating (Thorlabs, GT13-03) was placed before the camera to spectrally separate the CL signal. Spectra were obtained with a beam energy of 3 keV and a beam current of 200 pA. The spectral information resided in the first diffraction order of the grating. Spectra were obtained by plotting the intensity of pixels in the first diffraction order as a function of distance from the center of the zeroth order. Spectra presented in Fig. 5E were obtained using this method. Details provided in SI Sections 3 and 16.

### Ensemble spectral measurements

For ensemble spectra, a dense sample of cathodophores was prepared. A 400 x 400 nm region of the sample was repeatedly scanned with the electron beam. CL was collected by the parabolic mirror and focused on the multimode fiber of a commercial spectrometer (Thorlabs, CCS200). Spectra were obtained with a beam energy of 3 keV and a beam current of 200 pA. Spectra presented in Fig. 1C were obtained using this method. Details provided in SI Section 3.

### CL image acquisition

Single-particle CL imaging was performed with an electron beam energy of 3 keV and a beam current of 160–200 pA. The working distance was 6–7 mm. Both the Zeiss SmartSEM software and custom LabVIEW software were used for image acquisition. A region of interest (∼20–100 pixels x 20–100 pixels) was captured with a pixel size of 4–8 nm. Four images were captured during each scan of the electron beam: one secondary electron (SE) image and three CL images in different spectral channels. CL images were smoothed by convolution with a 2D Gaussian function of 0.7 pixels standard deviation. To characterize emission properties of nanocrystals, 50 frames were captured with a dwell time of 1 ms each, resulting in an effective beam dwell time of 50 ms/pixel. These conditions translate to a current density of 3–12 pA/nm^2^ and a total electron dose of ∼62.5 x 10^6^ electrons/pixel. For multicolor experiments, images were acquired with a beam dwell time of 0.1–1 ms and an effective beam dwell time of 30–50 ms. Details provided in SI Section 4.

### Drift correction

Drift correction was performed by acquiring multiple sequential frames using a short dwell time, instead of a single image with a long dwell time. Cathodophores themselves were used as fiducial markers in SE images. The position of a cathodophore was determined by fitting a 2D Gaussian function to its SE image. Euclidean distance between the positions of the cathodophore in successive SE frames was calculated and rounded to the nearest integer. This distance was then used to translate SE and CL images to account for the drift during image acquisition. Details provided in SI Section 4.

### CL emission rate and photons/frame

CL properties of a cathodophore were determined by first localizing it in the SE images and then determining its emission properties at the corresponding location in the CL images. The position of the cathodophore in an SE image was determined by fitting a 2D Gaussian function. The CL image was then fit to a 2D Gaussian function of the form:

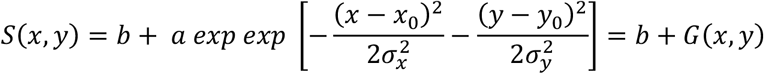

where (*x*_0_, *y*_0_) is the center position of the cathodophore, *b* is the background, *a* is amplitude, and *σ*_*x*_ and *σ*_*y*_ are the standard deviations of the 2D Gaussian function. *G*(*x, y*) is the 2D Gaussian fit to the cathodophore without the background. Importantly, in this equation, *x*0, *y*0, *σ*_*x*_, and *σ*_*y*_ were constrained by the 2D Gaussian fit of the SE image. The rate of CL emission was determined as 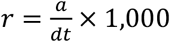, and the total number of photons emitted by the cathodophore in a frame was calculated by summing over *G*(*x, y*).

### Signal-to-noise ratio (SNR) analysis

SNR of a cathodophore in a CL image was determined using the equation *S*/*N*, where *S* corresponds to the summation over *G*(*x, y*), i.e., *S* = ∑*G*(*x, y*). Only pixels within a 2*σ* radius from the center of the Gaussian fit were considered. The noise, *N_i_*, for pixel (*x, y*) was calculated as 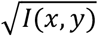 where *I*(*x, y*) is the CL intensity of that pixel. To determine the overall noise, *N*, the per-pixel noise, *N*_*i*_, was added in quadrature over the pixels within a 2*σ* radius from the center of the cathodophore as 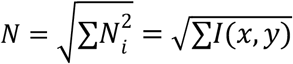. Hence, the overall SNR for a single cathodophore was given by 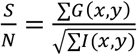.

### Size of the cathodophore

The diameter (also referred to as size) of a cathodophore was determined from the oversampled SE images taken with the ZEISS SmartSEM software (pixel size 0.2–0.6 nm). To determine the diameter of a cathodophore, a 2D Gaussian function was fit to its SE image. Diameter was defined as the full width at half maximum (FWHM) of this fit. FWHM was calculated from the standard deviation of the Gaussian fit as 2.36*σ*. Here *σ* is the average of the two standard deviations (*σ*_*x*_ and *σ*_*y*_) of the 2D Gaussian fit.

### Electrobleaching analysis

To analyze electrobleaching, CL from cathodophores was measured at successive time points. A total of 50 images was captured, each with a pixel dwell time of 1 ms. Electrobleaching was evaluated by monitoring the change in CL signal in binned 5 ms intervals (i.e., 5 frames in each bin). Details provided in SI Section 17.

### Biological sample preparation

Cultured cells were prepared for CL imaging via osmium tetroxide treatment and hexamethyldisilazane (HMDS) drying, based on previously reported protocols^27,28^. Briefly, HEK293T cells were seeded on plasma-cleaned Si wafers, washed with PBS, and fixed with formaldehyde and glutaraldehyde in PBS for 15 min. Cells were washed with PBS and incubated in osmium tetroxide in PBS for 10 min. Then, cells were washed repeatedly with MilliQ water for 2 hours. Cells were gradually transitioned into ethanol by increasing from 10% ethanol to 100% ethanol using steps of 10%. Upon reaching 100% ethanol, cells were incubated in fresh ethanol for 20 min twice. Next, cells were gradually transitioned into HMDS by increasing from 25% HMDS in ethanol to 100% HMDS using steps of 25%. When 100% HMDS was reached, cells were incubated in fresh HMDS for 20 min twice. Finally, the majority of HMDS volume was removed, and cells were air dried. If appropriate, hexane-suspended cathodophores were drop-cast onto the cells. Cells were sputter-coated with 80:20 Pt:Pd. Details provided in SI Section 18.

## Acknowledgements

This work was supported by the Scialog Award from the Gordon and Betty Moore Foundation, the Aramont Fellowship Fund for Emerging Science Research, NIH grant R01 GM146791, NIH grant R21 GM146127, and startup funds from Harvard University. J.B.C. was supported in part by the Harvard QBio Student Award. A.D. and I.M. were supported by the Harvard Systems Biology Internship Program. Harvard QBio and the Harvard Systems Biology Internship Program are supported by the NSF-Simons Center for the Mathematical & Statistical Analysis of Biology at Harvard University. K.S. was supported by the Harvard MCO SROH Internship Program and The Leadership Alliance’s Summer Research Early Identification Program. Part of this work was performed at the Harvard University Center for Nanoscale Systems (CNS), a member of the National Nanotechnology Coordinated Infrastructure Network (NNCI), which is supported by the National Science Foundation under NSF award no. ECCS-2025158. The authors thank Arvind Srinivasan, Ami Thakrar, Daphne-Eleni Archonta, Balmiki Kumar, and Ethan Garner for useful discussions, as well as James MacArthur for assistance with CL instrumentation. The authors are grateful to Adam Cohen, Rachelle Gaudet, Martin Gruebele, Prashant Jain, Catherine Murphy, Daniel Needleman, Bridget Queenan and Ariane Vertanian for critical reading of the manuscript.

## Author Contributions

S.A.R., J.B.C., and M.B.P. conceived the project, designed experiments, analyzed the data, and interpreted the results. S.A.R. developed the CL detection system and implemented the hardware-software integration for simultaneous SEM and CL imaging. S.A.R. developed imaging and data analysis pipeline. S.A.R. and J.B.C. performed CL imaging. J.B.C. optimized the nanocrystal synthesis protocol. J.B.C. performed TEM imaging of nanocrystals. J.B.C. performed cell culture and optimized the biological sample preparation protocol for SEM imaging. S.A.R., J.B.C., A.N., K.S., and I.M. synthesized nanocrystals. E.R.S. assisted with implementing automated scanning. S.A.R. and A.D. performed Monte Carlo simulations of electron trajectories. S.M. developed the probabilistic classification framework and analysis methods for color assignment in multicolor images. A.C. assisted with biological sample preparation for SEM imaging. S.A.R., J.B.C., A.N., K.S., I.M., and A.C. washed nanocrystals for SEM imaging. S.A.R., J.B.C., and M.B.P. wrote the manuscript. M.B.P. supervised the research.

