## Supplementary Information for "Lanthanide Cathodophores for Multicolor Electron Microscopy"

#### 1. Synthesis of cathodophores

**Materials:** Oleic acid (Sigma-Aldrich, Cat# 364525-1L, 90%, technical grade), 1-octadecene (SigmaAldrich, Cat# O806-1L, 90%, technical grade), lanthanide chlorides [samarium trichloride hexahydrate (Sigma-Aldrich, Cat# 204277-5G, ≥99.99%), gadolinium trichloride hexahydrate (Sigma-Aldrich, Cat# 203289-25G, 99.999%), terbium trichloride hexahydrate (Sigma-Aldrich, Cat# 212903-5G, 99.9%), dysprosium trichloride hexahydrate (Sigma-Aldrich, Cat# 289272-25G, 99.9%), holmium trichloride hexahydrate (Sigma-Aldrich, Cat# 289213-5G, 99.9%), yttrium trichloride hexahydrate (Sigma-Aldrich, Cat# 464317-25G, 99.99%)], sodium hydroxide (Sigma-Aldrich, Cat# 221465-500G, ACS Reagent, ≥97.0%, pellets), methanol (Sigma-Aldrich, Cat# 34860-1L-R, ≥99.9%), ammonium fluoride (Sigma-Aldrich, Cat# 216011-100G, ACS Reagent, ≥98.0%), n-hexane (Sigma-Aldrich, Cat# HX0302-3, 95%), ethanol (Ethyl alcohol, Sigma-Aldrich, Cat# 459844-1L, 200 proof, ACS Reagent, ≥99.5%).

**Core nanocrystals:** $NaSmF_4$, $NaTbF_4$, $NaDyF_4$, $NaHoF_4$, and $NaGdF_4$ core nanocrystals were synthesized by the coprecipitation method and based on previously reported protocols[1–3]. 0.5 mmol of the appropriate lanthanide chloride hydrate was combined with 3 mL of oleic acid and 7.5 mL of 1-octadecene in a 100 mL three-neck round-bottom flask. The flask was connected to a Schlenk line, with a heating mantle placed underneath and a thermocouple inserted into the flask through a vacuum-tight adapter. Both the heating mantle and the thermocouple were controlled by the Digi-Sense TC9600 temperature control module. Flask contents were stirred continuously using a magnetic egg-shaped stir bar. The reaction was placed under vacuum and the temperature was set to 160 °C for 30 min. The heating mantle was removed, and the solution was cooled to <30 °C, then exposed to air. Next, 1 M sodium hydroxide in methanol and 0.4 M ammonium fluoride in methanol were sonicated to homogeneity separately. A nucleation precursor solution was prepared by adding 1.25 mL of 1 M sodium hydroxide to 5 mL of 0.4 M ammonium fluoride. The resulting precursor solution was vortexed for 10 s and added dropwise to the lanthanide oleate solution in the three-neck round-bottom flask. The reaction was mixed for 60 min while exposed to air at room temperature. Methanol was evaporated by increasing the temperature to 70 °C, and the reaction temperature was maintained between 70 °C and 80 °C for 30 min while exposed to air. Then, for the nanocrystal growth step, the reaction was placed under an argon atmosphere and the temperature was increased to 320 °C at a rate of 18–28 °C/min (see **SI Fig. 1**). The reaction temperature was maintained for 60 min before cooling to <30 °C by removing the heating mantle. The entire volume of the reaction mixture

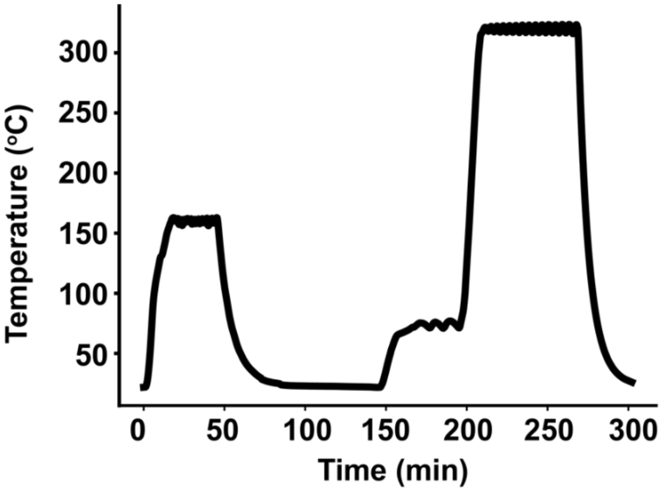

**SI Fig. 1:** Typical temperature profile of a lanthanide nanocrystal synthesis reaction measured using a thermocouple. The thermocouple was placed in an NMR tube inserted into the round-bottom reaction flask through a vacuum-tight adapter. Thermocouple measurements were recorded using a Digi-Sense TC9600 temperature controller.

was placed in a borosilicate glass scintillation vial. Core nanocrystals were stored as-synthesized in oleic acid and 1-octadecene at room temperature. For smaller nanocrystals, the synthesis was performed as described above, except that the 320 °C reaction temperature was maintained for either 15 min ($NaHoF_4$) or 2.5 min ($NaHo_{0.8}Lu_{0.2}F_4$), not 60 min, before cooling to <30 °C (SI Fig. 3). For dual-doped nanocrystals (e.g., $NaHo_{0.5}Dy_{0.5}F_4$ or $NaHo_XGd_{1-X}F_4$),

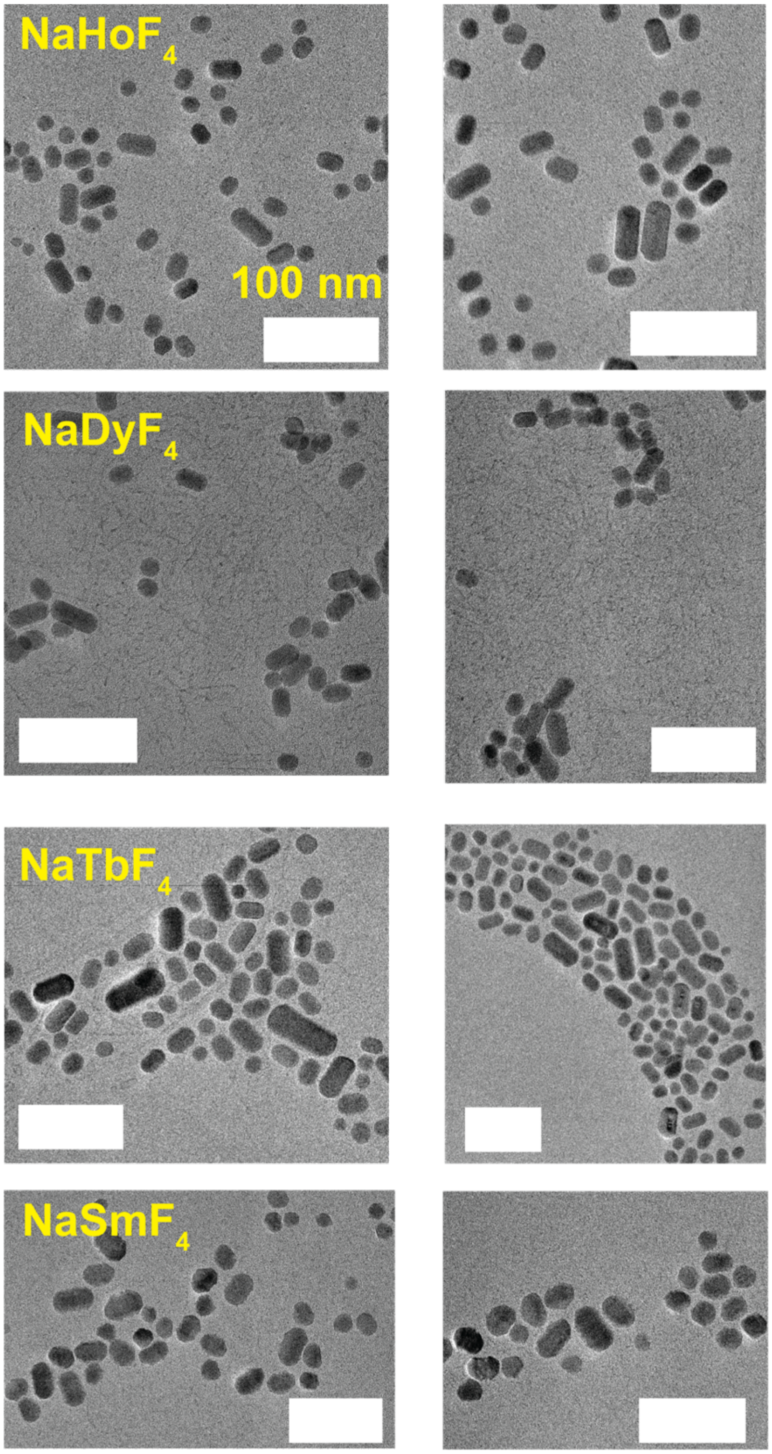

**SI Fig. 2:** TEM characterization of $NaHoF_4$, $NaDyF_4$, $NaTbF_4$, and $NaSmF_4$ core cathodophores synthesized using a 60 min growth step. Images were acquired at 200 keV beam energy using a JEOL JEM-F200 S/TEM in TEM mode. **Scale bars:** 100 nm.

synthesis was performed as described above, except that the 0.5 mmol of lanthanide chloride hydrates was divided into two quantities based on the desired molar ratio of the two lanthanide elements

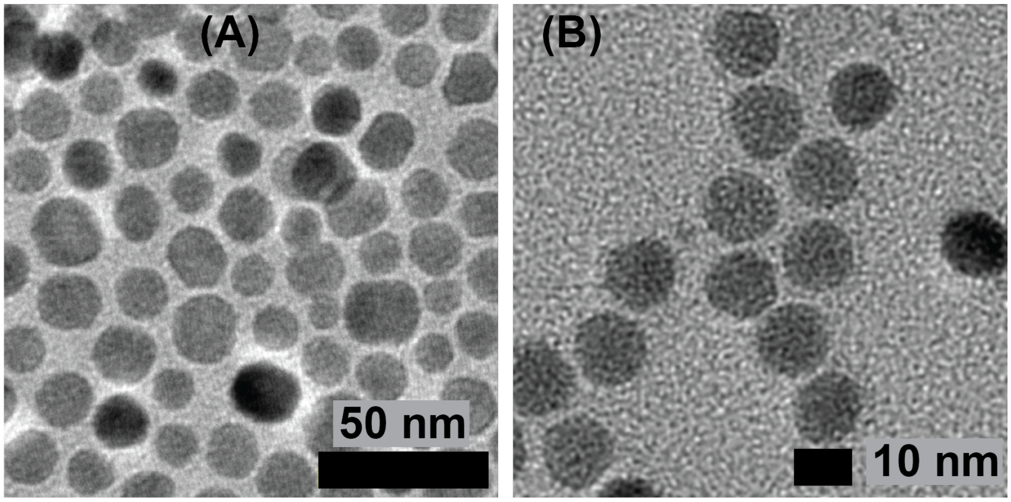

**SI Fig. 3: (A)** TEM image of $NaHoF_4$ core cathodophores synthesized using a 15 min growth step. (B) TEM image of a $NaHo_{0.8}Lu_{0.2}F_4$ cathodophores synthesiszed using a growth step of 2.5 min. Images were acquired at 200 keV beam energy using a JEOL JEM-F200 S/TEM in TEM mode.

**Core-shell nanocrystals:** Core-shell nanocrystals were synthesized by first synthesizing core nanocrystals as described previously. The entire volume of as-synthesized core nanocrystals was then washed by mixing with 5 mL of ethanol and centrifuging at 5,000 × g for 10 minutes. The pellet was resuspended in 1 mL of n-hexane, the ethanol wash was repeated once, and the pellet was resuspended in 5 mL of n-hexane. To begin shell synthesis, 0.5 mmol of the appropriate lanthanide chloride hydrate was combined with 3 mL of oleic acid and 7.5 mL of 1-octadecene in a 100 mL three-neck round-bottom flask. The flask was connected to a Schlenk line, with a heating mantle placed underneath and a thermocouple inserted into the flask through a vacuum-tight adapter. Both the heating mantle and the thermocouple were controlled by the Digi-Sense TC9600 temperature control module. Flask contents were stirred continuously using a magnetic egg-shaped stir bar. The reaction was placed under vacuum and the temperature was set to 160 °C for 30 min. The heating mantle was removed, and the solution was cooled to <30 °C, then exposed to air. Next, the entire volume of n-hexane-suspended core nanocrystals was added to the flask. 1 M sodium hydroxide in methanol and 0.4 M ammonium fluoride in methanol were sonicated to

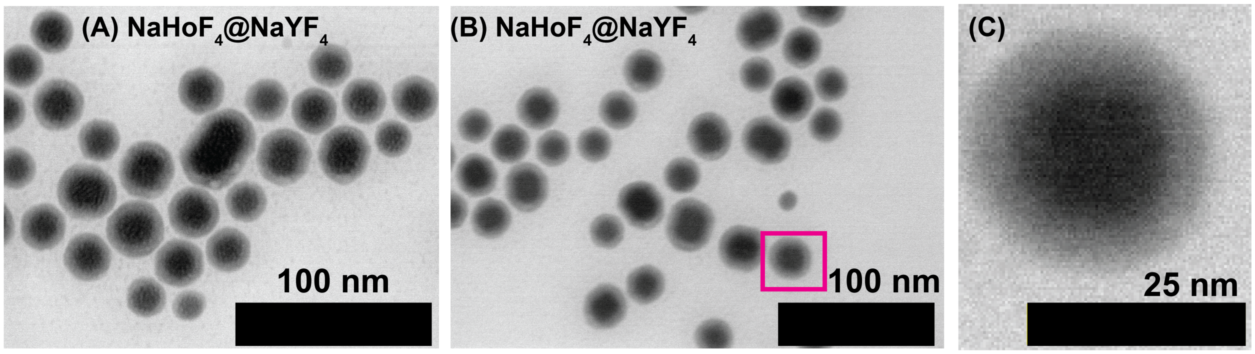

**SI Fig. 4:** TEM characterization of $NaHoF_4$@$NaYF_4$ core-shell cathodophores. Images were acquired at 200 keV beam energy using a JEOL JEM-2100 TEM.

homogeneity separately. A nucleation precursor solution was prepared by adding 1.25 mL of 1 M sodium hydroxide to 5 mL of 0.4 M ammonium fluoride. The resulting precursor solution was vortexed for 10 s and added dropwise to the flask. The reaction was mixed for 60 min while exposed to air at room temperature. n-Hexane and methanol were evaporated by increasing the temperature to 70 °C, and the reaction temperature was maintained between 70 °C and 80 °C for 30 min while exposed to air. Then, for the shell growth step, the reaction was placed under an argon atmosphere and the temperature was rapidly increased to 320 °C. The reaction temperature was maintained for 60 min before cooling to <30 °C by removing the heating mantle. The entire volume of the reaction mixture was placed in a borosilicate glass scintillation vial. Core-shell nanocrystals were stored as-synthesized in oleic acid and 1-octadecene at room temperature.

#### 2. Nanocrystal sample preparation

For nonlocal CL excitation experiments involving a dense nanocrystal region and a sparse nanocrystal region separated by a well-defined edge, the samples were prepared as follows: 0.5 mL of as-synthesized nanocrystals was washed by mixing with 0.5 mL of ethanol and centrifuging at 3,500 × g for 3 minutes. The pellet was resuspended in 0.5 mL of n-hexane, mixed with 0.5 mL of ethanol, and centrifuged again; this step was repeated for a total of five washes. After the fifth wash, the pellet was resuspended in 0.5 mL of n-hexane. A piece with a total area of ~1 $cm^2$ was cut from a p-type Si wafer (Ted Pella, Cat# 16015), cleaned with 70% ethanol, and plasma cleaned. Then, half of its surface was covered with tape (Scotch Magic Tape). 5 μL of the desired n-hexane-suspended nanocrystals was drop-cast onto the exposed half of the Si wafer's surface. After the n-hexane dried, drop-casting was repeated four times to create a thick layer of dense nanocrystals. Finally, the tape was removed from the Si wafer's surface, the desired n-hexane-suspended nanocrystals were diluted 100–1,000x in n-hexane to achieve sparsity, and 5 μL was drop-cast onto the wafer.

For experiments requiring samples of single nanocrystals with minimal aggregation, 0.5 mL of as-synthesized nanocrystals was washed by mixing with 5 mL of ethanol and centrifuging at 5,000 × g for 10 minutes. The pellet was resuspended in 1 mL of n-hexane, mixed with 5 mL of ethanol, and centrifuged again; this step was repeated for a total of five washes. After the fifth wash, the pellet was resuspended in 5 mL of n-hexane and the solution was left undisturbed overnight. For further sample preparation, nanocrystals were pipetted from the top of the solution to avoid collecting precipitated nanocrystal aggregates. For characterization by TEM, 2 μL of the nanocrystal solution was drop-cast onto a copper TEM grid (Ted Pella, Cat# 01810); for characterization by SEM, 5 μL of the nanocrystal solution was drop-cast onto an ethanol- and plasma-cleaned p-type Si wafer (total area ~1 $cm^2$).

#### 3. CL detection system

CL was collected by a parabolic mirror installed inside the vacuum chamber of a ZEISS SUPRA 55VP SEM. See **SI Fig. 5** for a schematic of the setup. The mirror was installed on a five-axis nanopositioning stage system (Attocube) to precisely align the center of the field-of-view (FOV) of the SEM to the focal point of the mirror. The mirror was custom-manufactured by diamond turning (B-con Engineering). The mirror had a focal length of 1 mm and collected CL over an average solid angle of ~1.34π steradians. The mirror collimated light from the sample and directed it outside the vacuum chamber through an anti-reflection-coated fused silica flat window (Thorlabs, VPW42-A) installed on the custom-modified side port of the SEM. The light

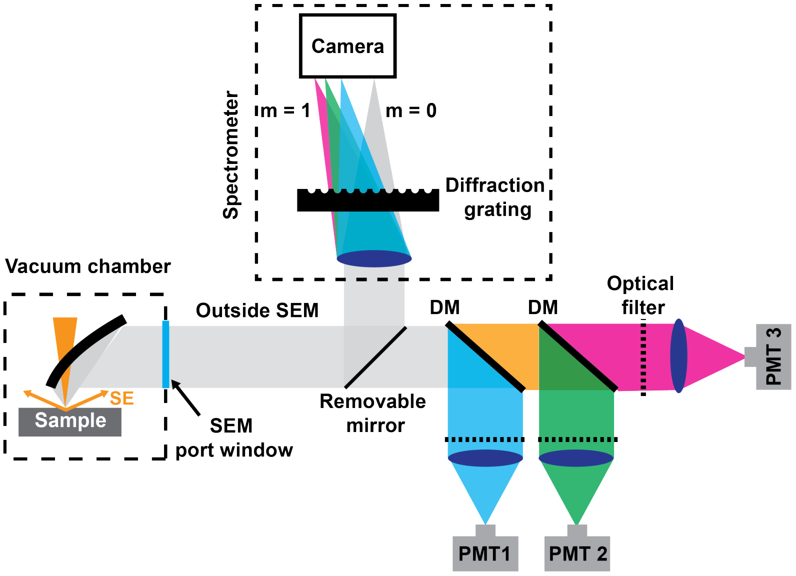

**SI Fig. 5:** Schematic of the CL detection system. A parabolic mirror was installed inside the vacuum chamber of the SEM to collect light from the sample. The mirror had a 500-µm-diameter aperture through which the electron beam passed to reach the sample. The mirror directed light outside the vacuum chamber, where a system of dichroic mirrors (DMs), optical filters, and lenses was used to focus it onto PMTs. Three PMTs were used to simultaneously collect light over three spectral ranges. To collect spectra, a removable mirror was inserted in the emission path to direct light towards the spectrometer consisting of a diffraction grating and an sCMOS camera.

was then spectrally separated using dichroic mirrors, filtered using band-pass filters matched to the emission peaks of the cathodophores (see **SI Fig. 6**), and focused onto PMTs (Hamamatsu, H7421-40) using 30-mm-focal-length tube lenses (Thorlabs, AC254-030-A-ML).

**Custom spectrometer:** For spectral measurements, a fold mirror was inserted in the emission path to redirect the CL towards the spectrometer. The light was focused onto an sCMOS camera (Hamamatsu, Orca-Fusion BT) using a 100-mm-focal-length lens (Thorlabs, AC254-100-A-ML). A diffraction grating (Thorlabs, GT13-03) was placed in front of the sCMOS camera to spectrally separate the CL signal. See SI section 16 for details on the acquisition of single particle spectra.

**Commercial spectrometer for ensemble measurements:** The spectra of the ensemble of nanocrystals were measured using a commercial spectrometer (Thorlabs, CCS200). Collimated light from the parabolic mirror was focused on the multimode fiber of the spectrometer. The accompanying software was used to obtain the spectra. Ensemble spectra were obtained by scanning a 400 nm x 400 nm region of dense cathodophores with a beam energy of 3 keV.

Note that the ensemble spectra presented in **Fig. 1C** and **SI Fig. 6** were obtained using the commercial spectrometer. All the other spectra presented in this work were obtained using the custom spectrometer.

#### 4. Image acquisition

**Cathodophore characterization:** Images were acquired using custom software written in LabVIEW. The software communicated with the hardware (SEM and the PMTs) through an NI DAQ card (National Instruments, PXIe-6368). Scanning of the electron beam, collection of CL signal on three PMTs, and collection of electron signal on the Everhart-Thornley detector of the SEM (SE channel) were synchronized. We navigated the sample using the ZEISS SmartSEM

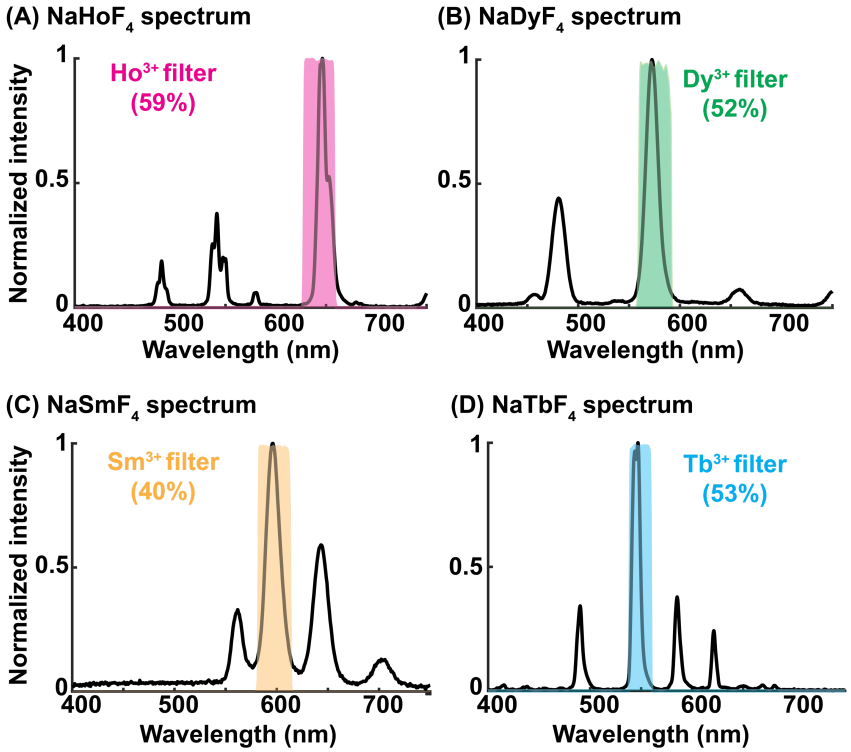

**(E) Transmission of cathodophores across different filters**

| Filter | Details | $NaHoF_4$ transmission | $NaDyF_4$ transmission | $NaSmF_4$ transmission | $NaTbF_4$ transmission |
|---|---|---|---|---|---|
| **$Ho^{3+}$ filter** | Chroma ET645/30X | 59% | 3% | 24% | 1% |
| **$Dy^{3+}$ filter** | Chroma FF03-575/25 | 3% | 52% | 18% | 14% |
| **$Sm^{3+}$ filter** | Semrock FF01-598/25 | 1.2% | 7% | 40% | 15% |
| **$Tb^{3+}$ filter** | Chroma ET550/20X | 14% | 2% | 4% | 53% |

**SI Fig. 6: (A–D)** CL emission spectra of $NaHoF_4$, $NaDyF_4$, $NaSmF_4$, and $NaTbF_4$ cathodophores, respectively. Spectra were obtained from an ensemble of cathodophores. Shaded bands in each subpanel show transmission profiles of the band-pass emission filters matched to the emission peaks of the respective dopants. **(E)** Information about the band-pass emission filters. CL transmission efficiency for each dopant across the filters is also shown.

software. Once a region of interest was identified, we switched to the “External Scan” setting of the SEM. In this setting, the electron beam scanning was controlled by the LabVIEW software. For each FOV, the oversampled SEM image from SmartSEM was saved, which was later used to determine the size of cathodophores. Furthermore, during each scan of the FOV with the “External Scan” setting, four images were acquired: one SE image and three CL images (in different spectral channels).

For the characterization of emission properties of cathodophores **(Fig. 4A–E, Fig. 5, Fig. 6C)**, CL images were acquired with a single cathodophore per image. We scanned a small region, 20x20 pixels, around the cathodophore with a pixel size of 4–6 nm, meeting the Nyquist criterion for the smallest cathodophores. Such an image acquisition pipeline, i.e., scanning a small region around nanocrystals, was important to reduce overall imaging times because the effective dwell times in CL imaging were ~3 orders of magnitude slower than typical SEM imaging. In summary, for each FOV two types of images were acquired:

1. An oversampled SE image using the ZEISS Smart SEM software at regular scan speeds of the SEM (dwell time: 1–3 μs). This image was used to determine the size of cathodophores.
2. A Nyquist-sampled SE and three CL images of the same FOV with longer dwell times. These images were used to characterize the emission properties of cathodophores.

**Multicolor imaging:** Images were acquired using the setup described in **SI Fig. 5**. Multiple images were taken at a shorter dwell time to minimize charging related effects. An increase in charging was noted for denser samples used in multicolor experiments, which may be attributed to a higher concentration of organic solvents left behind after washing. To accommodate this variability, we varied the beam dwell time between 100 μs and 1 ms, based on the extent of charging. Images were acquired with an effective beam dwell time of 30–50 ms per pixel. Drift correction was performed post-image acquisition.

#### 5. Experiments to quantify nonlocal CL signal

The experiments discussed in **Fig. 2** were performed with a tube lens of 30 mm focal length, which resulted in a FOV >300 μm (>150 μm radius) as shown in **SI Fig. 7A**. Hence, CL signal from the sample region within 150 μm of the excited cathodophore was collected by the PMTs. We also reduced the FOV of the imaging system to confirm that the nonlocal CL signal was in fact originating from the sample, and was being collected by the parabolic mirror. To this end, we modified the FOV to ~25 μm radius by changing the focal length of the tube lens to 400 mm **(SI Fig. 7A)**. When we repeated the experiment with dense $NaDyF_4$ and sparse $NaHoF_4$ cathodophores, the nonlocal signal in the $Dy^{3+}$ color channel disappeared when $NaHoF_4$ cathodophores were imaged at a distance >20 μm away from the edge **(SI Fig. 7B)**. However, signal in the $Ho^{3+}$ color channel was independent of the distance from the edge, which was expected if this signal originated from the excited cathodophore placed at the center of the FOV **(SI Fig. 7C)**.

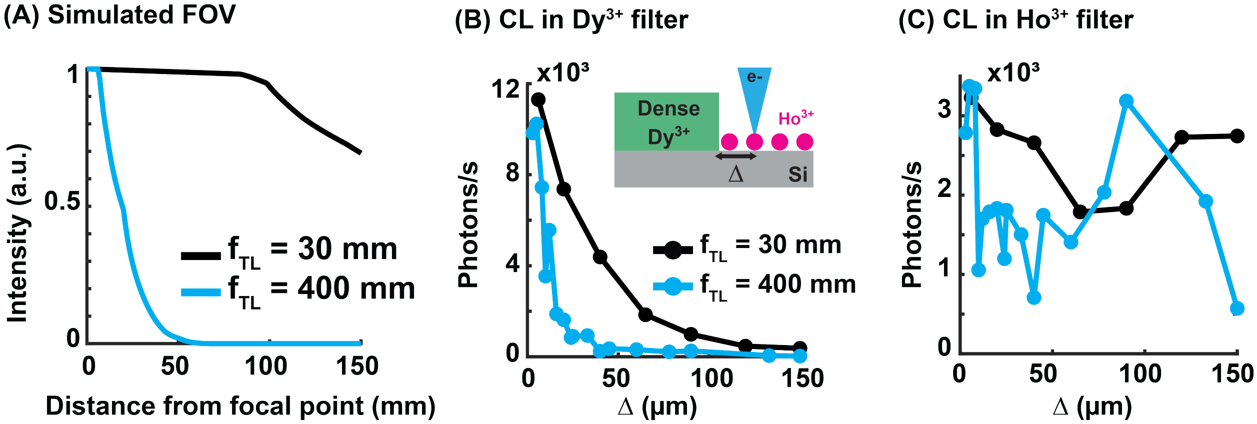

**SI Fig. 7: (A)** Simulation of light captured from a point source on a PMT of 5 mm diameter (matched to the PMTs used in this work). The point source was imaged at different distances from the focal point of the parabolic mirror for two different focal lengths of the tube lens (lenses placed in front of the PMTs in **SI Fig. 5**). The intensity was obtained by ray tracing and served as a measure of the field of view (FOV) of the CL imaging system. The FOV had >150 μm radius for the 30 mm focal length tube lens, which was reduced to ~25 μm for the 400 mm focal length tube lens. **(B, C)** Edge experiment with dense $NaDyF_4$ and sparse $NaHoF_4$ cathodophores. Sparse $NaHoF_4$ cathodophores were imaged at different distances from the dense region's edge. CL in $Dy^{3+}$ and $Ho^{3+}$ color channels are shown in (B) and (C), respectively.

#### 6. Two-color imaging

**SI Fig. 8** shows representative examples of images of proximal $NaHoF_4$ and $NaDyF_4$ cathodophores. Despite their proximity, the cathodophores could be distinguished using spatially separated emission in the $Ho^{3+}$ and $Dy^{3+}$ color channels. **SI Fig. 9** shows examples of two-color imaging in dense samples of $NaHoF_4$ and $NaDyF_4$ cathodophores.

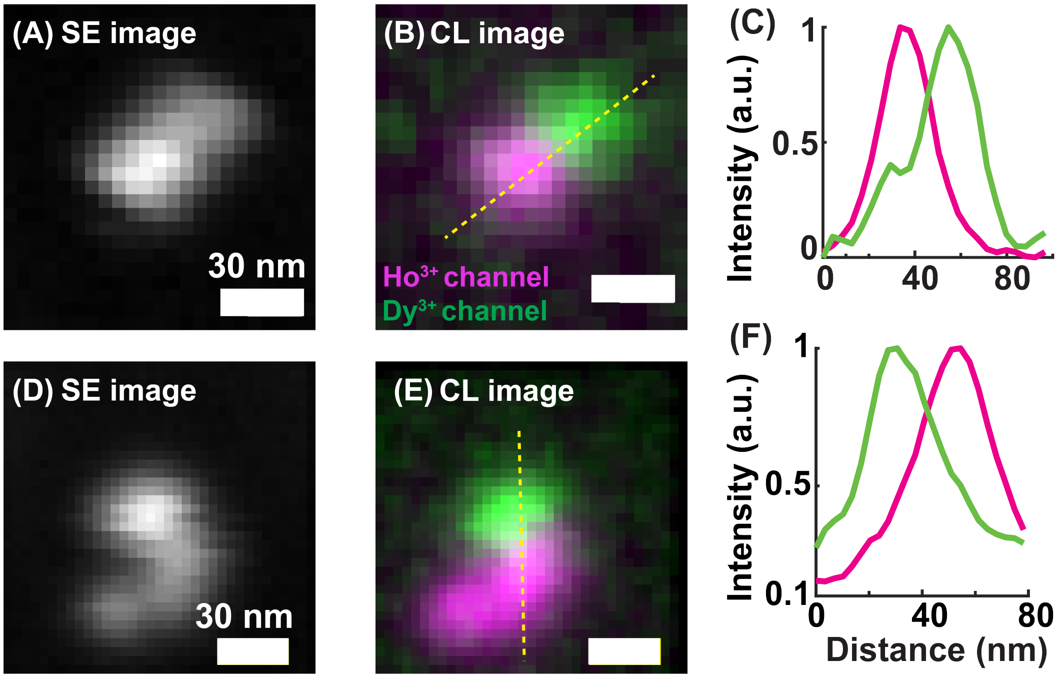

**SI Fig. 8: (A, D)** SE images of two adjacent cathodophores. **(B, E)** CL images of the cathodophores in (A) and (D) respectively, obtained by merging signals in $Ho^{3+}$ color channel (magenta) and $Dy^{3+}$ color channel (green). **(C, F)** Cross-sectional profiles of the CL images along the lines shown in (B) and (E), respectively.

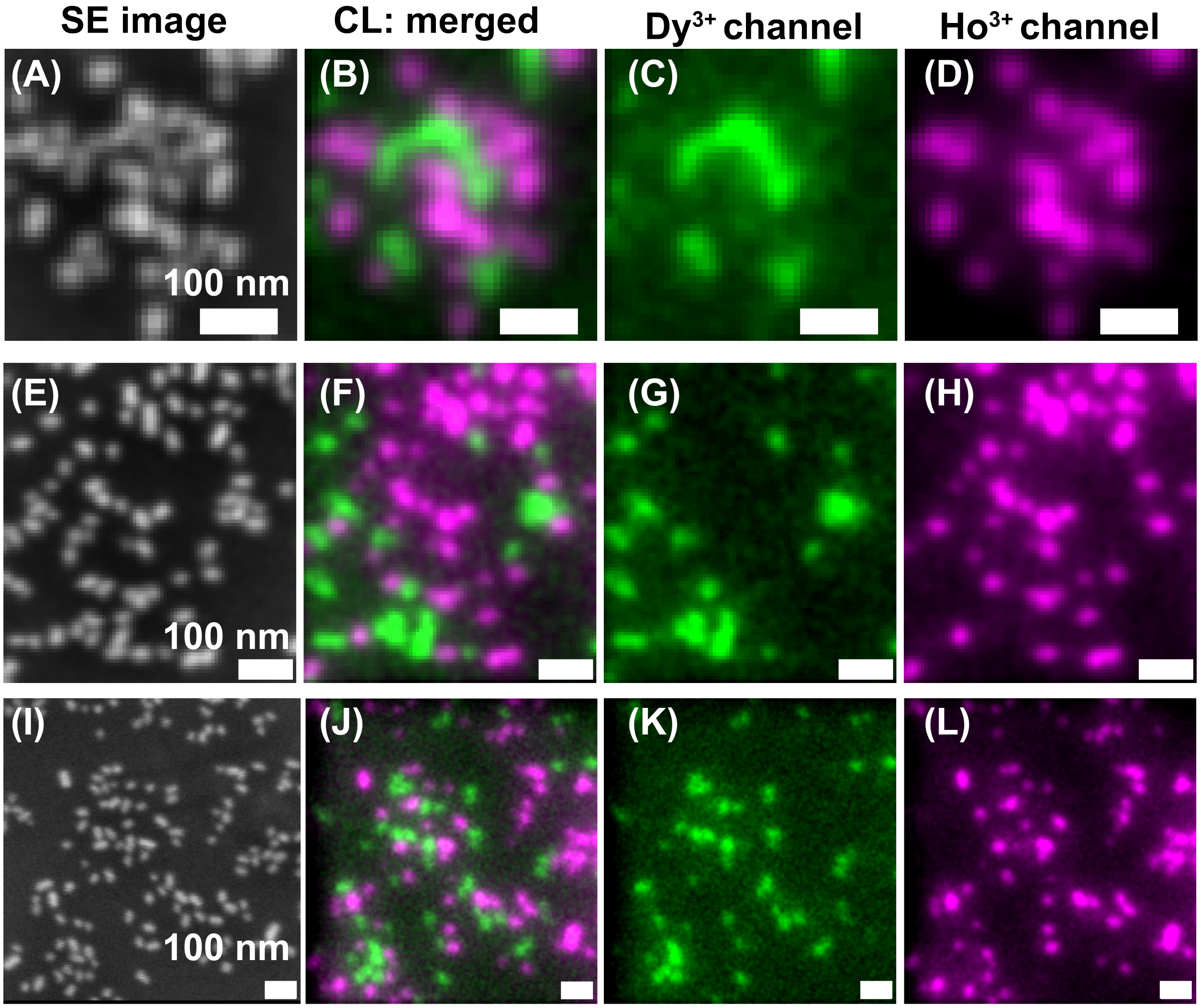

**SI Fig. 9: (A, E, I)** SE images of dense monolayers of samples containing $NaHoF_4$ and $NaDyF_4$ cathodophores. **(B, F, J)** The corresponding CL images. CL images were obtained by merging CL images from the respective **(C, G, K)** $Dy^{3+}$ and **(D, H, L)** $Ho^{3+}$ color channels.

#### 7. Three-color imaging

**SI Fig. 10** shows the three-color image from **Fig. 3,** along with the three CL spectral channels. More examples of three-color CL images are shown in **SI Fig. 11.** These images were obtained by imaging $NaHoF_4$, $NaDyF_4$ and $NaTbF_4$ cathodophores in one FOV.

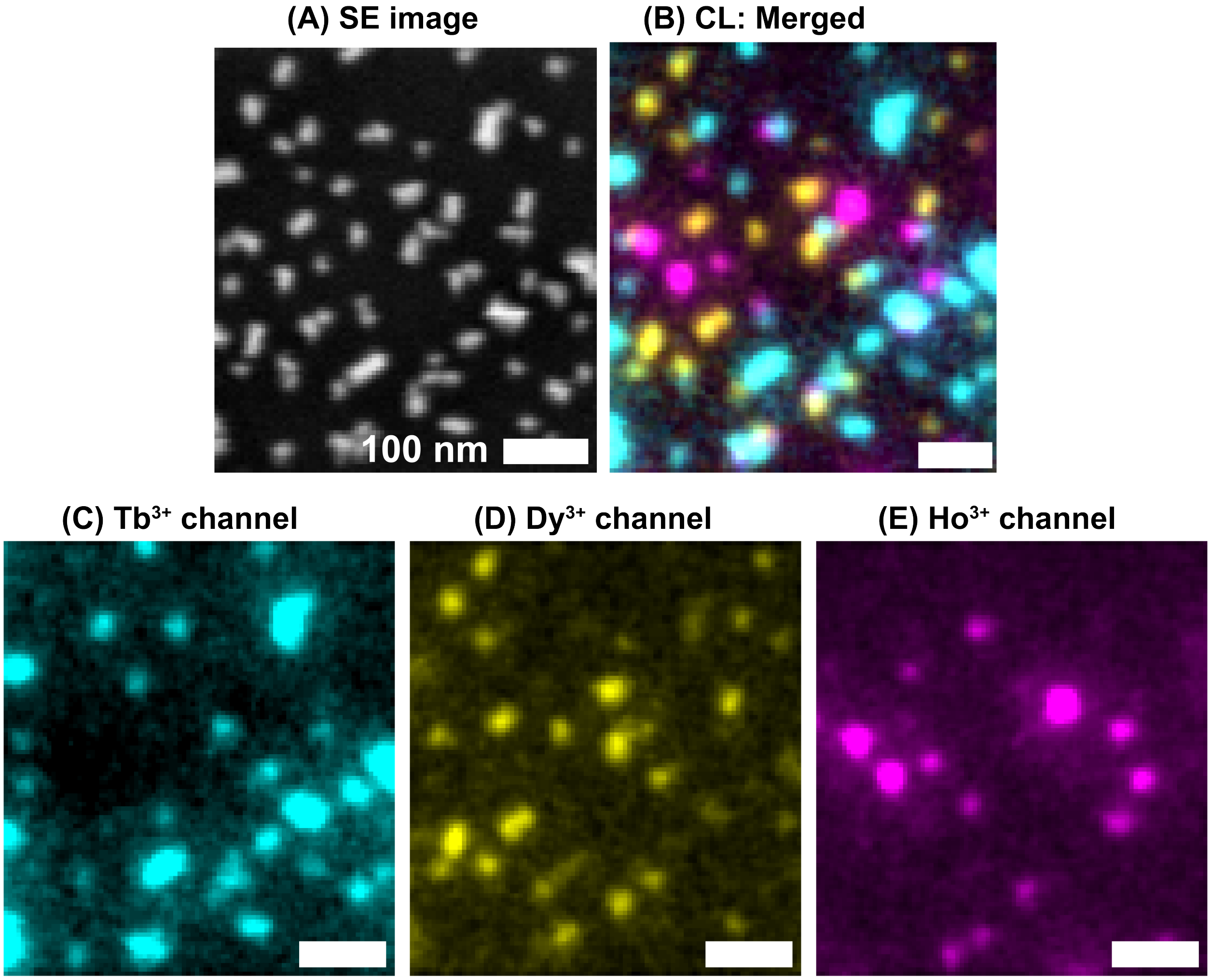

**SI Fig. 10:** Additional information on three-color imaging shown in **Fig. 3K–N**. **(A)** SE image of a dense monolayer containing $NaHoF_4$, $NaDyF_4$, and $NaTbF_4$ cathodophores. **(B)** The corresponding CL image obtained by merging CL images from **(C)** $Tb^{3+}$, **(D)** $Dy^{3+}$, and **(E)** $Ho^{3+}$ color channels.

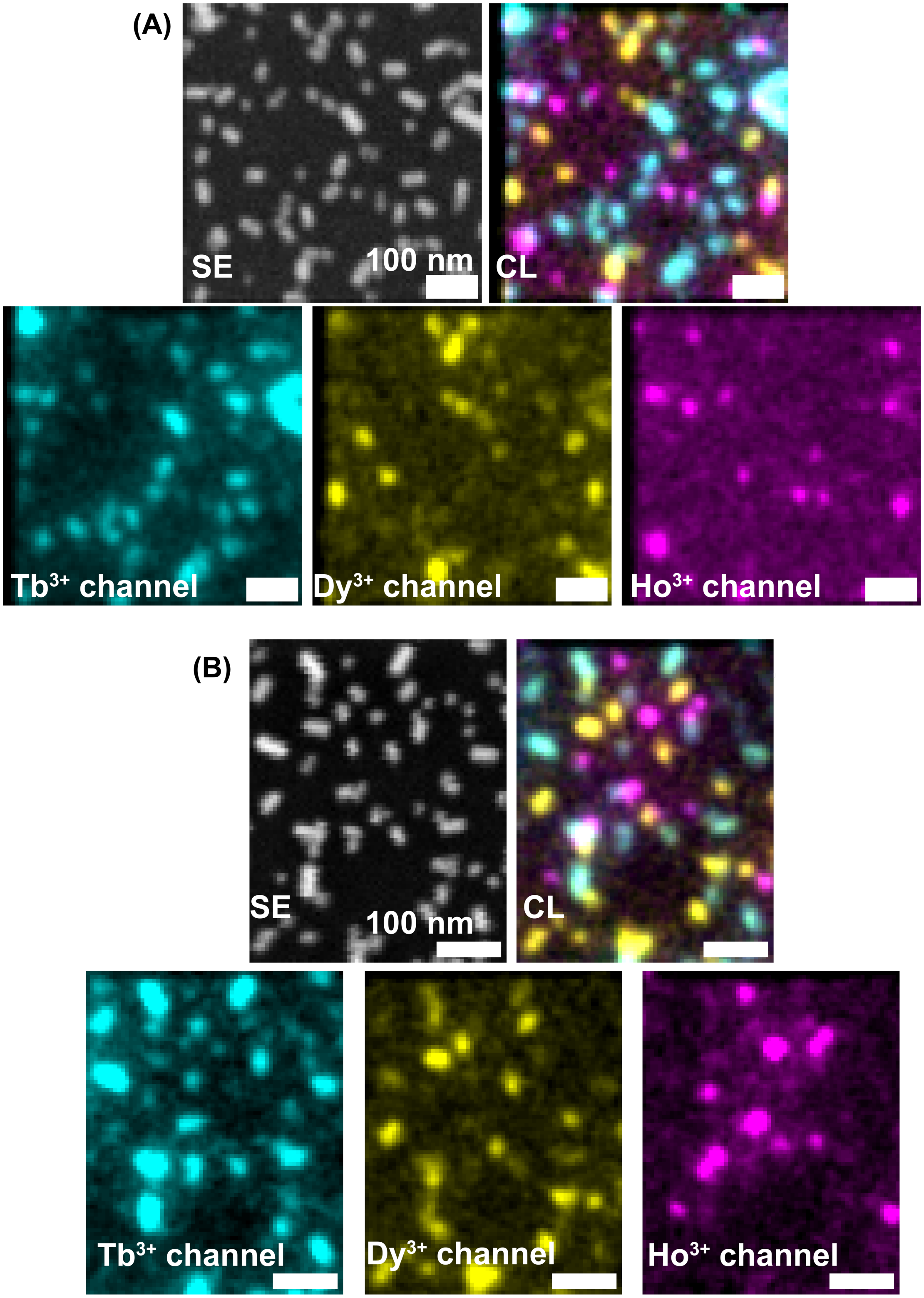

**SI Fig. 11:** Additonal representative examples of three-color imaging using $NaHoF_4$, $NaDyF_4$, and $NaTbF_4$ cathodophores.

### 8. Automated dopant assignment in dense multicolor samples

#### Approach

We have developed a probabilistic classification model based on Bayesian inference for automatically assigning dopants to cathodophores given the CL detection rates measured in $Ho^{3+}$ and $Dy^{3+}$ spectral channels **(SI Fig. 12)**.

#### Model training

First, we trained our classification model using CL detection rates from single-cathodophore measurements. Specifically, we fitted a bivariate Gaussian distribution to the SE image of a cathodophore to obtain the cathodophore's position and size. We used these results as initial guesses in the following constrained optimization problem. Namely, in each CL channel separately, we used maximum likelihood estimation (MLE) to obtain the estimators of a bivariate Gaussian distribution modeling the cathodophore's CL signal. Noise was assumed to follow Poisson statistics. The CL detection rates were defined as the amplitudes of the obtained bivariate Gaussian distributions divided by the pixel dwell time.

We then compiled the detection rates from a set of $NaHoF_4$ single-cathodophores (N = 113) in each spectral channel to model the relationship between the detection rate versus cathodophore diameter and beam dwell time for each channel. Based on these results, we further modeled the distribution of CL detection rates for these $NaHoF_4$ cathodophores. For a given dwell time and diameter, we modeled the distribution of CL detection rates by a bivariate Gaussian distribution, whose location vector consisted of the expected detection rates in each spectral channel, and whose covariance matrix reflected the variability in the rates in each channel. We used the obtained distribution as a likelihood function to compute the probability that a cathodophore of a given diameter, at a given dwell time, was $Ho^{3+}$-doped. We then repeated this analysis using a set of $NaDyF_4$ single-cathodophores (N = 65), to obtain a likelihood function for $Dy^{3+}$-doped cathodophores.

#### Dopant assignment

**Probabilistic dopant assignment – for cathodophores whose CL detection rates could be measured in both spectral channels:** We assigned a dopant to a cathodophore by computing the probability that the cathodophore was made of each possible dopant, given the detection rates in each spectral channel, and selecting the dopant corresponding to the greatest of these probabilities. Specifically, we applied Bayes' theorem, using as prior probabilities the known fractions of each dopant in the sample (e.g., 50% for each $Ho^{3+}$ and $Dy^{3+}$ in **SI Fig. 12)**; as likelihood functions, bivariate Gaussian distributions whose location vectors and covariance matrices had been obtained during model training (as explained above); evidence probability calculated using the law of total probability; and as input evidence, the CL detection rates measured in each spectral channel.

**Heuristic classification rules – for cathodophores whose CL detection rates could be measured at most in one spectral channel:** We applied the following set of rules to assign—or not—a dopant to a cathodophore, based on whether, in each spectral channel, a p-value could be computed, and a local maximum was detected (see below for more details on p-values and local maxima).

1. If no p-value could be computed in any channel: no dopant was assigned.
2. If a p-value could be computed in only one channel: the dopant was assigned corresponding to the channel of the only computed p-value (e.g., $Ho^{3+}$ dopant was assigned if the only p-value computed was from the $Ho^{3+}$ channel).
3. If a p-value could be computed in both channels, and:

a. If no local maximum was detected in either channel: the dopant corresponding to the channel with the smallest p-value was assigned.
b. If a local maximum was detected in only one channel: the dopant corresponding to the channel with the local maximum was assigned.
c. If local maxima were detected in both channels: probabilistic dopant assignment was performed.

#### Estimating CL detection rates in dense multi-cathodophore samples

The main challenge, in dense multi-cathodophore samples, was to minimize the bias that could be induced by neighboring cathodophores on the photon measurements from the cathodophore of interest. To address this issue, we used cathodophore-specific masks. To this end, we started by modelling the ensemble of cathodophores and the background in a multi-cathodophore SE image (e.g., **SI Fig. 12A**) to obtain a model of the image composed of an ensemble of bivariate Gaussian distributions on a quadratic surface (**SI Fig. 12B**). We then defined a cathodophore mask as the ensemble of pixels within 2-$\sigma$ of the distribution center (with $\sigma$ the standard deviation of the Gaussian distribution) and that were not shared with any other cathodophore mask (**SI Fig. 12C**).

Equipped with these cathodophore-specific masks, we assessed the statistical significance and whether a local maximum could be detected for each cathodophore, in each spectral channel of the CL micrograph (**SI Fig. 12D**). First, we assessed, for each cathodophore and in each spectral channel, whether a p-value could be computed and whether a local maximum could be detected. Specifically, we attempted to run a chi-square goodness-of-fit test to compare the statistical significance of the CL pixel values under the cathodophore mask versus the CL pixel values in the local background near the cathodophore; however, if the test had less than one degree of freedom, it could not be run, and the p-value could not be obtained. We also assessed whether a local maximum could be detected in the CL pixel values under the cathodophore mask by fitting linear and quadratic surfaces to them: if the adjusted R-squared of the fitted quadratic surface was the largest, and if both coefficients in the quadratic terms were negative, then we considered that a local maximum was detected.

For cathodophores for which both p-values could be computed and local maxima were detected, we estimated the CL detection rates in each spectral channel (**SI Fig. 12 E–F**). The analysis was similar to that performed to estimate the CL detection rates of single-cathodophores. Specifically, in each channel separately, we performed MLE assuming a Poisson noise, to obtain the estimators of a bivariate Gaussian distribution modeling the cathodophore's CL pixel values under the cathodophore mask and in the local background near the cathodophore. **SI Fig. 12 E–F** illustrate how cathodophore-specific masks helped minimize the potential biases induced by neighboring cathodophores on the estimation of the CL signal from the cathodophore of interest, even when the neighboring cathodophores were bright.

Finally, we performed dopant assignment using our probabilistic classification model, based on the estimated CL detection rates in each spectral channel, as detailed above (**SI Fig. 12 G–H**).

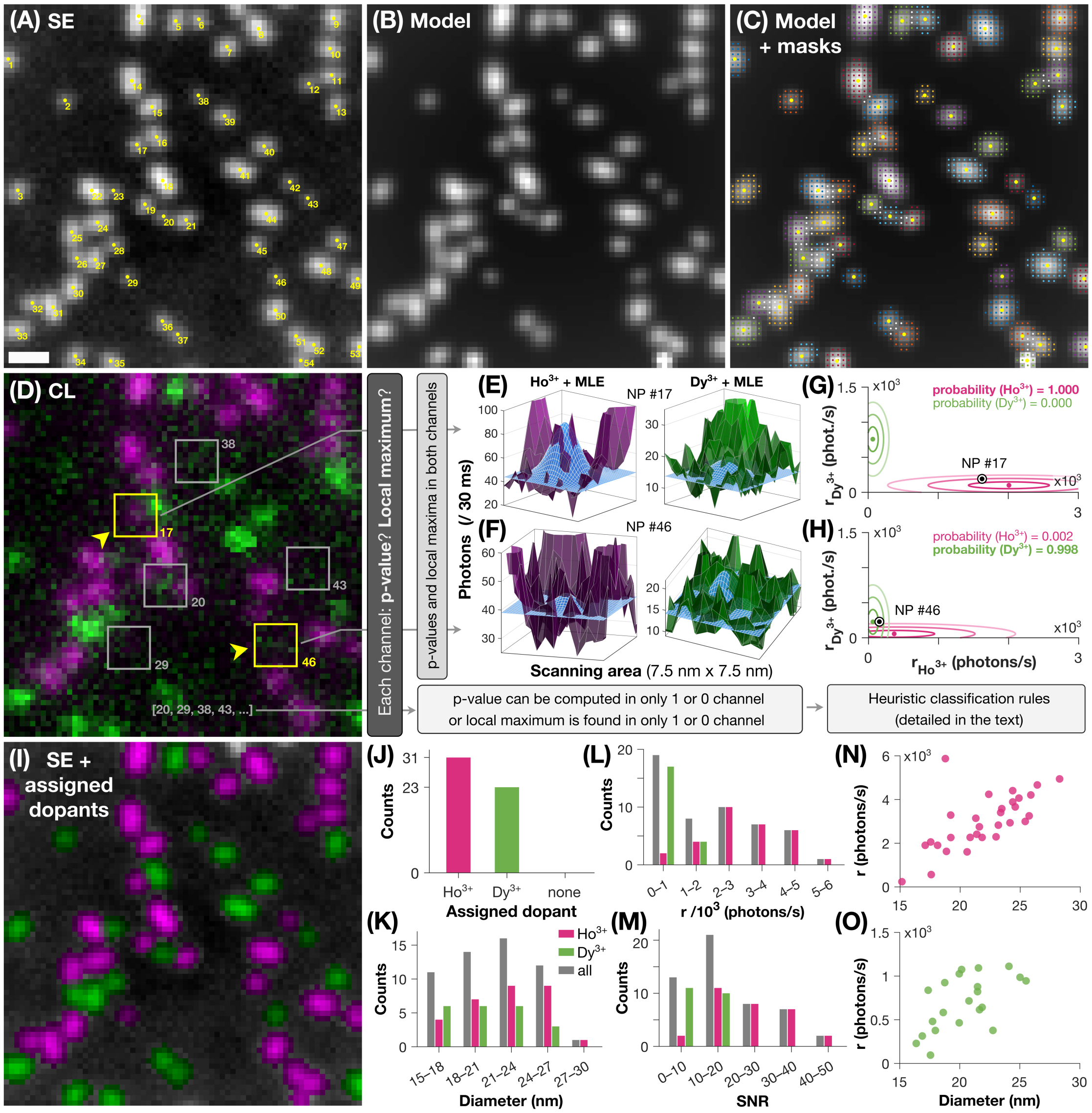

**SI Fig. 12: (A)** SE image containing both $NaHoF_4$ and $NaDyF_4$ cathodophores (identical to **Fig. 3G**) with cathodophore numerical identifiers overlaid. **(B)** Model for the SE image in (A), in which each cathodophore is represented as a bivariate Gaussian distribution and the background is represented as a quadratic surface. **(C)** Cathodophore-specific masks (small color dots) and centers (large yellow dots) obtained from, and overlaid on, the image model in (B). A cathodophore mask was defined as the ensemble of pixels within two standard deviations of the center of its Gaussian distribution, and that were not shared with any other cathodophore mask. Pixels that were shared between at least two cathodophore masks are shown as small white dots. **(D)** CL image showing signal in both $Ho^{3+}$ and $Dy^{3+}$ channels (identical to **Fig. 3H**). The yellow boxes highlight the two cathodophores (#17 and #46) whose probabilistic dopant assignment analyses are detailed in (E–H). The yellow arrowheads indicate the viewing angles used in (E) and (F). The gray boxes show cathodophores whose dopants were assigned by following heuristic classification rules. **(E–F)** Estimation of CL detection rates. CL signals in the $Ho^{3+}$ (left, magenta shades) and $Dy^{3+}$ (right, green shades) channels for cathodophores #17 (E) and #46 (F), and the associated bivariate Gaussian distributions (blue) resulting from maximum likelihood estimation (MLE) of the CL signals under the cathodophore-specific masks. The masks helped to reduce biases on the maximum likelihood estimates that neighboring cathodophores could otherwise induce. **(G–H)** Bayesian inference-based probabilistic dopant assignment. The estimated detection rates, $r_{Ho3+}$ and $r_{Dy3+}$, of cathodophore #17 (G) and #46 (H) (black-and-white dots) are compared to the detection rate statistics in our single-cathodophore training datasets (color ellipses and dots; magenta, $NaHoF_4$ single-cathodophores; green, $NaDyF_4$ single-cathodophores). The concentric ellipses represent the 1-σ, 2-σ, and 3-σ (innermost to outermost; σ is the standard deviation) levels of the bivariate Gaussian distributions with which we modeled CL detection rates in the training datasets. The color dots represent the expected values. Note that the likelihood function in each channel depended on the cathodophore size, and thus differed between cathodophores #17 and #46. We assigned to each cathodophore the dopant with the greatest posterior probability. **(I)** SE image colorized according to assigned dopant (magenta, assigned $Ho^{3+}$; green, assigned $Dy^{3+}$). **(J–M)** Multicolor sample characterization (see also **SI Table 1**). **(J)** Statistics of dopant assignment. **(K)** Occurrences of cathodophore diameters. **(L)** Occurrences of estimated cathodophore CL detection rates in the channel of the assigned dopant. **(M)** Occurrences of signal-to-noise ratio (SNR) in the channel of the assigned dopant. **(N, O)** CL detection rates in the channel of the assigned dopant as a function of cathodophore diameter for $Ho^{3+}$-doped (N) and $Dy^{3+}$-doped (O) cathodophores. (A–D, I) Contrast was adjusted. **Scale bar:** 50 nm.

| Cathodophore # | 1 | 2 | 3 | 4 | 5 | 6 | 7 | 8 | 9 | 10 | 11 | 12 | 13 | 14 | 15 |
|---|---|---|---|---|---|---|---|---|---|---|---|---|---|---|---|
| **Assigned dopant** | $Ho^{3+}$ | $Dy^{3+}$ | $Dy^{3+}$ | $Ho^{3+}$ | $Dy^{3+}$ | $Ho^{3+}$ | $Ho^{3+3+}$ | Ho | $Ho^{3+}$ | $Ho^{3+}$ | $Ho^{3+}$ | $Ho^{3+}$ | $Dy^{3+}$ | $Ho^{3+}$ | $Ho^{3+}$ |
| **Diameter (nm)** | 21.8 | 16.9 | 21.6 | 18.8 | 20.2 | 17.5 | 21.3 | 25.9 | 19.2 | 23.2 | 24.2 | 20.6 | 21.5 | 26.4 | 23.4 |
| **Photons/s** | 2265 | 314 | 615 | 5872 | 1075 | 2057 | 3143 | 4208 | 3287 | 2827 | 2935 | 1604 | 878 | 4667 | 3404 |
| **SNR** | 20 | 5 | 10 | 47 | 14 | 13 | 22 | 34 | 26 | 26 | 24 | 12 | 12 | 39 | 28 |

| Cathodophore # | 16 | 17 | 18 | 19 | 20 | 21 | 22 | 23 | 24 | 25 | 26 | 27 | 28 | 29 | 30 |
|---|---|---|---|---|---|---|---|---|---|---|---|---|---|---|---|
| **Assigned dopant** | $Dy^{3+}$ | $Ho^{3+}$ | $Ho^{3+}$ | $Ho^{3+}$ | $Dy^{3+}$ | $Ho^{3+}$ | $Ho^{3+}$ | $Dy^{3+}$ | $Ho^{3+}$ | $Ho^{3+}$ | $Dy^{3+}$ | $Dy^{3+}$ | $Dy^{3+}$ | $Ho^{3+}$ | $Ho^{3+}$ |
| **Diameter (nm)** | 25.0 | 18.9 | 28.3 | 20.8 | 18.0 | 23.0 | 24.9 | 22.8 | 24.4 | 25.7 | 20.0 | 25.5 | 20.8 | 15.2 | 22.4 |
| **Photons/s** | 986 | 1626 | 4943 | 2267 | 377 | 2293 | 4065 | 378 | 3888 | 3253 | 1026 | 945 | 716 | 238 | 4239 |
| **SNR** | 12 | 14 | 44 | 16 | 5 | 17 | 34 | 6 | 35 | 25 | 10 | 12 | 9 | 2 | 32 |

| Cathodophore # | 31 | 32 | 33 | 34 | 35 | 36 | 37 | 38 | 39 | 40 | 41 | 42 | 43 | 44 | 45 |
|---|---|---|---|---|---|---|---|---|---|---|---|---|---|---|---|
| **Assigned dopant** | $Ho^{3+}$ | $Ho^{3+}$ | $Dy^{3+}$ | $Dy^{3+}$ | $Dy^{3+}$ | $Dy^{3+}$ | $Dy^{3+}$ | $Dy^{3+}$ | $Dy^{3+}$ | $Dy^{3+}$ | $Dy^{3+}$ | $Ho^{3+}$ | $Dy^{3+}$ | $Ho^{3+}$ | $Ho^{3+}$ |
| **Diameter (nm)** | 24.6 | 18.1 | 21.6 | 18.7 | 17.4 | 17.7 | 20.0 | 16.8 | 21.9 | 21.5 | 24.1 | 17.6 | 17.6 | 24.4 | 17.1 |
| **Photons/s** | 3661 | 1905 | 1093 | 925 | 838 | 481 | 465 | n/a | 642 | 822 | 1112 | 564 | 95 | 4406 | 1905 |
| **SNR** | 30 | 13 | 14 | 12 | 8 | 6 | 6 | n/a | 10 | 11 | 17 | 6 | 1 | 37 | 15 |

| Cathodophore # | 46 | 47 | 48 | 49 | 50 | 51 | 52 | 53 | 54 |
|---|---|---|---|---|---|---|---|---|---|
| **Assigned dopant** | $Dy^{3+}$ | $Dy^{3+}$ | $Ho^{3+}$ | $Ho^{3+}$ | $Ho^{3+}$ | $Ho^{3+}$ | $Ho^{3+}$ | $Dy^{3+}$ | $Ho^{3+}$ |
| **Diameter (nm)** | 16.4 | 18.6 | 23.5 | 19.2 | 21.6 | 21.4 | 25.4 | 20.3 | 21.1 |
| **Photons/s** | 229 | 583 | 3582 | 2261 | 2752 | 2409 | 2996 | n/a | n/a |
| **SNR** | 3 | 8 | 31 | 17 | 25 | 19 | 19 | n/a | n/a |

**SI Table 1:** Dopant assignment the cathodophores identified in **SI Fig. 12A** using our classification model, along with cathodophore diameter, and CL detection rate and signal-to-noise ratio (SNR) in the spectral channel of the assigned dopant. For cathodophores #38, #53, and #54, neither channel presented a local maximum, thus dopant assignment was performed based on p-values, and photon rates and SNR could not be computed.

#### 9. Impact of nonlocal excitation on single-particle CL characterization

The nonlocal excitation also impacted our single-color experiments. These experiments were an important first step to characterize the emission properties of cathodophores prior to using them for multicolor imaging. In the presence of nonlocal excitation, the CL signal collected from an isolated cathodophore was independent of its size. This is because the signal was affected by the surroundings of the cathodophore, i.e., its distance from the luminescent clusters that were excited by stray electrons (**SI Fig. 13A**). Once the nonlocal excitation was mitigated, we saw a linear dependence between the rate of CL detection from cathodophores and their size (**SI Fig. 13B**), which is consistent with theoretical predictions (see **SI Fig. 21**).

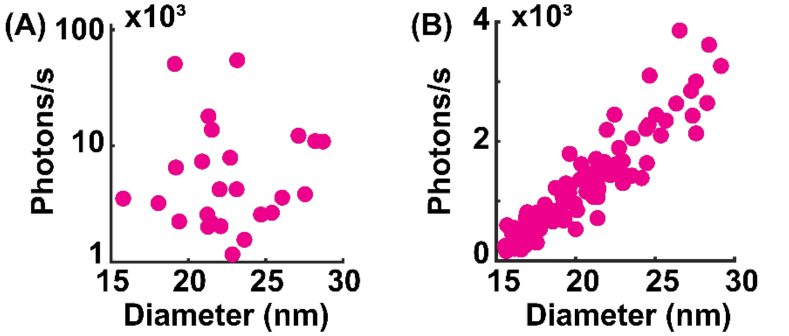

**SI Fig. 13: (A)** Rate of CL detection from single $NaHoF_4$ cathodophores in a sample with clusters of $NaHoF_4$ cathodophores. CL collected from single cathodophores was independent of their diameter (FWHM) due to nonlocal CL from the clusters. **(B)** Rate of CL detection from single $NaHoF_4$ cathodophores in a sparse sample. The rate of detection depended linearly on the diameter of cathodophores. CL was measured in the $Ho^{3+}$ color channel.

#### 10. Image analysis for single-particle characterization

Our image analysis involved:

- Drift correction
- Localization of the cathodophore in SE images
- Determination of the rate of CL detection and SNR from CL images
- Determination of the size of the cathodophore

#### Drift correction

**Need for drift correction:** A few hundred to a few thousand photons per second were collected from each cathodophore. Such a low rate of CL detection required imaging cathodophores with a beam dwell time of tens of milliseconds to achieve the required SNR for detection and classification. Although cathodophores were stable under the electron beam over multiple seconds, such a long beam dwell time led to a buildup of charge in the imaged region. As a result of this charging, the beam was deflected away from the imaged region, causing an apparent drift of the sample as shown in **SI Fig. 14**.

**Drift correction during data acquisition:** We typically acquired 50 frames, each with a pixel dwell time of 1 ms. During the acquisition of these frames, the cathodophore could drift out of the FOV. This problem was further exacerbated by the fact that we imaged a small FOV, 100 nm × 100 nm, with the cathodophore in the center. To ensure that the cathodophore remained in the FOV, we adjusted the center of the electron beam scan after each frame. To do this, we used the fact that cathodophores, composed of heavy metals, were visible in the SE channel in each frame. Drift correction was performed by determining the position of the cathodophore in the FOV and adjusting the scan region accordingly. The cathodophore's position was determined as the location of the maximum intensity SE pixel in the FOV.

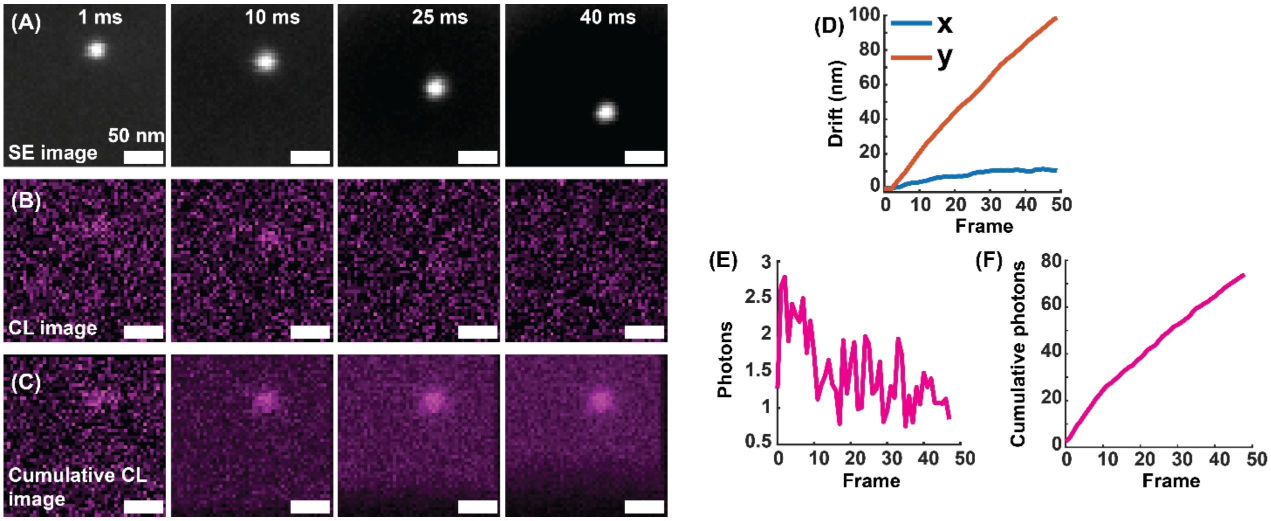

**SI Fig. 14:** **(A)** SE images of a cathodophore subjected to different total durations of electron beam exposure. The duration each pixel was exposed to the electron beam is shown in the top right corner of each image. Images were taken with a beam dwell time of 1 ms. **(B)** CL images acquired simultaneously with the SE images in (A). **(C)** Cumulative CL images obtained by summing the CL images from (B), to obtain the effective beam dwell times displayed in (A). Drift correction was applied prior to summing the images. **(D)** Displacement of the cathodophore relative to its position in the first frame. **(E)** Number of photons collected from the cathodophore as a function of the frame number. The number of photons corresponded to the amplitude of the 2D Gaussian fit to the CL images of the cathodophore as shown in (B). **(F)** Cumulative photons collected from the cathodophore, corresponding to the amplitude of the 2D Gaussian fit to the cumulative CL images of the cathodophore as shown in (C).

**Post-acquisition drift correction**: The second round of drift correction was conducted after image acquisition. Here, the position of the cathodophore was determined for all frames by fitting a 2D Gaussian function to each SE image frame. These positions were then used to translate the SE and CL frames to correct for the drift. The drift-corrected frames were then summed for further analysis.

#### Rate of CL detection

See **SI Fig. 15** for an overview of our method to calculate the rate of CL detection from cathodophores. The method involved localizing a cathodophore in SE images and determining its emission characteristics at the corresponding location in the CL images as described below:

**Localization of cathodophores in SE images:** The visibility of cathodophores in the SE channel allowed us to precisely localize them. The position of a cathodophore in its SE image was determined by fitting a 2D Gaussian function to the image.

**Number of photons:** To determine the number of photons collected from the cathodophore, a 2D Gaussian function, $S(x, y)$, was fit to the summed CL image:

$$S(x, y) = b + a \exp\left[-\frac{(\mathrm{x} - \mathrm{x}_0)^2}{2\sigma_x^2} - \frac{(y - y_0)^2}{2\sigma_y^2}\right] = b + G(x, y)$$

where $(x_0, y_0)$, is the center position of the cathodophore, $b$ is the background, $a$ is amplitude, and $\sigma_x$ and $\sigma_y$ are the standard deviations of the 2D Gaussian function. $G(x, y)$ is the 2D Gaussian fit to the cathodophore without the background. Importantly, in this equation, $x0, y0$, $\sigma_x$ and $\sigma_y$ were constrained by the 2D Gaussian fit to the SE image.

The rate of CL detection $(r)$, i.e., the number of photons collected from the cathodophore per second, was determined as

$$r = \frac{a}{\mathrm{d}t}$$

where $\mathrm{d}t$ is the pixel dwell time. The amplitude of the Gaussian function was used to determine the rate of CL detection (instead of the total number of photons in the CL image) because with our pixel size of 4–6 nm, a cathodophore was excited multiple times during image acquisition. The amplitude of the Gaussian corresponds to the most efficient excitation of the cathodophore (see **SI Fig. 15**).

#### Signal-to-noise ratio

To determine the signal-to-noise ratio (SNR) of the CL image of a cathodophore, we used the following equation[1]:

$$\mathrm{SNR} = \frac{\mathrm{Signal}}{\mathrm{Noise}} = \frac{\sum G(x, y)}{\mathrm{Noise}}.$$

The $\mathrm{Signal}$ is the total number of photons collected from the cathodophore in a CL image. This number was obtained by summing the pixels of the fitted 2D Gaussian function. Only pixels within $2\sigma$ from the center $(x_0, y_0)$ were included in the sum. Since the noise followed Poisson statistics (see **SI Fig. 22**), for the $i^{th}$ pixel:

$$\text{Pixel noise} = N_i = \sqrt{I(x, y)}\,.$$

Here, $I(x, y)$ is the CL image intensity at pixel $(x, y)$. To determine the total noise for the cathodophore, the noise was added in quadrature over the pixels within the $2\sigma$ radius of the cathodophore's center $(x_0, y_0)$:

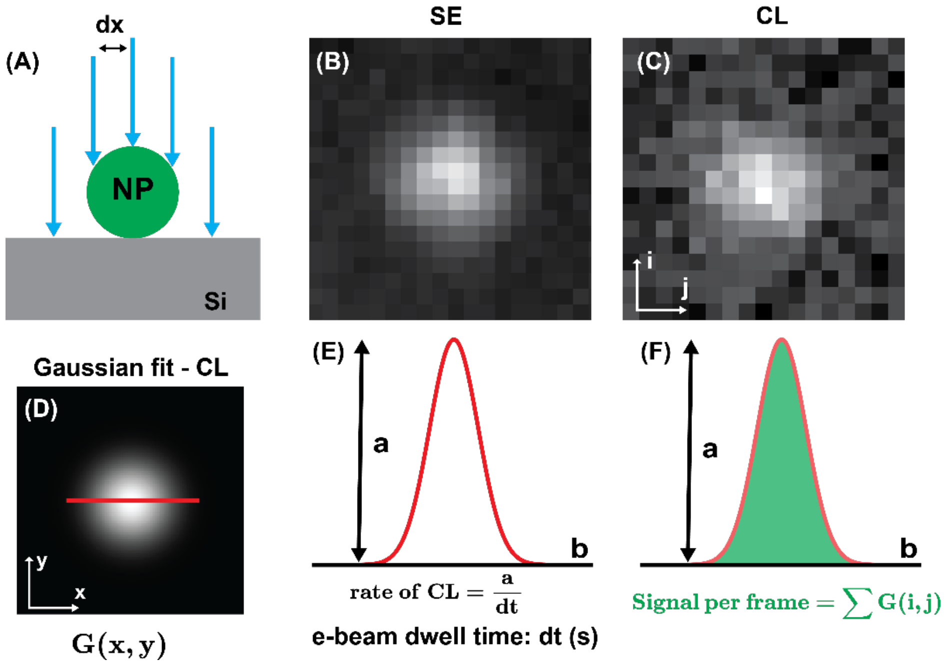

**SI Fig. 15: (A)** An illustration showing that a cathodophore was excited multiple times during image acquisition. To achieve Nyquist sampling of the smallest cathodophores, a pixel size (dx) of 4–6 nm was used in the experiments. **(B)** SE image of a cathodophore and **(C)** its corresponding CL image. **(D)** 2D Gaussian fit to the CL image from (C). **(E)** Cross-sectional profile of the 2D Gaussian fit taken along the red line shown in (D). The rate of CL was determined using the amplitude of the 2D Gaussian fit, which corresponded to the most efficient excitation of the cathodophore. **(F)** The number of photons collected from the cathodophore in a frame was determined by the area beneath the 2D Gaussian fit.

$$\text{Noise} = \sqrt{\sum N_i^2} = \sqrt{\sum I(x,y)}\,.$$

Hence,

$$\text{SNR} = \frac{\sum G(x,y)}{\sqrt{\sum I(x,y)}}\,.$$

#### Size of cathodophores

The size of cathodophores was determined from the oversampled SE images taken with the ZEISS Smart SEM software. These images had dimensions of 768 × 1024 pixels, with a pixel size of 0.2–0.3 nm. To determine the nanocrystal size, we fitted a 2D Gaussian function to the image of each cathodophore. Size was defined as the full width at half maximum (FWHM) of this fit. FWHM was calculated from the standard deviation of the Gaussian fit, as $2.355\,\sigma$. Here $\sigma$ is the average of the two standard deviations of the Gaussian fit.

It is worth noting that the oversampled images were acquired at fast scan speeds (1–3 µs pixel dwell time), three orders of magnitude faster than those used for CL imaging. Therefore, they did not suffer from the same charge buildup and subsequent drift, which would impact size determination. Additionally, more pixels per cathodophore improved Gaussian fitting.

#### 11. Spectrum of Holmium chloride

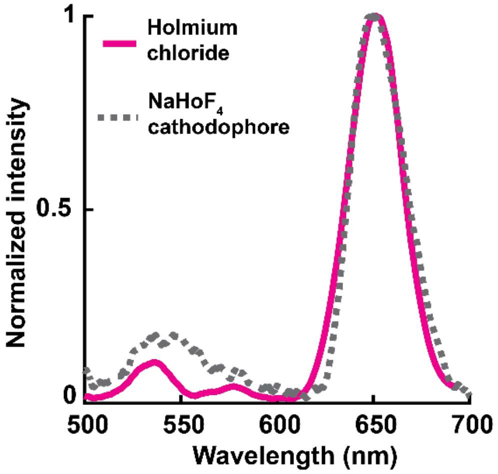

**SI Fig. 16:** Spectum of holmium chloride and its comparison to the spectrum of a single $NaHoF_4$ cathodophore.

We prepared a sample of holmium chloride on a Si wafer and measured its spectrum using our spectrometer. Results are shown in **SI Fig. 16**. The spectrum of holmium chloride matched the spectrum of $NaHoF_4$ cathodophores. A single-cathodophore spectrum was obtained by imaging the cathodophore in a sparse sample, as in **SI Fig. 28**. This result suggested that direct excitation of lanthanide ions, outside of the context of a nanocrystal, could produce CL.

#### 12. Optimizing electron beam current for CL imaging

**Rate of CL as a function of beam current:** **SI Fig. 17A** shows the relationship between the CL detection rate from single cathodophores, and the electron beam current. The beam current was measured using a Faraday cup. The figure shows that there was no significant change in CL signal from cathodophores for different beam current values. Different beam currents were obtained by changing the SEM aperture sizes, ranging from 20 μm to 120 μm, and by using “normal” and “high” current modes of the SEM.

However, with increased beam current, we also observed an increase in the apparent size of cathodophores in SE images (**SI Fig. 17B**). This increase in size could be due to the deterioration of SEM resolution at higher beam currents (e.g., due to Coulombic effects), particularly when working with large apertures, at a long working distance, and imaging through the aperture of the parabolic mirror. **SI Fig. 17C** shows that the background CL increased for higher beam currents, indicating an increase in CL emission from Si as a function of beam current.

**Electrobleaching and beam current:** We also investigated the stability of $NaHoF_4$ cathodophores for different values of beam current. Results are shown in **SI Fig. 18** for beam currents of 200 pA and 862 pA. The figure was generated by obtaining the average CL signal from cathodophores of diameters between 17–30 nm (as measured from SE images) as a function of the cumulative pixel dwell time of the electron beam. It shows that the electrobleaching increased with an increase in beam current.

In summary, although a higher beam current provided more electrons to excite lanthanide ions in the cathodophores, it also led to more electrobleaching of these ions. This effect resulted in a net decrease in the brightness of cathodophores. Based on these results we used a beam current of 160–200 pA for our CL-SEM experiments.

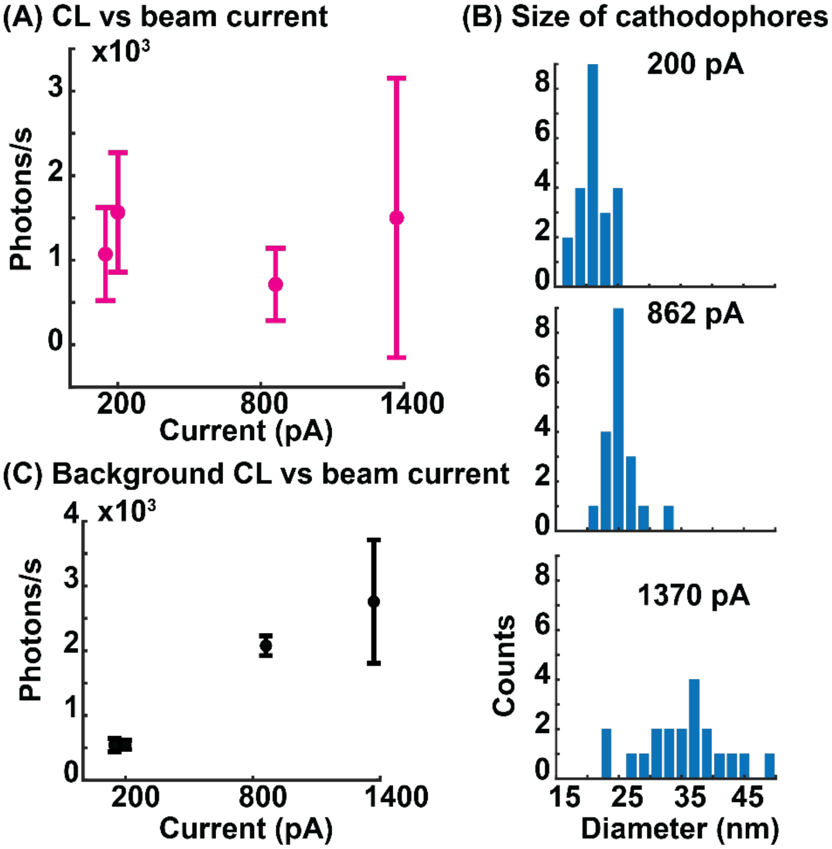

**SI Fig. 17: (A)** CL detection rate from $NaHoF_4$ cathodophores as a function of electron beam current. No significant change in the detection rate was observed. CL was measured in the $Ho^{3+}$ channel. **(B)** Histograms of cathodophore sizes at different beam currents, as determined from SE images. An increase in size at higher beam currents could be due to deterioration in the resolution of the SEM at higher currents. This deterioration may be due to poor focusing of the electron beam when working with high currents (Coulombic effects), wide SEM apertures (required for high currents), long working distances, and a beam energy of only 3 keV. **(C)** Background CL from the Si substrate as a function of beam current.

#### 13. Optimizing electron beam energy for CL imaging

**Monte Carlo simulations:** We performed Monte Carlo simulations in CASINO software[4] to determine the optimal electron beam energy to achieve efficient excitation of cathodophores. A $NaHoF_4$ nanocrystal on a Si substrate was excited with different beam energies. 5,000 trajectories were simulated for each energy, and the energy deposited in the nanocrystal was calculated. Simulations were performed for two diameters of the nanocrystal: 20 nm and 30 nm. Results are shown in **SI Fig. 19A**. The optimal excitation energy was 0.75 keV for the 20 nm diameter nanocrystal, and 1 keV for the 30 nm diameter nanocrystal. At lower beam energies, the nanocrystal was not fully excited because the beam could not penetrate the nanocrystal. On the other hand, at higher beam energies, the beam went through the nanocrystal without exciting its entire volume (**SI Fig. 19B**).

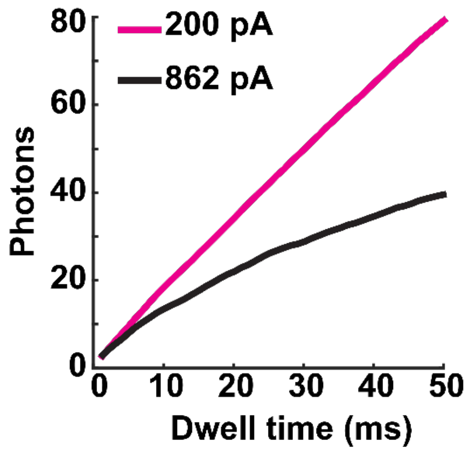

**SI Fig. 18:** Number of photons collected from $NaHoF_4$ cathodophores as a function of total electron beam dwell time, for beam currents of 200 pA and 862 pA. 50 images were taken with a dwell time of 1 ms. Images with an effective dwell time of N ms were then generated by summing N consecutive frames. The number of photons collected from a cathodophore corresponded to the amplitude of its 2D Gaussian fit in a CL image. Higher nonlinearity for 862 pA indicates increased electrobleaching. CL was measured in the $Ho^{3+}$ color channel.

**SI Fig. 19B** also shows angles at which backscattered and secondary electrons traveled in vacuum when the 20-nm nanocrystal was excited by the electron beam. At 1 keV, the electrons emerged nearly uniformly at all angles (90 degrees is antiparallel to the electron beam). In contrast, at both lower beam energies (0.25 keV in the figure) and higher beam energies (10 keV in the figure), the angles were arranged in a cosine-like distribution. This was because the high-energy beam penetrated the substrate, and the electrons scattered in directions away from the beam remained trapped within the substrate. Similarly, for lower beam energies, such electrons were trapped within the nanocrystal.

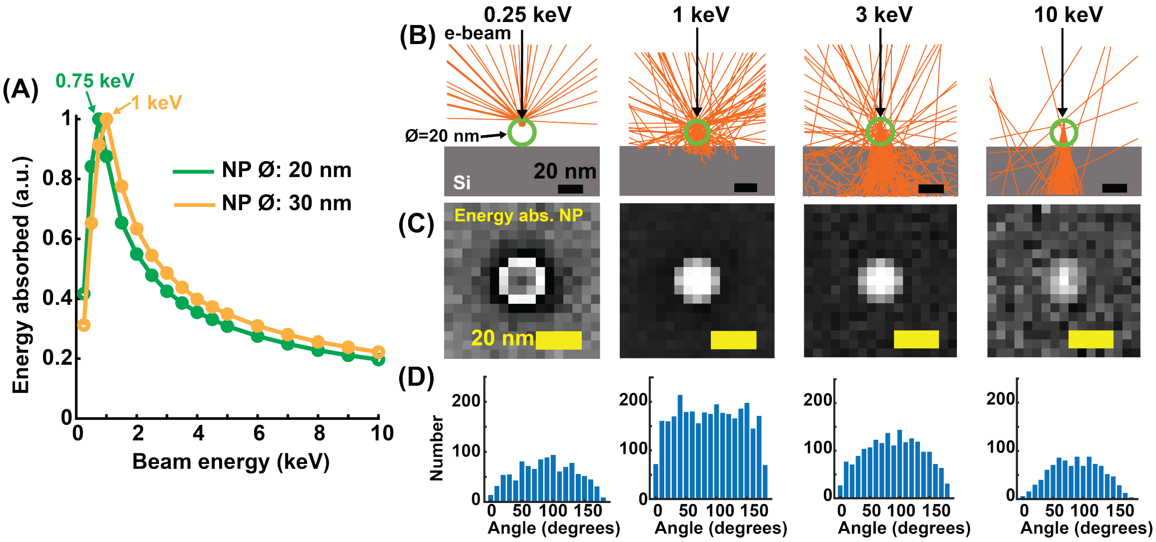

**SI Fig. 19: (A)** Monte Carlo simulations to determine the energy absorbed by $NaHoF_4$ nanocrystals of two diameters, 20 nm and 30 nm, when excited by an electron beam of different energies. 5,000 trajectories were simulated for each beam energy. **(B)** Electron trajectories for the 20-nm $NaHoF_4$ nanocrystal (green circle) excited by the electron beam of different energies. The primary electron beam is shown as a black arrow. Both backscattered and secondary electron trajectories are shown as orange lines. **(C)** Energy absorbed by the nanocrystal at each scanned position of the electron beam for the beam energies shown in (B). Each pixel of the image shows energy absorbed by the nanocrystal when the electron beam was focused at that position. **(D)** The angles at which secondary and backscattered electrons were emitted into vacuum after interaction with the sample. 90 degrees is antiparallel to the electron beam.

**Experiments to determine the optimal beam energy:** **SI Fig. 20A** shows CL from $NaHoF_4$ cathodophores of 20–25 nm diameter at different beam energies. For energies ≥3 keV, the CL signal did not change significantly. However, we observed a reduction in the CL signal for beam energies below 3 keV. This result was different from Monte Carlo simulations. The contradiction was due to the sub-optimal performance of our SEM when low beam energies were used to image at long working distances through the 500-µm-diameter aperture of the parabolic mirror. A long working distance (~6.5 mm) was needed for CL imaging because the parabolic mirror was installed between the electron gun and the sample. This suboptimal performance was also seen when we plotted the rate of CL detection from cathodophores as a function of their size (**SI Fig. 20B**). We obtained a similar CL signal for the beam energies of 3 keV and 6 keV. However, at 2.5 keV the CL signal reduced for a given cathodophore size. We attributed this reduction to the suboptimal focusing of the electron beam, which is consistent with the increase in cathodophore size shown in **SI Fig. 20B**.

Although the CL signal did not change significantly for the beam energies of 3–8 keV, the background CL from the Si substrate increased (**SI Fig. 20C**). This was because more electrons penetrated the substrate at higher beam energies. Moreover, the contrast generated by the nanocrystals decreased at high beam energies as shown in **SI Fig. 20D.** This decrease in contrast is explained by Monte Carlo simulations, which showed that, at high beam energies, many electrons passed through the cathodophores ballistically, without scattering. Based on these results, we chose 3–4 keV as the optimal beam energy for our CL-SEM experiments.

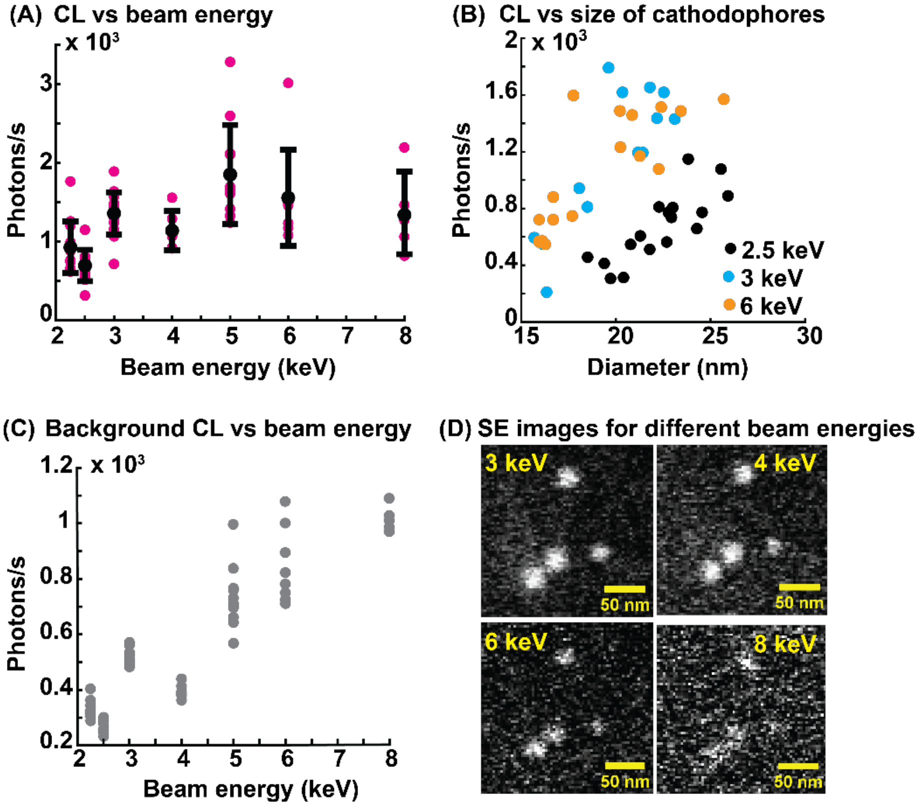

**SI Fig. 20:** **(A)** CL detection rate from $NaHoF_4$ cathodophores as a function of electron beam energy. The diameter of cathodophores was 20–25 nm. No significant change in the rate was observed for beam energies ≥3 keV. CL was measured in the $Ho^{3+}$ color channel. **(B)** CL detection rate as a function of cathodophore size for three beam energies: 2.5 keV, 3 keV, and 6 keV. The lower CL detection rate at 2.5 keV was likely due to the loss of SEM resolution when imaging at long working distances (>6 mm) using low beam energies. **(C)** Background CL from Si substrate as a function of beam energy. **(D)** SE images of the same cathodophores for different beam energies. At higher energies, the contrast was reduced because many electrons passed through the cathodophores ballistically without scattering.

**Energy absorbed by cathodophores of different sizes:** We performed Monte Carlo simulations to determine the energy absorbed by cathodophores as a function of their size under different beam energies. $NaHoF_4$ nanocrystals with diameters of 5–100 nm were simulated. The electron beam energy was varied from 0.5–10 keV. The beam diameter was set to 4 nm. 5,000 trajectories were simulated for each condition. Results are shown in **SI Fig. 21A**. The energy absorbed by the nanocrystal increased as a function of its size, up to the point where the beam interaction volume matched the size of the nanocrystal. Beyond this point, an increase in the nanocrystal diameter did not change the interaction volume inside the nanocrystal, and therefore the energy absorbed by the nanocrystal did not change (see **SI Fig. 21B**, top row). For example, for 0.5 keV and 1 keV, the energy absorbed by the nanocrystal plateaued beyond diameters of 15 nm and 40 nm respectively. However, for energies >1 keV,

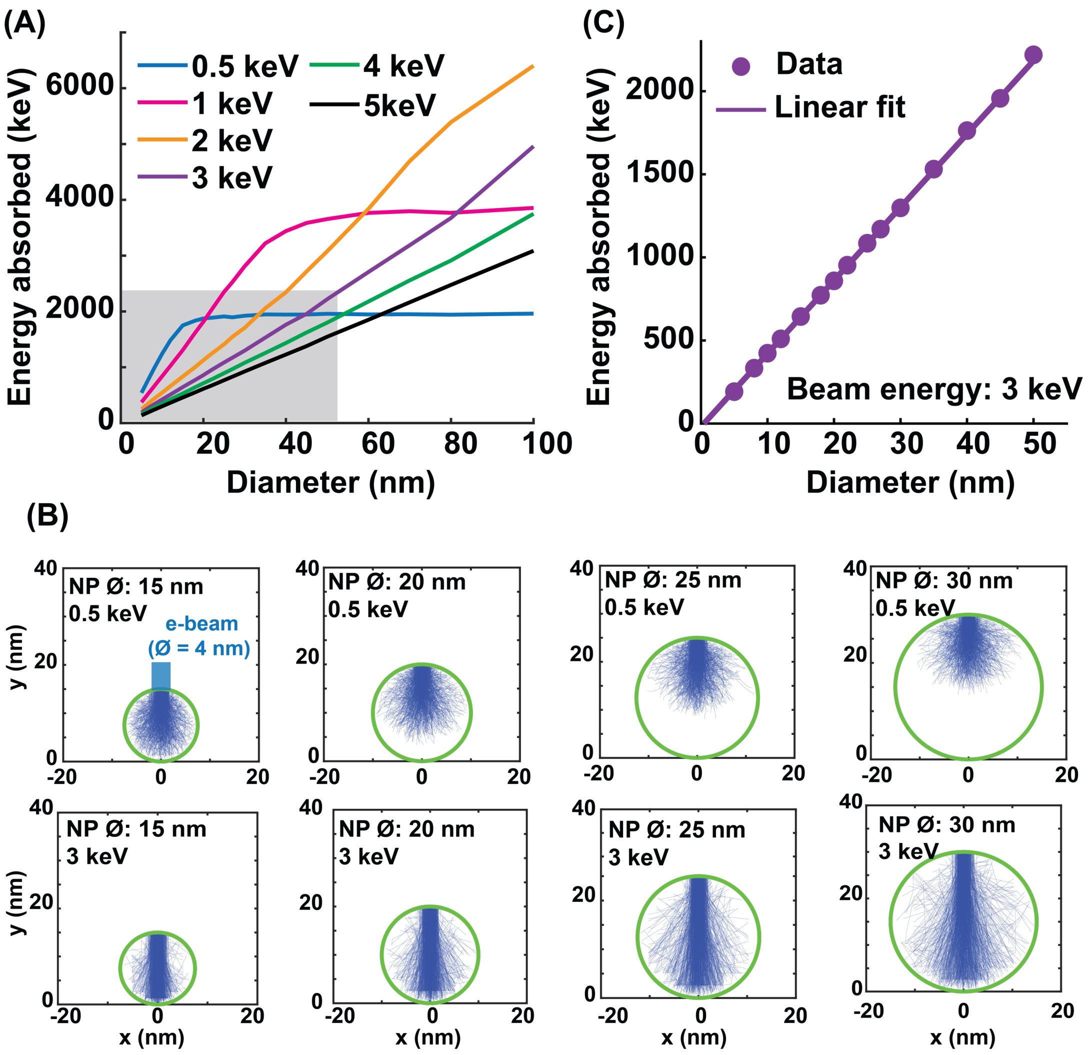

**SI Fig. 21: (A)** Monte Carlo simulations to determine the energy absorbed by $NaHoF_4$ nanocrystals of different sizes when excited by different electron beam energies. Energy absorbed by the nanocrystal increased monotonically as a function of its size until the beam interaction volume matched the size of the nanocrystal. Beyond this size, the energy absorbed by the nanocrystal did not change with an increase in the nanocrystal diameter because the interaction volume inside the nanocrystal did not change. **(B)** Electron trajectories within the nanocrystal for four different sizes (15 nm, 20 nm, 25 nm, and 30 nm), for beam energies of 0.5 keV (top row) and 3 keV (bottom row). **(C)** Focusing on the shaded region in (A) for 3 keV: energy absorbed by nanocrystals of different sizes when excited by a 3 keV electron beam. The data (circles) fit well with a straight line, indicating a linear relationship between the energy absorbed by nanocrystals as a function of size at 3 keV excitation beam energy.

which optimally excited nanocrystals with diameters >100 nm, a monotonic increase in the energy absorbed by the nanocrystal was observed for all the simulated sizes.

When the beam energy was significantly higher than the optimal energy, we observed a linear relationship between the energy absorbed by the nanocrystal and its size. This linear trend was observed because at such energies, the entire volume of the nanocrystal was not excited. Instead, the overlap between the electron interaction volume and the volume of the nanocrystal could be approximated by a vertical cylinder with a diameter corresponding to the size of the primary electron beam and the length given by the $z$-dimension (height) of the nanocrystal. So, the height of the nanocrystal primarily determined the energy absorbed by the nanocrystal (see **SI Fig. 21B**, bottom row).

**SI Fig. 21C** shows results of exciting nanocrystals with diameters ranging from 5–50 nm, with a 3 keV beam. We observed a linear increase in the energy absorbed by the nanocrystal as a function of its size. Note that we used a beam energy of 3 keV in our CL experiments. Assuming that CL emission is proportional to the energy absorbed by the nanocrystal (true for fluorescence in the linear regime within a factor given by the quantum yield), we expected a linear increase in the CL from cathodophores as a function of their size. **Fig. 5A** experimentally confirmed that the CL increased linearly as a function of cathodophore size.

#### 14. Background from Si substrate

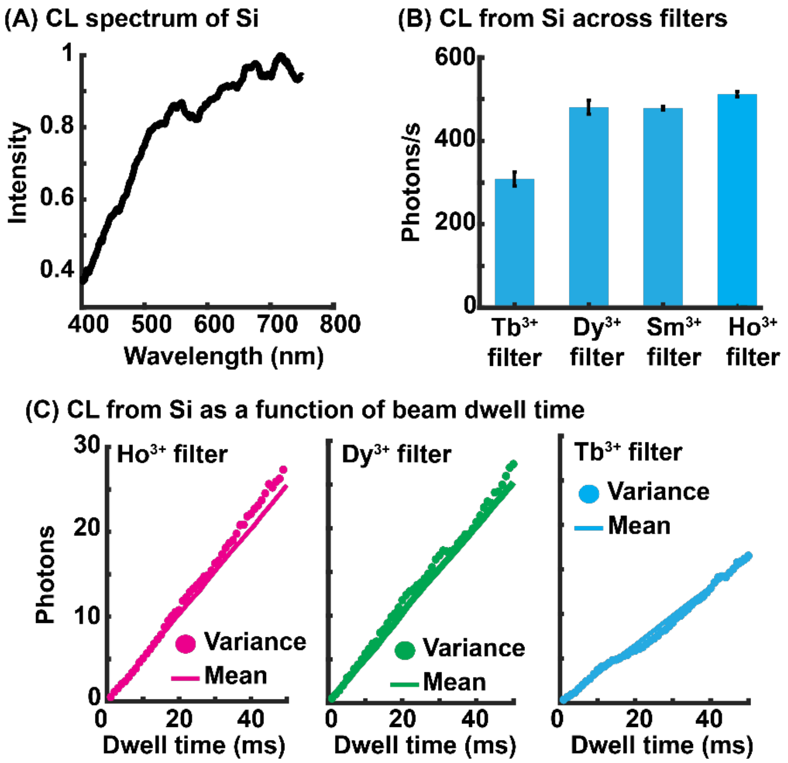

**SI Fig. 22: (A)** CL spectrum of Si obtained using our spectrometer (**SI Fig. 5**). A 100 nm × 100 nm region of Si was repeatedly scanned to collect the spectrum. **(B)** CL signal from Si captured across the spectral filters matched to the emission peaks of $NaHoF_4$, $NaDyF_4$, $NaSmF_4$, and $NaTbF_4$. Error bars correspond to the standard deviation in the average CL signal collected from ten different regions of the Si wafer. CL was collected with an effective beam dwell time of 50 ms (50 frames with 1 ms dwell time each). **(C)** CL signal from Si as a function of the effective beam dwell time. Mean and variance of the signal were equal for all dwell times, suggesting a Poisson distribution of noise.

**<u>CL signal from Si substrate:</u>** We used Si wafers as a substrate for CL imaging due to its flatness, conductivity, and low CL. The Si substrate had a wide CL emission spectrum and led to ~500 detected photons/s on average in $Ho^{3+}$, $Dy^{3+}$ and $Sm^{3+}$ color channels, and ~300 detected photons/s in the $Tb^{3+}$ color channel as shown in **SI Fig. 22A&B**. This CL from Si appeared as background when cathodophores were imaged.

CL from Si increased linearly as a function of electron beam dwell time. Additionally, the mean of the CL signal matched its variance for all dwell times, which was consistent with Poisson statistics of the noise (**SI Fig. 22C**). Mean and variance were calculated from pixels in CL images of the Si substrate acquired at different dwell times. For our typical dwell time of 50 ms (50 frames captured at 1 ms dwell time, and then summed) we got an average of ~25 photons per pixel in $Ho^{3+}$, $Dy^{3+}$, and $Sm^{3+}$ color channels and ~15 photons per pixel in the $Tb^{3+}$ color channel.

**<u>Simulations to analyze the detection limit:</u>** We conducted simulations to determine the minimum CL emission rate required for the detection of CL from cathodophores. CL images were simulated for the typical electron beam dwell time of 50 ms. The background was set to 25 counts per pixel (500 photon/s/pixel) with a Poisson distribution to match the CL from Si across our band-pass filters, as discussed in the previous section. CL signal from cathodophores was modeled as a 2D Gaussian function. Cathodophores with varying levels of CL signal were simulated by changing the amplitude of the simulated 2D Gaussian function. Simulated images were analyzed using the image processing protocol discussed in **SI Fig. 15**. As expected, the SNR of CL images increased with an increase in the number of photons collected from cathodophores as shown in **SI Fig. 23A&B**. **SI Fig. 23E** shows representative simulated images with different SNRs.

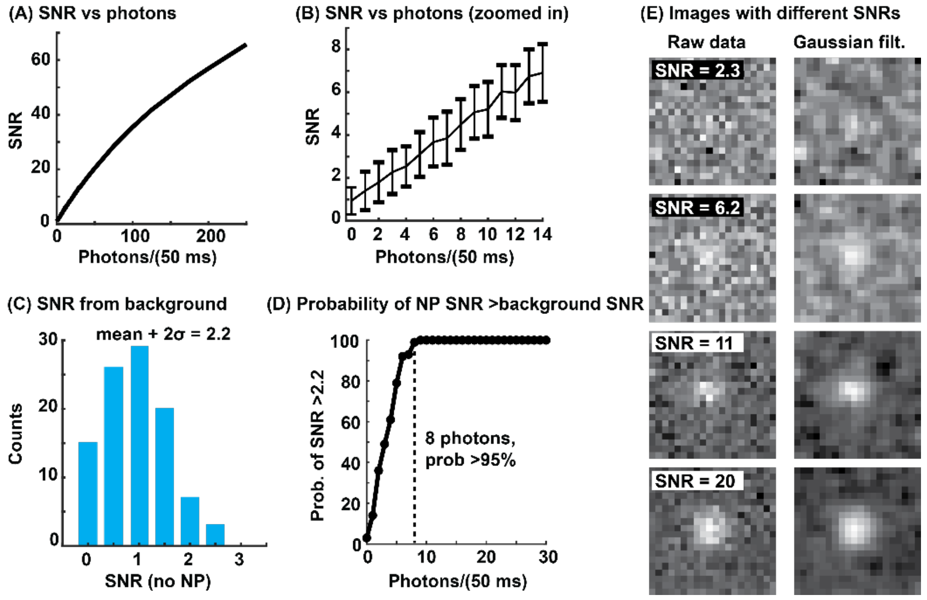

**SI Fig. 23: (A)** SNR of simulated CL images as a function of the number of photons collected from nanocrystals, given a beam dwell time of 50 ms. **(B)** A magnified region of the plot in (A). Error bars show the standard deviation for SNR across 100 frames. Images were generated with a Poisson background with an average of 25 photons to match the CL from Si at our typical beam dwell time of 50 ms. Nanocrystals were simulated as a 2D Gaussian function above the background. Nanocrystals with different numbers of photons were generated by changing the amplitude of the 2D Gaussian. 100 images were simulated for each photon count. **(C)** Histogram of SNRs obtained by fitting a 2D Gaussian function to the background across 100 simulated images. **(D)** Probability of obtaining an SNR >2.2 for different numbers of photons collected from nanocrystals. The number 2.2 corresponds to the $(mean + 2\sigma)$ of the SNR distribution in (C). **(E)** Simulated images of nanocrystals at different SNRs and their respective Gaussian filtered images ($\sigma = 0.7\ pixels$).

To determine the detection limit, we analyzed images without cathodophores. In this case, the simulated images contained only the background from the Si substrate. Analysis of these images corresponded to fitting a 2D Gaussian to the background, where the amplitude of the fitted Gaussian was determined by the background noise (i.e., Poisson noise from Si substrate). The histogram in **SI Fig. 23C** shows the distribution of SNRs for images without the cathodophores. The distribution had a $mean + 2\sigma$ value of 2.2, which we used as the lower limit for the detection of cathodophores. Based on this criterion, we needed at least 8 photons (160 photons/s) from a cathodophore to detect it with 95% confidence. For CL detection rates exceeding 160 photons/s, we could detect the cathodophore in at least 95 out of 100 iterations. **Fig. 5A** shows that the detection rate of 160 photons/s corresponds to a diameter of ~15 nm for $Ho^{3+}$, $Dy^{3+}$ and $Tb^{3+}$ cathodophores, which was also the $x$-intercept of the linear fits shown in the figure. These results confirmed that the background from the Si substrate dictated the minimal photon count necessary for the statistically significant detection of CL from cathodophores.

#### 15. NaGdF$_4$ nanocrystals as a control sample

**Significance of using NaGdF$_4$ nanocrystals as a control:** Background CL from Si substrate determined the minimum brightness of cathodophores required for their detection. However, detection of a cathodophore in a specific spectral channel could also be due to nonlocal excitations. We took steps to mitigate these excitations, which included optimizing the sample preparation protocol and optically isolating the emission path inside the SEM with a lens tube.

Nevertheless, some nonlocal CL remained, especially for larger cathodophores, and it was important to ensure that it did not impact our analysis for the classification of cathodophores across all sizes. One potential cause of this nonlocal signal could be the enhanced CL from the Si substrate: excitation of cathodophores led to a higher number of secondary electrons, which could interact with the substrate. And the Si substrate, as we learned from **SI Fig. 22C**, showed brighter luminescence at larger beam currents. Additionally, even with the lens tube optically isolating the emission path inside the SEM, some nonlocal CL could potentially reach the PMTs.

As discussed earlier, cathodophores were composed of high-$Z$ elements and produced more secondary electrons than the Si substrate. Therefore, it was important for the control to match the characteristics (shape, size, and atomic number ($Z$)) of the cathodophores, while being non-cathodoluminescent to ensure that it only appeared luminescent due to nonlocal excitation. $NaGdF_4$ particles met these criteria, so we used them as a control instead of Si.

**SNR of NaGdF$_4$ nanocrystals:** $NaGdF_4$ nanocrystals were imaged across different color filters. The SNR was calculated for these images and results are shown in **Nevertheless, some nonlocal** CL remained, especially for larger cathodophores, and it was important to ensure that it did not impact our analysis for the classification of cathodophores across all sizes. One potential cause of this nonlocal signal could be the enhanced CL from the Si substrate: excitation of cathodophores led to a higher number of secondary electrons, which could interact with the substrate. And the Si substrate, as we learned from **SI Fig. 22C,** showed brighter luminescence at larger beam currents. Additionally, even with the lens tube optically isolating the emission path inside the SEM, some nonlocal CL could potentially reach the PMTs. **A**. The figure shows that the SNR from $NaGdF_4$ nanocrystals increased with an increase in their size, which could be a result of increased nonlocal excitation due to a higher number of secondary electrons emitted from larger nanocrystals. For bona fide cathodophores (e.g., $NaHoF_4$) to be detected in their respective channels, they had to be brighter than the $NaGdF_4$ nanocrystals. This was indeed the case for all dopants ($Ho^{3+}$, $Dy^{3+}$, $Sm^{3+}$ and $Tb^{3+}$) as shown in **Nevertheless,**

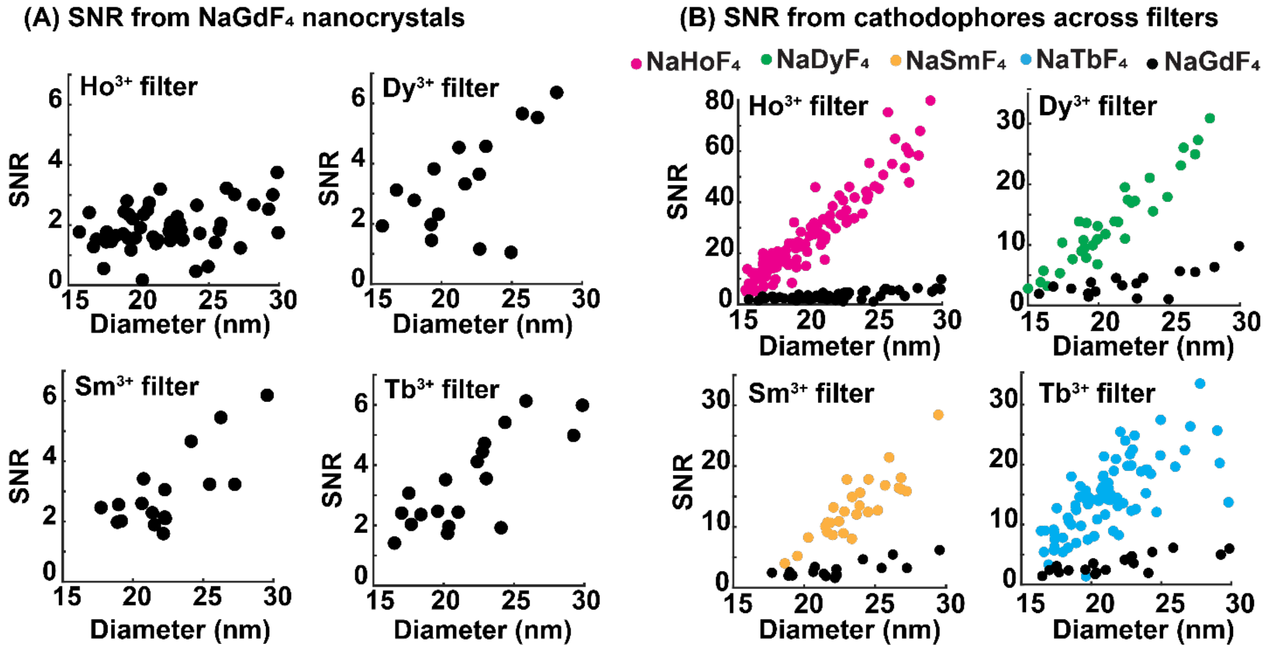

**SI Fig. 24: (A)** SNR from control, non-emitting $NaGdF_4$ nanocrystals as a function of their size. Each subpanel shows SNR in one of the four spectral filters matched to the emission peaks of $NaHoF_4$, $NaDyF_4$, $NaSmF_4$, and $NaTbF_4$ cathodophores. SNR was determined by fitting a 2D Gaussian function to the location of a nanocrystal in its CL image. The location of the nanocrystal was determined from the SE image of the same region. **(B)** SNR from $NaHoF_4$, $NaDyF_4$, $NaSmF_4$, and $NaTbF_4$ cathodophores across the filters matched to their respective emission peaks. SNR from $NaGdF_4$ is also shown (in black) for comparison.

**some nonlocal** CL remained, especially for larger cathodophores, and it was important to ensure that it did not impact our analysis for the classification of cathodophores across all sizes. One potential cause of this nonlocal signal could be the enhanced CL from the Si substrate: excitation of cathodophores led to a higher number of secondary electrons, which could interact with the substrate. And the Si substrate, as we learned from **SI Fig. 22C,** showed brighter luminescence at larger beam currents. Additionally, even with the lens tube optically isolating the emission path inside the SEM, some nonlocal CL could potentially reach the PMTs. **B**.

**<u>Smallest cathodophores classified in our CL-SEM experiments:</u>** Next, we determined the minimum size of cathodophores that could be classified in our CL-SEM experiments. To this end, we compared the SNR from CL images of $NaHoF_4$, $NaDyF_4$, and $NaTbF_4$ cathodophores of different sizes to the SNR from $NaGdF_4$ particles of the same size. Results are shown in **SI Fig. 25**. Cathodophores were classified as containing a specific dopant if the SNR of their CL images was higher than $mean + 2\sigma$ of the SNR obtained from $NaGdF_4$ nanocrystals of the same size. Each dopant was analyzed in the CL channel matched to its emission peak.

We found that we could classify $NaHoF_4$, $NaDyF_4$ and $NaTbF_4$ down to 15 nm diameter. $NaHoF_4$ could be detected with 100% probability across all sizes. For $NaDyF_4$, a 100% detection probability was achieved for cathodophores exceeding 16 nm. **SI Fig. 25C**, **F&I** show the cathodophores of different sizes used in the analysis. It is important to note that 15 nm was the smallest diameter of nanocrystals visible in our SE images. This could be because at 3 keV of beam energy, cathodophores smaller than 15 nm did not produce enough secondary electrons to be detected.

**<u>Frames required for detection:</u>** We also determined the number of frames (1 ms pixel dwell time per frame) required for the classification of cathodophores, i.e., the number of frames required to achieve the SNR essential for classification. Results are shown in **SI Fig. 26.** The figure shows that our typical effective dwell time of 50 ms was sufficient to detect cathodophores across all sizes.

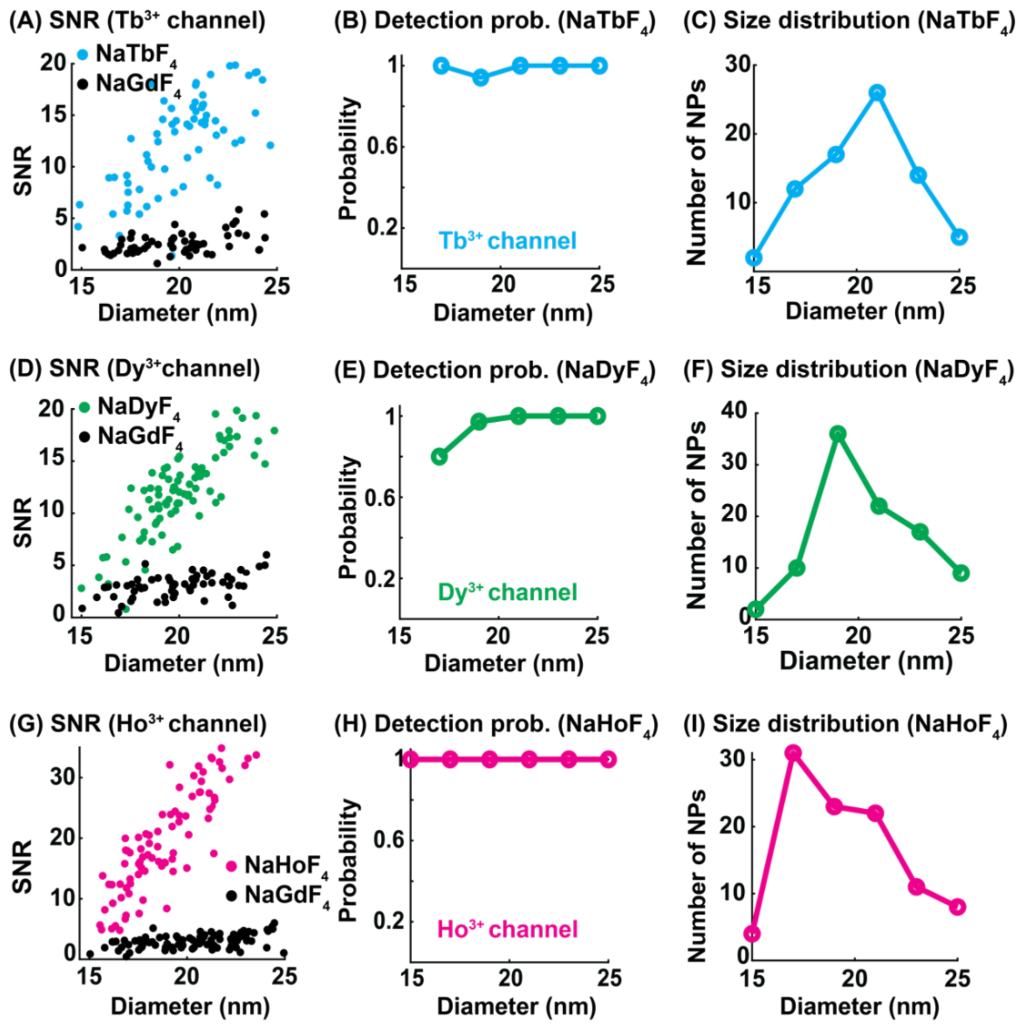

**SI Fig. 25: (A)** SNR from $NaTbF_4$ and $NaGdF_4$ nanocrystals in the $Tb^{3+}$ color channel as a function of their size. **(B)** Probability of detecting $NaTbF_4$ cathodophores in the $Tb^{3+}$ color channel as a function of their size. **(C)** Size distribution of $NaTbF_4$ cathodophores used in the analysis in (A) and (B). **(D)** SNR from $NaDyF_4$ and $NaGdF_4$ nanocrystals in the $Dy^{3+}$ color channel as a function of size. **(E)** Probability of detecting $NaDyF_4$ cathodophores in the $Dy^{3+}$ color channel as a function of their size. **(F)** Size distribution of $NaDyF_4$ cathodophores used in the analysis in (D) and (E). **(G)** SNR from $NaHoF_4$ and $NaGdF_4$ nanocrystals in the $Ho^{3+}$ color channel as a function of their size. **(H)** Probability of detecting $NaHoF_4$ cathodophores in the $Ho^{3+}$ color channel as a function of their size. **(I)** Size distribution of $NaHoF_4$ cathodophores used in the analysis in (G) and (H). (B, E, H) The probability of detecting cathodophores was calculated from the fraction of cathodophores with an SNR higher than the $(mean + 2\sigma)$ of the SNR distribution from $NaGdF_4$ nanocrystals of the same size and in the corresponding color channel. (C, F, I) Diameter was measured as the FWHM of a cathodophore in its SE image.

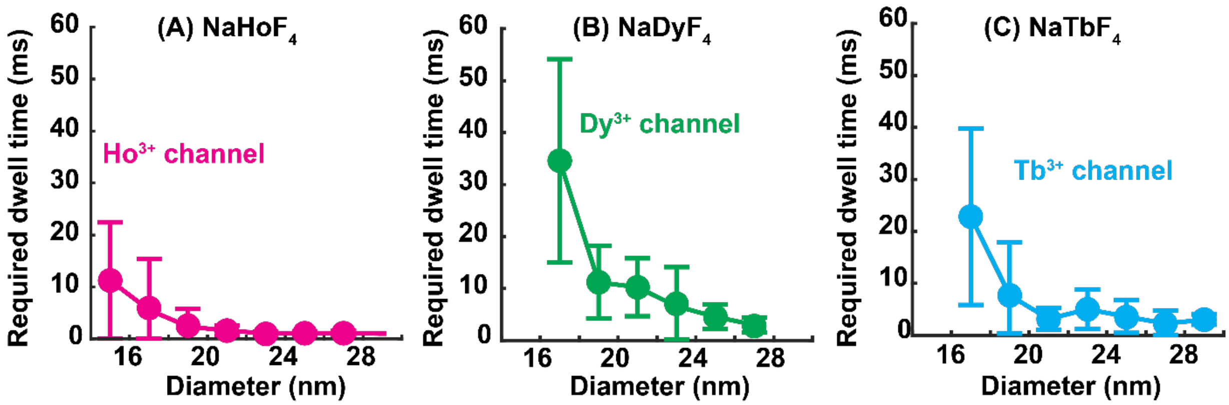

**SI Fig. 26: (A)** Effective beam dwell time required to detect $NaHoF_4$ cathodophores in the $Ho^{3+}$ color channel. **(B)** Effective beam dwell time required to detect $NaDyF_4$ cathodophores in the $Dy^{3+}$ color channel. **(C)** Effective beam dwell time required to detect $NaTbF_4$ cathodophores in the $Tb^{3+}$ color channel. (A–C) 50 frames were acquired with a beam dwell time of 1 ms. Images with an effective dwell time of N ms were generated by summing N consecutive frames. Cathodophores were detected if their SNR was higher than $(mean + 2\sigma)$ of the SNR distribution from same-sized $NaGdF_4$ nanocrystals, acquired using the same dwell time and in the corresponding color channels.

#### 16. CL spectra

**Acquisition of single-particle spectra:** To obtain single-cathodophore spectra, we prepared a sparse sample of cathodophores containing a specific dopant. CL from the cathodophore was collected by repeatedly scanning the electron beam over it. The CL emitted by the cathodophore was collected by the parabolic mirror and projected onto the spectrometer: the light was focused onto the camera through the diffraction grating (see **SI Fig. 5**).

**Analysis:** Spectra were extracted from images taken with the sCMOS camera. The spectral information resided in the first diffraction order of the grating. Spectra were obtained by plotting the intensity of pixels in the first diffraction order as a function of distance from the center of the zeroth order. This distance was then converted to wavelength using a calibration sample with a known spectrum. A moving average filter was applied to the distance to remove high-frequency noise from the spectra.

**SI Fig. 27** shows spectra of two $NaHoF_4$ cathodophores. **SI Fig. 27A&B** show SE images of the cathodophores along with their spectra in **SI Fig. 27C&D** respectively. For both cathodophores, the spectrum of $Ho^{3+}$ ions was observed superimposed on the spectrum of Si. The larger cathodophore emitted a higher signal above the background. The figure also shows that the spectra of $NaHoF_4$ cathodophores diminished over time due to electrobleaching.

To extract the spectrum of cathodophores from the measured spectrum, we assumed that the latter was a linear combination of the spectra of lanthanide ions and Si. Assuming $A_{Si}$ is the spectrum of Si substrate and $A_{NP}$ is the spectrum of the cathodophore (obtained from an ensemble of cathodophores), the measured spectrum, $I$, can be written as:

$$I = a_{Si}A_{Si} + a_{NP}A_{NP}$$

or

$$I = A \times a$$

where $A = [A_{Si}\ A_{NP}]$ and $a = [a_{Si}\ a_{NP}]^T$. $a_{Si}$ and $a_{NP}$ represent the respective contributions from

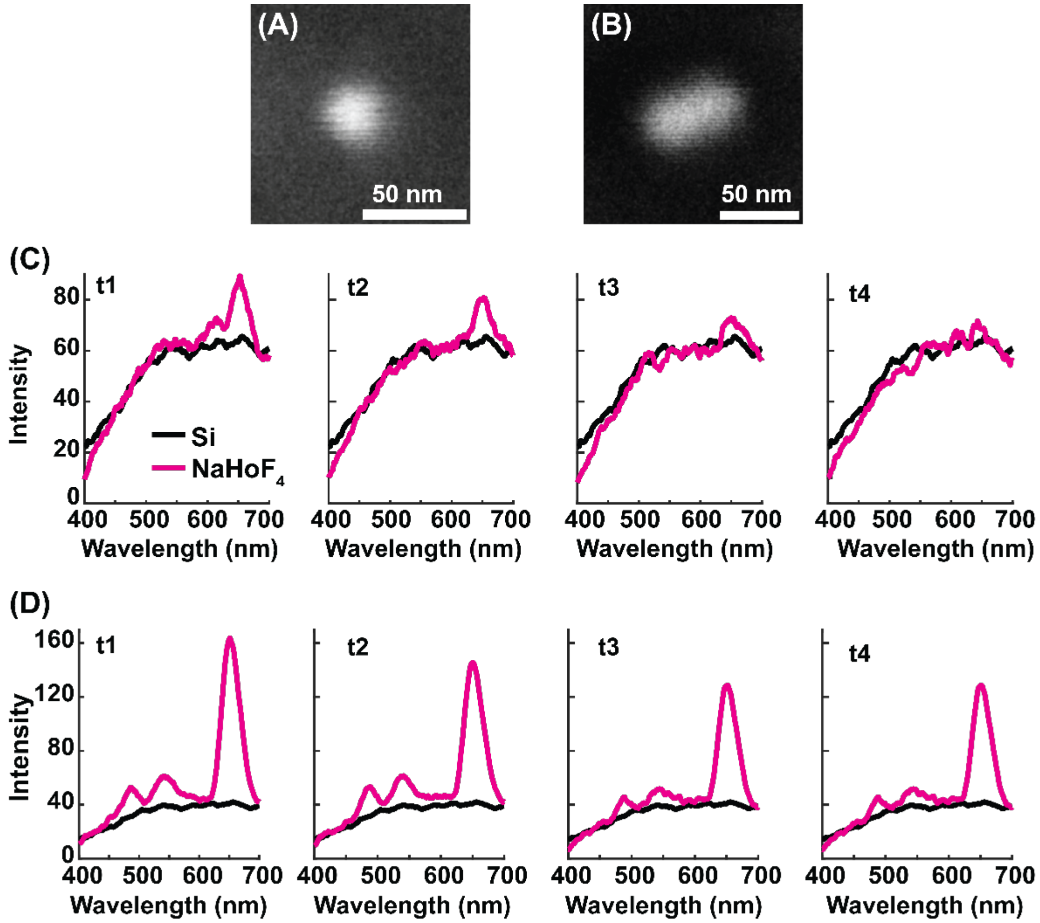

**SI Fig. 27: (A, B)** SE images of two $NaHoF_4$ cathodophores of different sizes. **(C, D)** Spectra of the cathodophores shown in A and B respectively. Spectra were obtained by repeatedly scanning the electron beam over a 100 nm × 100 nm region around the cathophores. Subpanels show spectra obtained at different time points, t1–t4. Signal from the cathodophores decayed over time due to electrobleaching. The Si substrate showed minimal electrobleaching.

Si and the cathodophore to the measured spectrum. The equation was then solved for the coefficient matrix, $a$. We solved this equation in Matlab using the 'mldivide()' function.

Finally, the spectrum of the cathodophore, $I_{NP}$, was calculated as follows:

$$I_{NP} = I - a_{Si}A_{Si}\ .$$

In other words, the spectrum of the cathodophore was obtained by subtracting the Si spectrum from the experimentally obtained spectrum.

**Example spectra:** **Fig. 5D** shows representative spectra of single cathodophores. In **SI Fig. 28** more examples of single-particle spectra are provided for $NaHoF_4$, $NaDyF_4$, $NaTbF_4$, and $NaSmF_4$ cathodophores. The figure shows that the spectra of single cathodophores matched their ensemble spectra, confirming the spectral stability of cathodophores at the single-particle level. This spectral stability allowed us to specifically select filters matched precisely to the primary emission peaks of the cathodophores.

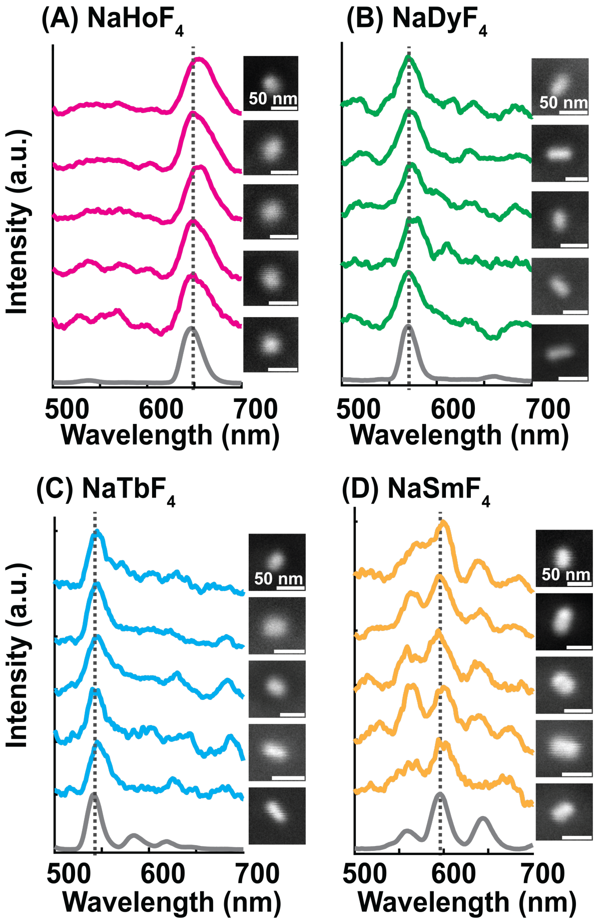

**SI Fig. 28: (A–D)** Single-particle spectra of $NaHoF_4$, $NaDyF_4$, $NaTbF_4$, and $NaSmF_4$ cathodophores respectively. Spectra were acquired using our custom spectrometer (**SI Fig. 5**) by continously scanning a 100 nm × 100 nm region around the cathodophores. Ensemble spectra of cathodophores are also shown (gray lines). Single-particle spectra of cathodophores matched their ensemble spectra. Dotted lines indicate the positions of the primary emission peaks of the cathodophores, for which the spectral band-pass filters were selected.

#### 17. Electrobleaching

**SI Fig. 29** shows CL collected from $NaHoF_4$ cathodophores in a CL image. CL from three different size ranges is shown. The electron beam dwell time was 5 ms. The CL signal in subpanels A–C correspond to the total CL signal collected from cathodophores within a frame. It was determined by calculating the area under the 2D Gaussian fit to the CL image of the cathodophore in the $Ho^{3+}$ color channel. The figure shows that the detected CL from cathodophores decreased over time, which was indicative of electrobleaching. For each size range, CL from $NaGdF_4$ nanocrystals of the same size is also shown. Notably, despite the electrobleaching, CL from $NaHoF_4$ cathodophores was higher than that of $NaGdF_4$ for all sizes and dwell times.

**SI Fig. 30** shows CL collected from $NaDyF_4$ cathodophores of three different sizes. Unlike $NaHoF_4$, $NaDyF_4$ displayed minimal electrobleaching, even for smaller, sub-20-nm cathodophores. Since $NaDyF_4$ cathodophores emitted fewer photons than $NaHoF_4$, their CL signal in some frames was comparable to the CL from the control, $NaGdF_4$. However, CL from $NaDyF_4$ for a dwell time of 50 ms (**SI Fig. 30D–F**) was higher than $NaGdF_4$, enabling their classification down to ~16 nm. These results show the effect of the dopant type on electrobleaching. However, the microscopic mechanism behind this difference remains to be understood.

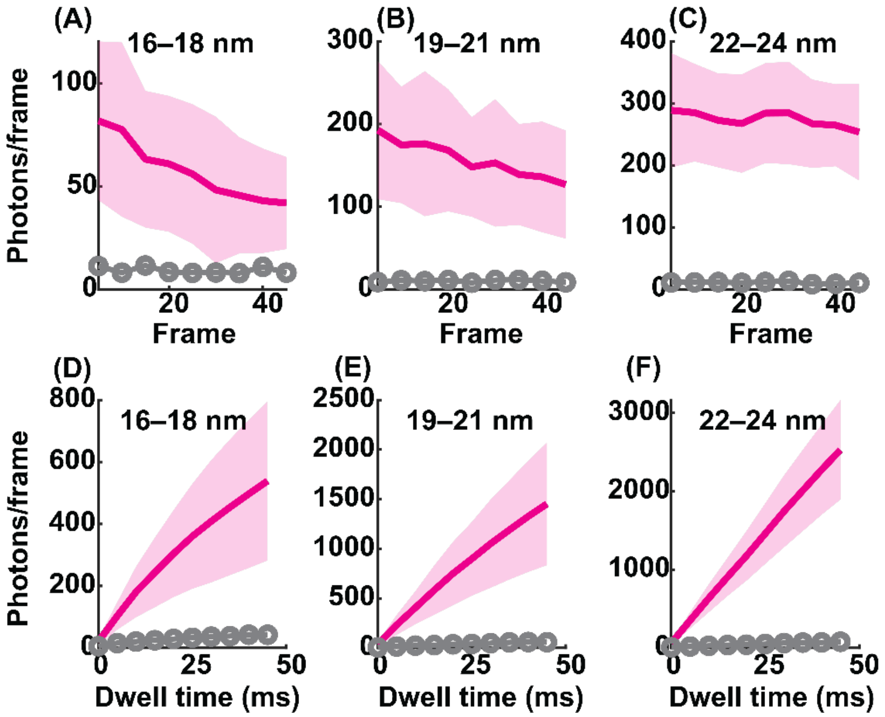

**SI Fig. 29: (A–C)** Photons/frame collected from $NaHoF_4$ in a CL image for three different cathodophore diameters. Photons/frame obtained from $NaGdF_4$ are also shown for a comparison (gray). Images had an effective beam dwell time of 5 ms. **(D–F)** Cumulative photons collected from $NaHoF_4$ cathodophores in a CL image for different beam dwell times. Images were obtained by summing photons/frame shown in (A–C). (A–F) Photons/frame correspond to the integrated area under the 2D Gaussian fit to the CL image of cathodophores. CL was measured in the $Ho^{3+}$ color channel. In (A–F), solid lines and shaded regions correspond to the mean and standard deviation of the data.

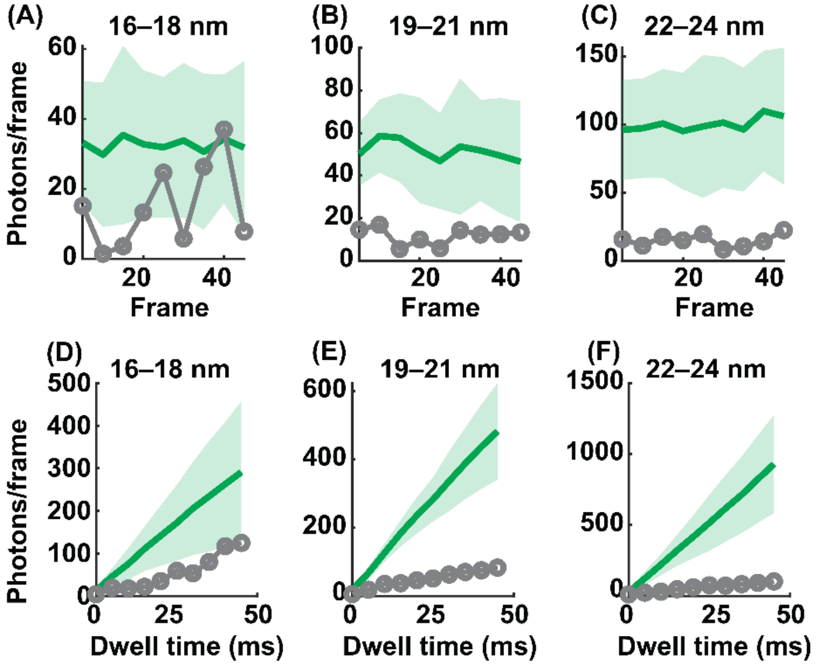

**SI Fig. 30: (A–C)** Photons/frame collected from $NaDyF_4$ in a CL image for three different cathodophore diameters. Photons/frame obtained from $NaGdF_4$ are also shown for a comparison (gray). Images had an effective beam dwell time of 5 ms. **(D–F)** Cumulative photons collected from $NaDyF_4$ cathodophores in a CL image for different beam dwell times. Images were obtained by summing photons/frame shown in (A–C). (A–F) Photons/frame correspond to the integrated area under the 2D Gaussian fit to the CL image of cathodophores. CL was measured in the $Dy^{3+}$ color channel. In (A–F), solid lines and shaded regions correspond to the mean and standard deviation of the data.

#### 18. Imaging single cathodophores on the surface of a mammalian cell

**Sample preparation for mammalian cells:** Cultured cells were prepared for CL imaging via osmium tetroxide treatment and hexamethyldisilazane (HMDS) drying based on previously reported protocols[5,6]. HEK293T cells were cultured in 50/50 DMEM/F-12 media with 10% fetal bovine serum and 1% penicillin/streptomycin. Cells were dissociated using 0.25% trypsin in PBS, then seeded on plasma-cleaned Si wafers. After adhering to the Si wafers overnight, cells were washed twice with PBS and fixed with 4% formaldehyde, 0.2% glutaraldehyde in PBS, pH 7.4 for 15 min with gentle agitation. Cells were washed three times with PBS and incubated in 1% osmium tetroxide ($OsO_4$) in PBS for 10 min with gentle agitation. Then, cells were washed with MilliQ water for 2 hours with gentle agitation, replacing water every 15 min. Cells were gradually transitioned into ethanol by increasing from 10% ethanol to 100% ethanol using steps of 10%, with 5 min of gentle agitation between each step. Upon reaching 100% ethanol, cells were gently agitated twice for 20 min with fresh ethanol each time. Next, cells were gradually transitioned into HMDS by increasing from 25% HMDS in ethanol to 100% HMDS using steps of 25%, with 5 min of gentle agitation between each step. When 100% HMDS was reached, cells were gently agitated twice for 20 min with fresh HMDS each time. Finally, the majority of HMDS volume was removed from the cells to leave a thin layer of HMDS, and cells were air dried. If appropriate, 5 µL of washed, hexane-suspended cathodophores was drop-cast onto the cells. Finally, cells were sputter-coated with a 2.5-nm-thick layer of 80:20 Pt:Pd to prevent charging during CL-SEM imaging.

**Spectra of cells without $OsO_4$:**

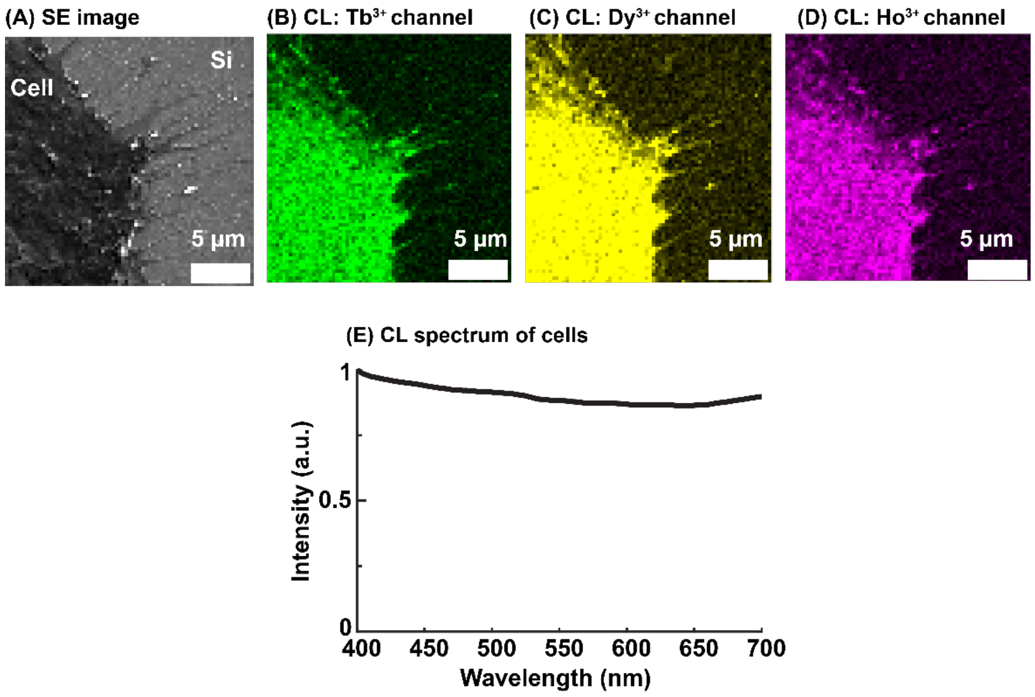

**SI Fig. 31: (A)** SE image of a HEK293T cell on a Si wafer. **(B, C, D)** Auto-CL from the cell in (A) in the $Tb^{3+}$, $Dy^{3+}$, and $Ho^{3+}$ color channels respectively. **(E)** Quantum-efficiency-corrected CL spectrum of the auto-CL of HEK293T cells.

#### 19. Nonlocal excitation from the SEM chamber

In our initial single-particle experiments, we observed CL from samples containing only non-emitting $NaGdF_4$ nanocrystals. This was surprising because these particles only emitted in the ultraviolet spectral range[7,8], which is filtered out by our emission-path optics and is not detected by the PMTs. The CL from these nanocrystals increased at shorter wavelengths **(SI Fig. 32A&C)**.

Investigating this phenomenon led us to conclude that this luminescence originated from the Everhart-Thornley detector that detects secondary electrons using a luminescent scintillator **(SI Fig. 32)**. It is this luminescence that was detected when an $NaGdF_4$ nanocrystal was excited

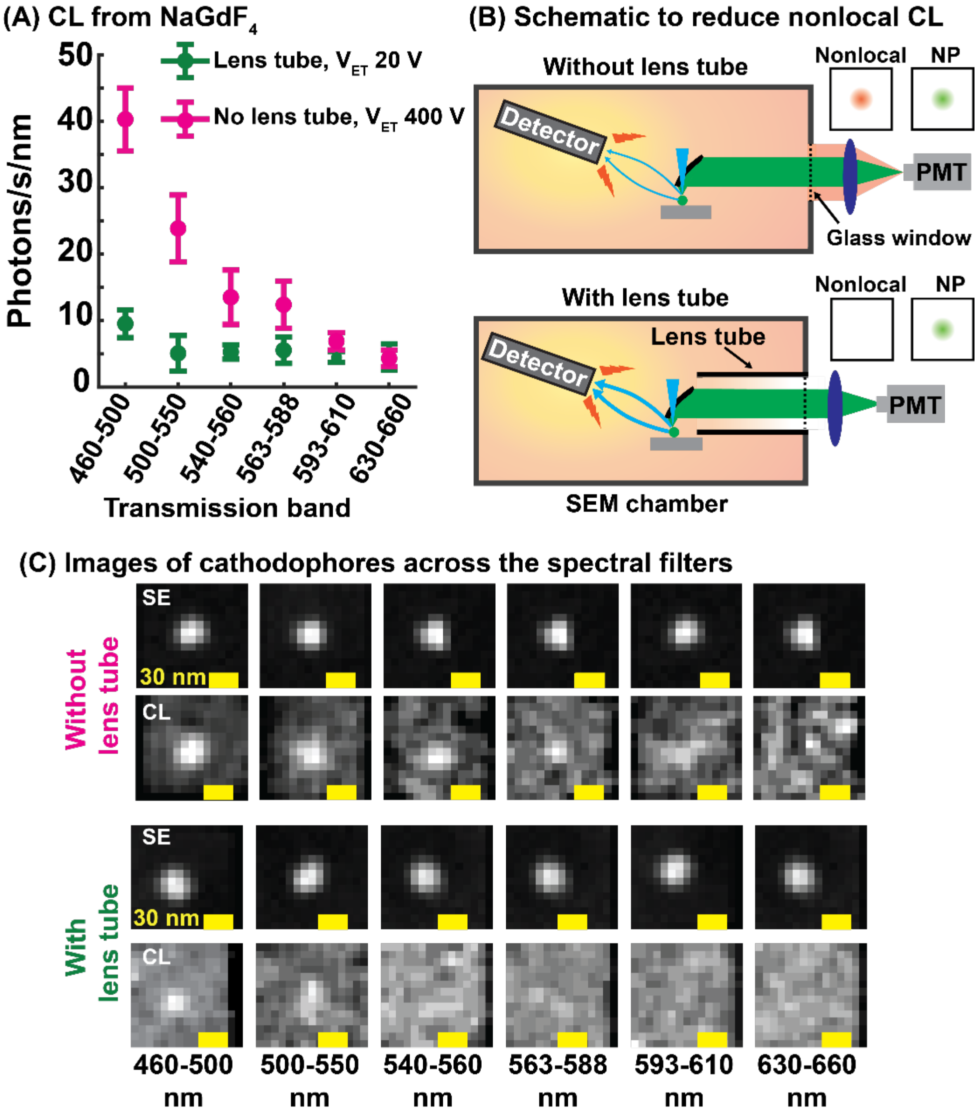

**SI Fig. 32: (A)** Rate of CL detection from non-luminescent $NaGdF_4$ nanocrystals across six spectral filters. The rate was normalized by the bandwidth of the filters. The rate is shown for two conditions: (i) without the lens tube and a voltage ($V_{ET}$) of 400 V on the Everhart-Thornley detector, and (ii) with the lens tube and a $V_{ET}$ of 20 V on the detector. The lens tube was used to prevent light emitted by the scintillator of the detector from reaching the PMTs. **(B)** A schematic showing nonlocal CL originating from the Everhart-Thornley detector upon excitation with secondary electrons. This nonlocal CL was prevented from reaching the detector by optically isolating the emission path inside the SEM with a lens tube. The tube extended from the side port of the SEM to the parabolic mirror. **(C)** SE and CL images of non-luminescent $NaGdF_4$ nanocrystals across the spectral filters shown in A, both without and with the lens tube. $V_{ET}$ was 400 V and 20 V, respectively.

with the electron beam. Deceivingly, this luminescence could be interpreted as CL from the nanocrystals because it was only observed when the electron beam was scanning across a nanocrystal, thus forming its image in the short-wavelength CL channels **(SI Fig. 32C)**. This is because the nanocrystal contained high-atomic number (high $Z$) ions ($Gd^{3+}$) while the substrate was composed of light atoms (Si), so more secondary electrons were generated when the electron beam was scanned across the nanocrystal, leading to larger secondary electron signal and increased luminescence from the scintillator in the Everhart-Thornley detector. We have addressed this issue by reducing the Everhart-Thornley bias voltage from 400 V to 20 V and optically isolating the detection path using a lens tube as shown in **SI Fig. 32B**.

#### 20. Dipole emission on different substrates

In our single-particle CL experiment, cathodophores were placed on top of a Si substrate inside the SEM vacuum chamber, and the parabolic mirror collected light emitted into vacuum by the cathodophores. To better understand the collection efficiency of the CL detection system, we used the FDTD solver in Lumerical software to perform electromagnetic simulations of an isotropic emitter placed at the Si-vacuum interface. In the absence of a Si substrate, an isotropic emitter emitted uniformly in all directions (**SI Fig. 33A**). However, the emission pattern changed when the emitter was placed on a substrate with a refractive index $n > 1$, as shown in **SI Fig. 33A**.

**SI Fig. 33B** shows fractions of energy emitted upwards, i.e., into vacuum, and into the substrate for different refractive indices ($n$) of the substrate. The emission into vacuum did not decrease monotonically as a function of $n$. Instead, the energy directed into vacuum reached its minimum at $n \sim 1.7$, and increased with a further increase in $n$. For Si ($n \sim 4$), ~47% of the total energy was emitted into vacuum. These simulations showed that for an isotropic emitter at a Si-vacuum interface, the fraction of energy directed into vacuum was not drastically different from the case in which there was no substrate (i.e., 50% for the isotropic $n = 1$ environment).

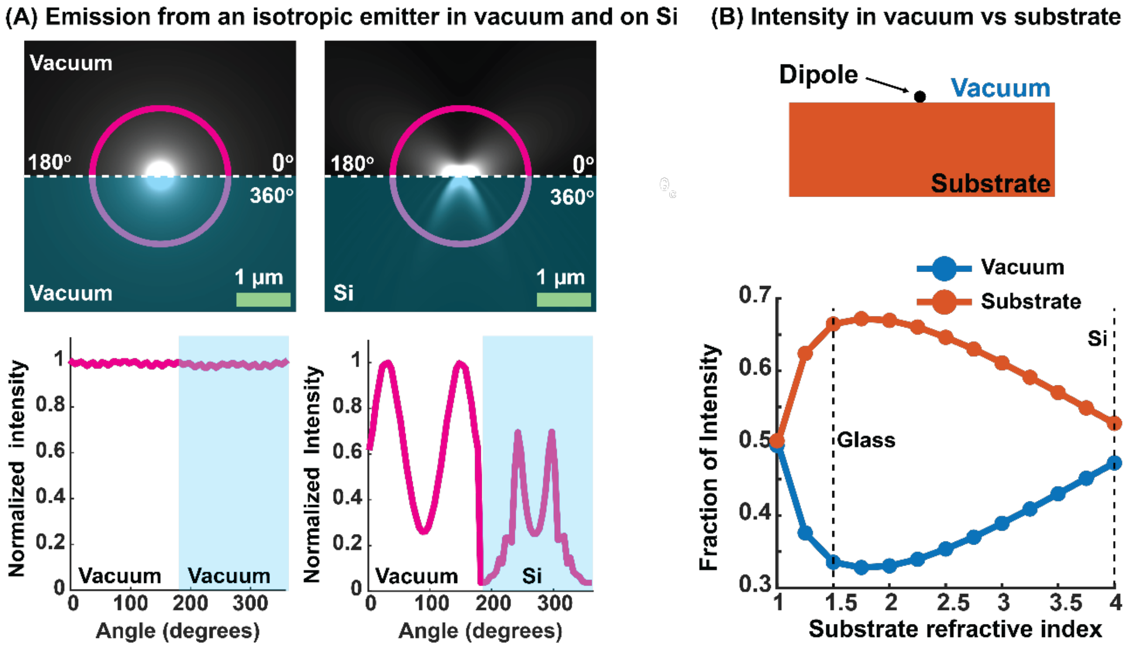

**SI Fig. 33:** **(A)** Emission profile of an isotropic emitter placed in vacuum and at the Si-vacuum interface. **(B)** Fraction of intensity emitted into vacuum and the substrate for different refractive indices of the substrate. For analysis in B, only the real part of the refractive index was considered. Simulations were performed using Lumerical FDTD.

#### 21. Repeatability

We investigated the consistency of CL signal from cathodophores across different syntheses. **SI Fig. 34** shows the rate of CL detection as a function of size for cathodophores obtained from four syntheses. The figure shows that the CL signal was consistent across the syntheses.

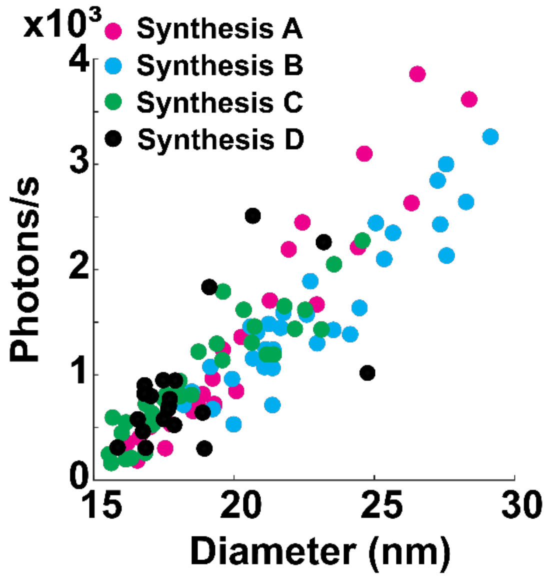

**SI Fig. 34:** CL detection rate from $NaHoF_4$ cathodophores from different syntheses. No change in the detection rate was observed. CL was measured in the $Ho^{3+}$ channel.

#### 22. Crosstalk between cathodophores in dense samples

**Simulations: Fig. 3A** shows results of Monte Carlo simulations performed to determine crosstalk between two adjacent nanocrystals, each 20 nm in diameter. In this setup, the crosstalk was minimal. However, this simulation only considered two nanocrystals. Our multicolor imaging was performed in dense monolayers of cathodophores. In such samples, the crosstalk could increase, particularly when a single cathodophore was surrounded by other cathodophores of a different color. To understand the crosstalk in such a sample, additional Monte Carlo simulations were performed using CASINO software. Simulations were performed with $NaHoF_4$ cathodophores of 20 nm diameter, and an electron beam energy of 3 keV. 1,000 trajectories were simulated for each condition.

**SI Fig. 35A** shows a situation where the excited nanocrystal (shown in green) was surrounded by eight other nanocrystals. We measured the energy deposited in the adjacent nanocrystals (shown in black) when the central nanocrystal was excited. The deposited energy was calculated at different distances, d, from the excited nanocrystal. Results are shown in **SI Fig. 35B.** The largest amount of energy was deposited into black nanocrystals for d = 0 nm, i.e., when the black nanocrystals were directly touching the green nanocrystal. In this scenario, 14.5% of energy, relative to the energy absorbed by the excited nanocrystal, was deposited in adjacent nanocrystals (~1.8% energy per adjacent nanocrystal, consistent with **Fig. 3A**). When the distance increased to 25 nm, the absorbed energy decreased to ~5%, or 0.63% per adjacent nanocrystal. We also performed a simulation with 24 nanocrystals surrounding the excited nanocrystal as shown in **SI Fig. 35C.** In this case, the energy deposited in the adjacent nanocrystals was 27.8%, or 1.16% per nanocrystal.

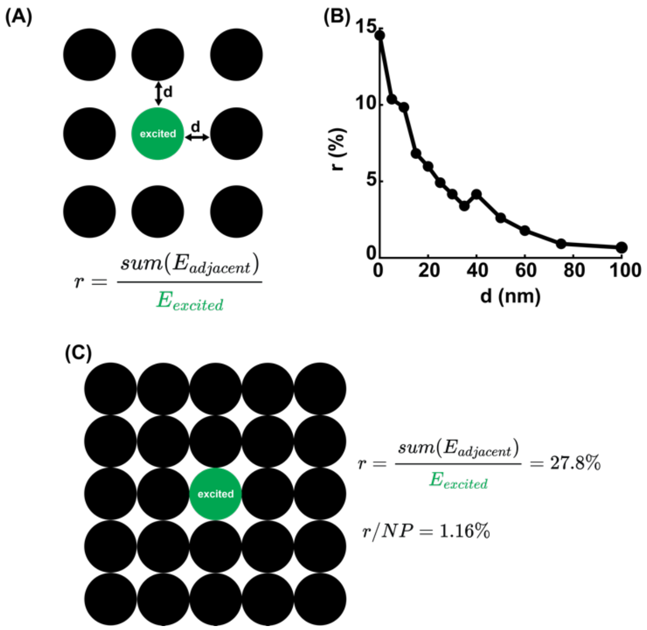

**SI Fig. 35: (A)** Monte Carlo simulations for the configuration when an excited central nanocrystal (green) was surrounded by adjacent nanocrystals (black). The nanocrystals were $NaHoF_4$, with 20 nm diameter. Electron beam energy was 3 keV and 1,000 trajectories were simulated. **(B)** Total energy (relative to energy deposited in the central nanocrystal) deposited in adjacent nanocrystals when the central nanocrystal was excited, shown as a function of the distance d between the central nanocrystal and adjacent nanocrystals. **(C)** Monte Carlo simulations for the configuration when an excited central nanocrystal was surrounded by 24 adjacent nanocrystals. 27.8% energy (relative to the central nanocrystal) was deposited in adjacent nanocrystals, corresponding to an average of 1.16% energy per nanocrystal.

These simulations show that, even in dense samples where cathodophores are in direct contact, the energy deposited in adjacent cathodophores is not prohibitive. However, despite this minimal energy absorbed per neighboring cathodophore, if a single cathodophore is surrounded by multiple cathodophores of a different color (e.g., $NaHoF_4$ surrounded by $NaDyF_4$), a detectable CL signal can appear in the wrong color channel. This potential crosstalk should be considered when classifying nanocrystals.

**Experiment:** **SI Fig. 36** shows two examples of two-color imaging when $NaHoF_4$ cathodophores were imaged adjacent to multiple $NaDyF_4$ cathodophores. In such a situation, some crosstalk was noticeable. For example, in **SI Fig. 36A** CL signal appeared in the $Dy^{3+}$ color channel when a $NaHoF_4$ cathodophore was excited (see black arrow in the cross-sectional profile). However, because the excited cathodophore absorbed considerably more energy than the surrounding nanocrystals, we could classify all the cathodophores in the FOV based on their emission in the two spectral channels.

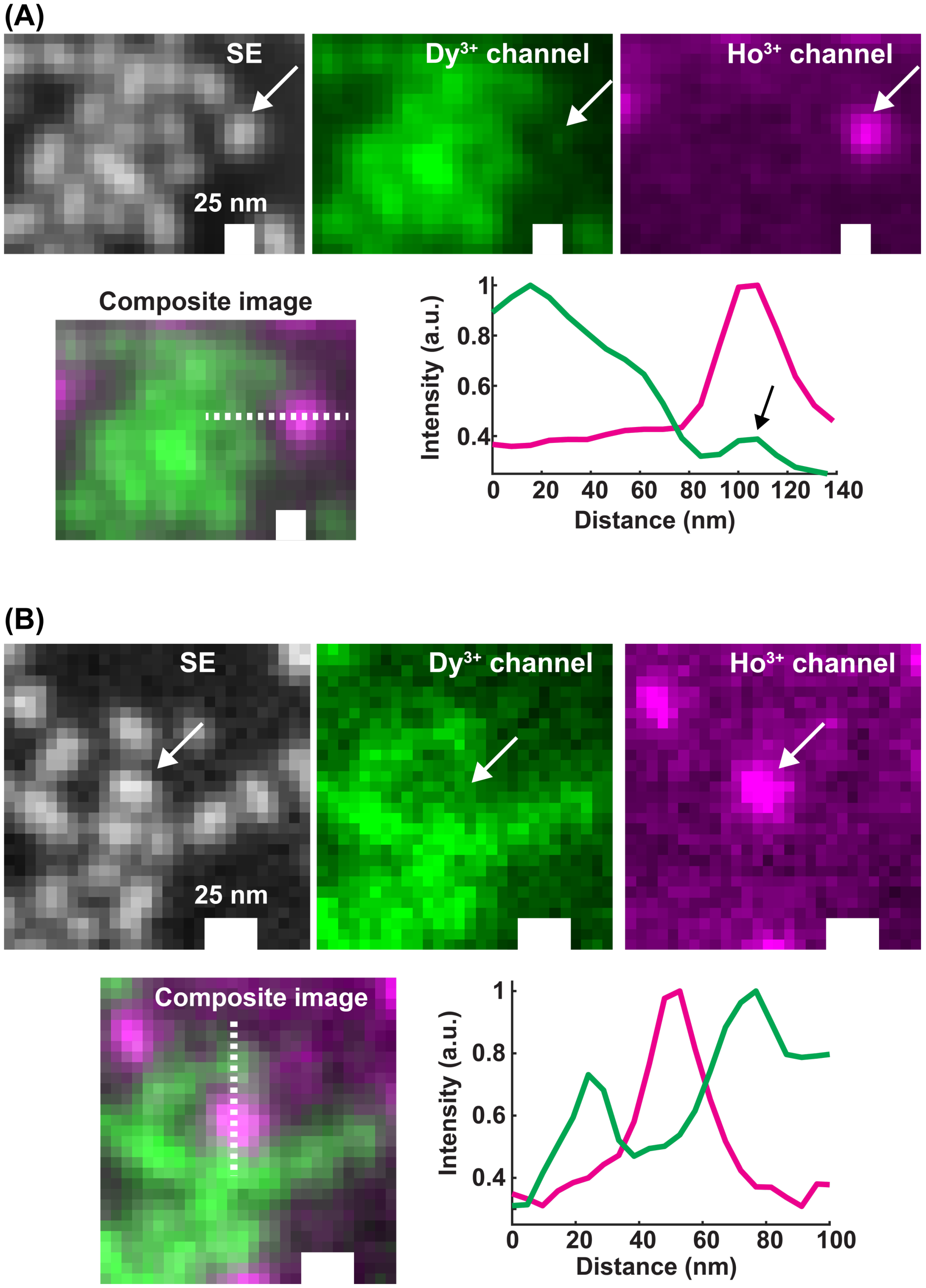

**SI Fig. 36: (A, B)** Two examples of two-color imaging in a dense, monolayered sample, when a $NaHoF_4$ cathodophore was close to multiple $NaDyF_4$ cathodophores. Crosstalk was minimal and the cathodophores could be distinguished from their CL signal in $Ho^{3+}$ and $Dy^{3+}$ color channels.
